# Protein-independent light harvesting governed by structural heterogeneity

**DOI:** 10.64898/2026.06.24.734171

**Authors:** Shun Arai, Tomomi Inagaki, Jiro Harada, Chihiro Azai, Toru Kondo

**Affiliations:** Division of Photophysical Biology, National Institute for Basic Biology, National Institutes of Natural Sciences; Okazaki, Aichi 444-8787, Japan; The Graduate University for Advanced Studies, SOKENDAI; Okazaki, Aichi 444-8787, Japan; Graduate School of Life Sciences, Ritsumeikan University; Kusatsu, Shiga 525-8577, Japan; Department of Medical Biochemistry, Kurume University School of Medicine; Kurume, Fukuoka 830-0011, Japan; School of Science and Engineering, Chuo University; Bunkyo-ku, Tokyo 112-8551, Japan; Graduate School of Science and Engineering, Chuo University; Bunkyo-ku, Tokyo 112-8551, Japan; Interconnective Photobiology Group, Exploratory Research Center on Life and Living Systems, National Institutes of Natural Sciences; Okazaki, Aichi 444-8787, Japan

## Abstract

Chlorosomes are the largest known photosynthetic light-harvesting antennas, yet unlike protein-based antennas, they lack protein scaffolds that organize pigment molecules and instead contain self-assembled tubular and lamellar bacteriochlorophyll aggregates. How these antennas achieve directional and efficient energy transfer has remained unresolved. By applying ultrafast transient absorption spectroscopy to individual wild-type and mutant chlorosomes, we resolved six kinetic components obscured by ensemble averaging and assigned each to an excitation process associated with either lamellar or tubular aggregates. Lamellar aggregates tend toward exciton localization and expand light-harvesting capacity, whereas tubular aggregates retain a relatively delocalized excitonic character and serve as the primary energy donors to the baseplate. Structural heterogeneity therefore emerges as a functional feature that balances light-harvesting capacity with robust directional energy transfer. These findings reveal a division of labor among self-assembled pigment aggregates for efficient light harvesting without protein scaffolds.

---

Photosynthetic organisms have evolved light-harvesting antenna proteins that bind pigments to capture sunlight and funnel excitation energy to reaction center (RC) protein complexes. In contrast, chlorosomes in green sulfur bacteria, the largest known light-harvesting antennas, lack protein scaffolds^1,2^. Nevertheless, they sustain highly efficient light harvesting, enabling photosynthesis even in the deep sea^3,4^. How such a protein-independent supramolecular antenna achieves efficient, directional energy transfer remains unresolved.

Chlorosomes are ellipsoidal structures (70–180 × 30–60 × 10–20 nm) enclosed by a lipid monolayer and densely packed with self-assembled aggregates of bacteriochlorophylls (BChls) *c, d*, and *e*^5^. Excitation energy absorbed by these aggregates is transferred to the baseplate, composed of CsmA dimers embedded in the lipid monolayer^6^, and then to the RC via the Fenna–Matthews– Olson (FMO) protein^7^. Within chlorosomes, pigments assemble through intermolecular interactions into lamellar aggregates and multilayered tubular aggregates (Fig. 1a)^8-11^. Studies using nuclear magnetic resonance (NMR), single-molecule spectroscopy, and theoretical calculations have provided insights into chlorosome photophysics^12-17^, but have primarily focused on aggregate-level properties. Consequently, the coordinated operation of tubular and lamellar aggregates together with the baseplate as an integrated light-harvesting system remains unclear.

**Fig. 1.**
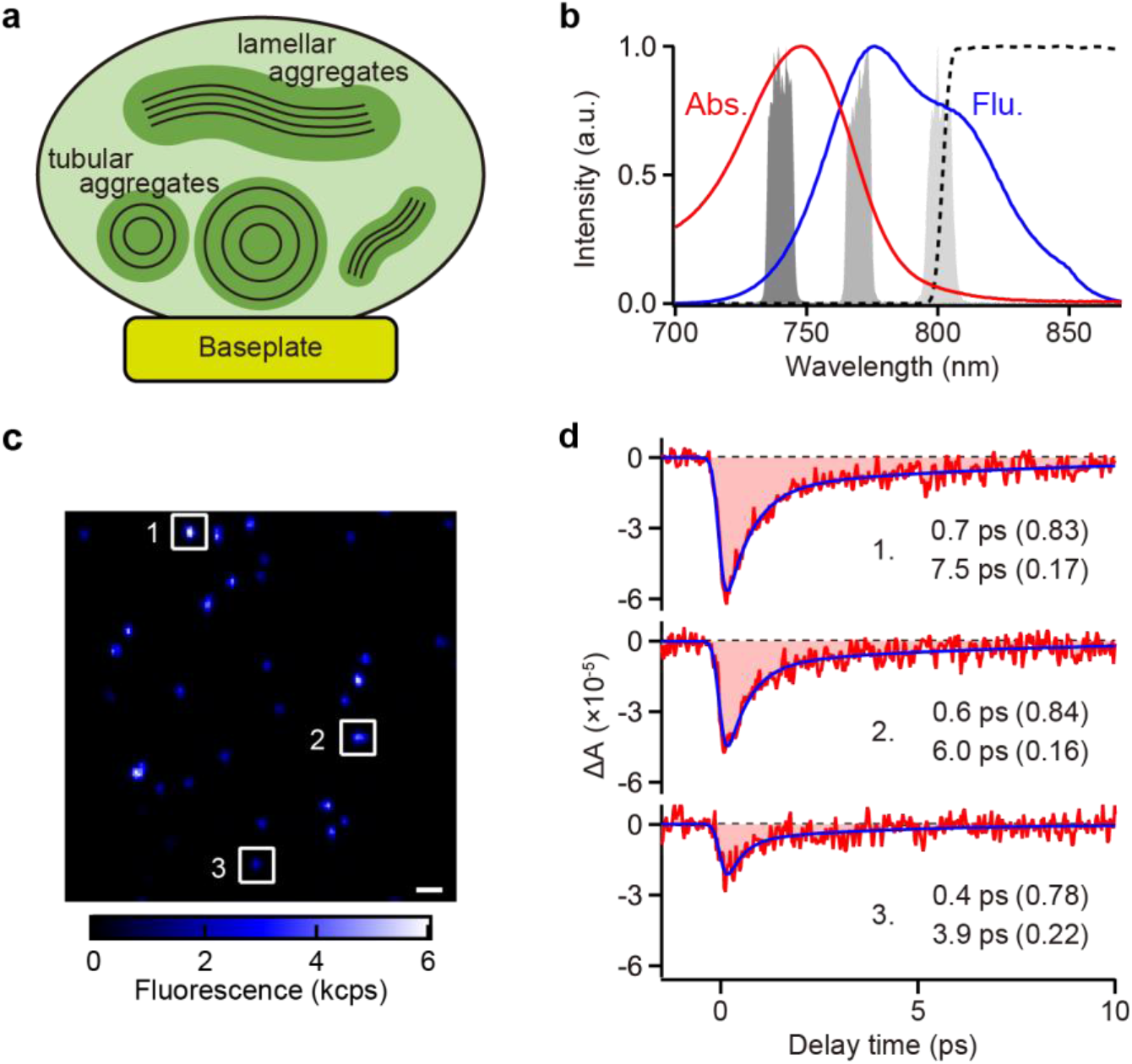
Single-chlorosome transient absorption (TA) spectroscopy. **a**, Schematic representation of a chlorosome containing tubular and lamellar aggregates and enclosed by a lipid monolayer. The baseplate is embedded in the lipid monolayer. **b**, Absorption (red) and fluorescence (blue) spectra of *tep_*WT chlorosomes. Spectral profiles of the pump beam at 740 nm (dark gray) and the probe beams at 770 nm (gray) and 800 nm (light gray), along with the fluorescence detection window at >805 nm (dashed line), are overlaid. **c**, Fluorescence image of individual chlorosomes. The scale bar represents 1 μm. **d**, TA signals of individual chlorosomes, numbered 1–3 in (c), with fitted curves (blue). The estimated time constants (*τ*) are indicated for each signal, with relative Δ*A* values shown in parentheses.

A central challenge is the structural heterogeneity of chlorosomes. The conformations and relative abundances of tubular and lamellar aggregates vary among individual chlorosomes^9,18^, and conventional ensemble spectroscopy averages over this heterogeneity, obscuring multistep photoprocesses. Resolving the mechanism therefore requires single-chlorosome measurements that connect kinetic heterogeneity with supramolecular organization.

Here, we combine single-particle spectroscopy, which resolves structural heterogeneity^14,15,19-21^, with transient absorption (TA) spectroscopy, which tracks excitation energy transfer (EET) processes^22-25^. Using this direct absorption approach^26^, rather than fluorescence-detected TA^27,28^, we resolve multiple kinetic components from heterogeneous distributions and identify their aggregate-specific photophysical signatures. These results reveal a division of labor in which lamellar aggregates expand light-harvesting capacity, whereas tubular aggregates mediate energy transfer to the baseplate while retaining a relatively delocalized excitonic character. This suggests that efficient light harvesting without protein scaffolds is achieved not by eliminating structural heterogeneity, but by tuning it to balance light-harvesting capacity and directional energy flow.

## Results

### Heterogeneous excitation dynamics of individual chlorosomes

Chlorosomes isolated from *Chlorobaculum* (*C*.) *tepidum* (*tep_*WT), which showed absorption and fluorescence maxima at 748 and 777 nm, respectively (Fig. 1b; see Supplementary Note 1), were spin-coated onto a coverslip. Fluorescence imaging under 740 nm excitation, with emission from the baseplate detected at >805 nm, resolved individual chlorosomes (Fig. 1c). TA signals, measured using a home-built TA microscope with pump and probe beams at 740 and 770 nm, respectively (see Methods and Supplementary Notes 2–5), exhibited heterogeneous decay kinetics among individual chlorosomes (Fig. 1d). Each decay trace was well fitted with a biexponential function (see Supplementary Note 6). Because an individual chlorosome contains a sub-ensemble of pigment aggregates, the observed decay kinetics likely reflect more than two components^25,29,30^, suggesting that the fitted parameters capture features of the underlying multicomponent kinetics within each chlorosome. Analysis of 194 particles yielded distributions of decay time constants (*τ*) and amplitudes (Δ*A*) for the two fitted components, as well as the total Δ*A*, defined as the sum of their amplitudes (see Supplementary Note 7). *τ* values exceeding the 10-ps experimental time window accounted for 9.0% of all fitted *τ* values and were excluded from subsequent statistical analyses because they were poorly constrained (see Supplementary Note 8). The *τ* distribution showed a distinct peak at ∼0.6 ps and a broader shoulder at ∼5 ps (Fig. 2a).

**Fig. 2.**
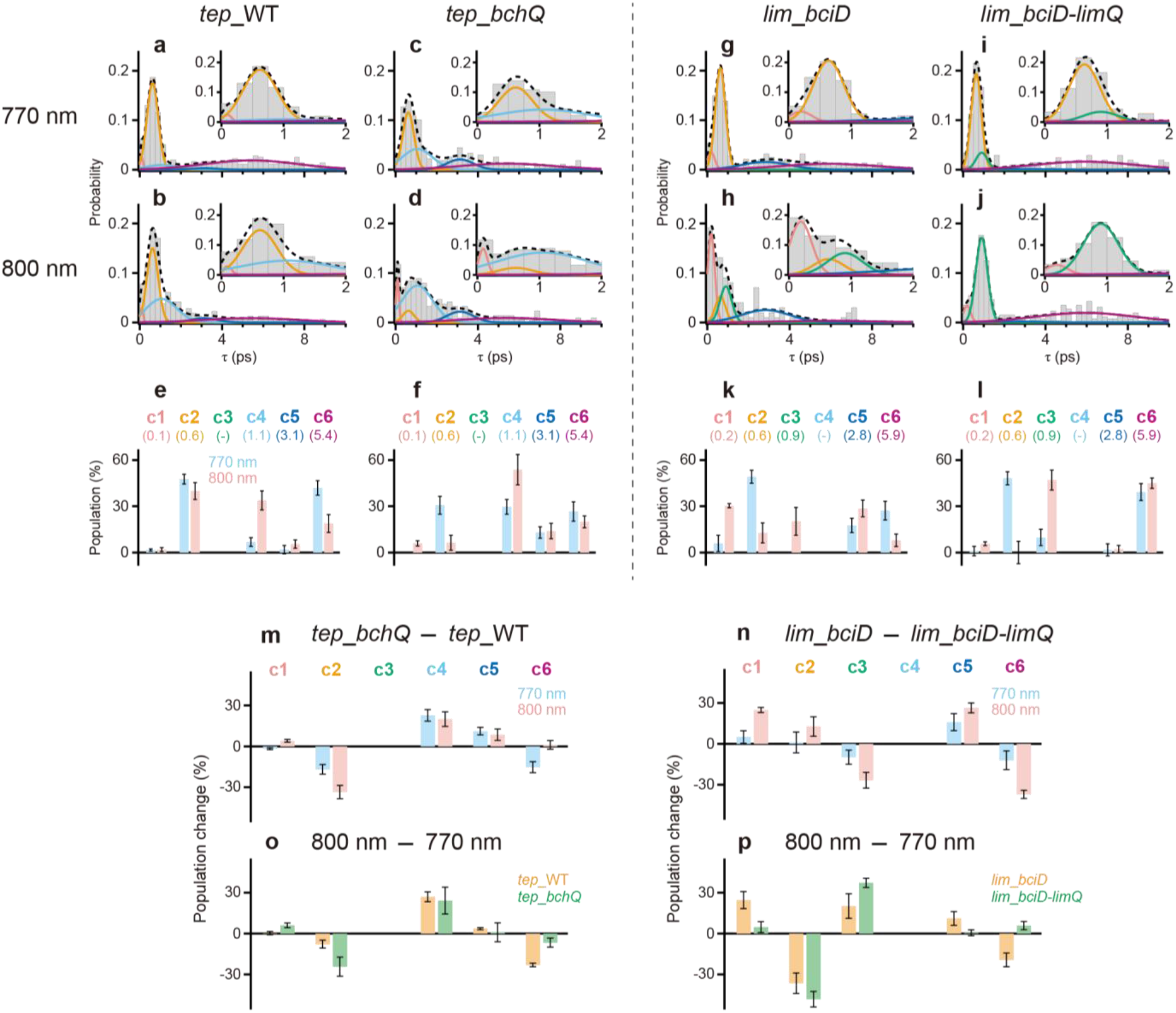
*τ*-distribution analysis. **a**–**d, g**–**j**, *τ* distributions of *tep_*WT (a, b), *tep_bchQ* (c, d), *lim_bciD* (g, h), and *lim_bciD-limQ* (i, j) chlorosomes obtained using probes at 770 nm (a, c, g, i) and 800 nm (b, d, h, j). The bin size is 0.25 ps. Expanded views of the 0–2 ps region are shown in the insets. Solid lines represent fitted curves for the *τ* components c1 (pink), c2 (orange), c3 (green), c4 (light blue), c5 (dark blue), and c6 (purple), together with their sum (dashed black line). **e, f, k, l**, *τ*-component populations in *tep_*WT (e), *tep_bchQ* (f), *lim_bciD* (k), and *lim_bciD-limQ* (l) chlorosomes at 770 nm (blue) and 800 nm (pink). *τ* values (in ps) for each component are indicated in parentheses. **m, n**, Mutation-induced changes in *τ*-component populations, calculated by subtracting the populations in *tep_*WT from those in *tep_bchQ* (m) and those in *lim_bciD-limQ* from those in *lim_bciD* (n), at 770 nm (blue) and 800 nm (pink). **o, p**, Probe-wavelength-dependent changes in *τ*-component populations, calculated by subtracting the populations at 770 nm from those at 800 nm in *tep_*WT (orange) and *tep_bchQ* (green) (o), and in *lim_bciD* (orange) and *lim_bciD-limQ* (green) (p). Error bars represent standard deviations estimated from five resampling trials of the *τ*-distribution analysis (see Supplementary Note 9). The numbers of chlorosomes included in the statistical analysis were n = 194, 148, 159 and 124 at 770 nm and n = 162, 111, 54 and 140 at 800 nm for *tep_*WT, *tep_bchQ, lim_bciD* and *lim_bciD-limQ*, respectively.

Because the long-wavelength tail of the chlorosome absorption spectrum includes the baseplate absorption band (Fig. 1b), EET to the baseplate is more selectively monitored at 800 nm than at 770 nm^23,30^. With the 800-nm probe, the *τ* distribution showed an additional peak at ∼1.5 ps and a reduced population around 5 ps (Fig. 2b). The fraction of *τ* > 10 ps decreased to 2.5%, down from 9.0% with the 770-nm probe.

### Genetic perturbations reveal aggregate-specific kinetic components

To determine which kinetic components are associated with lamellar or tubular aggregates, we analyzed chlorosomes from *tep_bchQ*, a *C. tepidum* mutant lacking BchQ, an enzyme that modifies the alkyl group at the C8^2^ position of BChl *c*^31^. This mutation shortens the C8^2^ side chains (Fig. S1), reducing steric hindrance and thereby promoting the assembly of tubular aggregates (see Supplementary Note 1)^9^. The *tep_bchQ* chlorosomes exhibited marked changes in the *τ* distribution. With the 770-nm probe (Fig. 2c), peaks appeared at ∼1 and ∼3 ps, and the population at ∼5 ps decreased relative to *tep_*WT (Fig. 2a). These features were more pronounced with the 800-nm probe (Fig. 2d), with a reduced population at ∼0.6 ps and the emergence of a distinct peak at <0.6 ps. Global fitting of the four *τ* distributions from *tep_*WT and *tep_bchQ* at 770 and 800 nm resolved five Gaussian components (Fig. 2a–d, solid lines; see Supplementary Note 9). The fitted peak centers, with full widths at half maximum (FWHMs) in parentheses, were 0.1 (0.2), 0.6 (0.7), 1.1 (1.7), 3.1 (1.5), and 5.4 (5.3) ps, with distinct populations as shown in Fig. 2e, f (see also Table S5a).

As a complementary perturbation, overexpression of the enzyme BchQ in *C. limnaeum* extends the C8^2^ side chain of BChl relative to that in the wild type^32^. Although *C. limnaeum* normally synthesizes BChl *e*, loss of the enzyme BciD induces BChl *c* biosynthesis, as in *C. tepidum*^33^. Thus, chlorosomes from a *C. limnaeum* mutant that both overexpresses BchQ and lacks BciD (*lim_bciD-limQ*) contain BChl *c* with elongated side chains (Fig. S1), likely promoting lamellar aggregate formation (see Supplementary Note 1). Comparison of this double mutant with the *lim_bciD* single mutant, which lacks only BciD and served as a reference, therefore enabled the assignment of components associated with lamellar aggregates.

In *lim_bciD*, the *τ* distribution exhibited a prominent peak at ∼0.6 ps and a broad peak around 3 ps when probed at 770 nm (Fig. 2g). At 800 nm, populations at ∼0.2 and ∼1 ps were enhanced (Fig. 2h). By contrast, chlorosomes from *lim_bciD-limQ* lacked the ∼3 ps peak at 770 nm and showed an increased population around 5 ps (Fig. 2i). At 800 nm, the population at ∼1 ps became more prominent (Fig. 2j). These four distributions were reproduced by global fitting with five Gaussian components, with peak centers (FWHMs) of 0.2 (0.4), 0.6 (0.6), 0.9 (0.7), 2.8 (2.8), and 5.9 (5.6) ps. These *τ* components closely matched those obtained from *tep_*WT and *tep_bchQ*, while their populations varied substantially (Fig. 2k, l; see also Table S5b).

These variations likely arose from differences in aggregate composition due to the C8^2^ side-chain length of BChl *c*^31,32,34^, which shifts the balance between tubular and lamellar aggregates. To isolate the effect of side-chain shortening, we subtracted the population of each *τ* component in the longer-side-chain sample from that in the shorter-side-chain sample, i.e., *tep_bchQ* minus *tep_*WT (Fig. 2m) and *lim_bciD* minus *lim_bciD-limQ* (Fig. 2n). The 0.1–0.2 ps components (c1) increased with side-chain shortening, indicating their origin in tubular aggregates. In contrast, the 0.6 ps components (c2) decreased and were assigned to lamellar aggregates. The 0.9 ps component (c3), observed in *lim_bciD* and *lim_bciD-limQ*, decreased, whereas the 1.1 ps component (c4), found in *tep_bchQ* and *tep_*WT, increased, suggesting that c3 and c4 arise from lamellar and tubular aggregates, respectively. Increases in the 2.8–3.1 ps components (c5) and decreases in the 5.4–5.9 ps components (c6) indicated their assignments to tubular and lamellar aggregates, respectively. Together, these genetic comparisons assigned c1, c4, and c5 to tubular aggregates and c2, c3, and c6 to lamellar aggregates.

Because EET to the baseplate is more selectively monitored at 800 nm, we identified baseplate-associated components by subtracting the *τ* populations at 770 nm from those at 800 nm in *C. tepidum* (Fig. 2o) and *C. limnaeum* (Fig. 2p). Components c3 and c4, attributed to lamellar and tubular aggregates, respectively, exhibited marked increases, suggesting that they reflect EET to the baseplate via the respective aggregates.

### Photophysical signatures of distinct kinetic components

We quantified the time constants (*τ*), absorbance changes (Δ*A*), and fluorescence intensity reflecting the amount of excitation energy reaching the baseplate, allowing estimation of three additional parameters: the total Δ*A*, defined as the sum of Δ*A* for the two decay components; the relative Δ*A*, defined as the ratio of Δ*A* for each decay component to the total Δ*A*; and the effective EET efficiency to the baseplate, defined as the ratio of fluorescence intensity to the total Δ*A* (see Supplementary Note 7). For *tep_*WT chlorosomes probed at 800 nm, we constructed two-dimensional (2D) distributions of *τ* versus these parameters. Fitting the 2D distributions with five 2D Gaussian functions (see Supplementary Note 10) yielded estimates of these parameters and their distribution widths for the five *τ* components (Fig. 3a–c), excluding c3, which was not observed in *tep_*WT chlorosomes (Fig. 2e). Similarly, in *tep_bchQ* chlorosomes, the relative Δ*A* for each *τ* component was estimated (Fig. 3d). All 2D distributions for wild-type and mutant chlorosomes at 770 and 800 nm and the corresponding fitted parameter distributions are shown in Figs. S20–S22 and Figs. S2–S4, respectively.

**Fig. 3.**
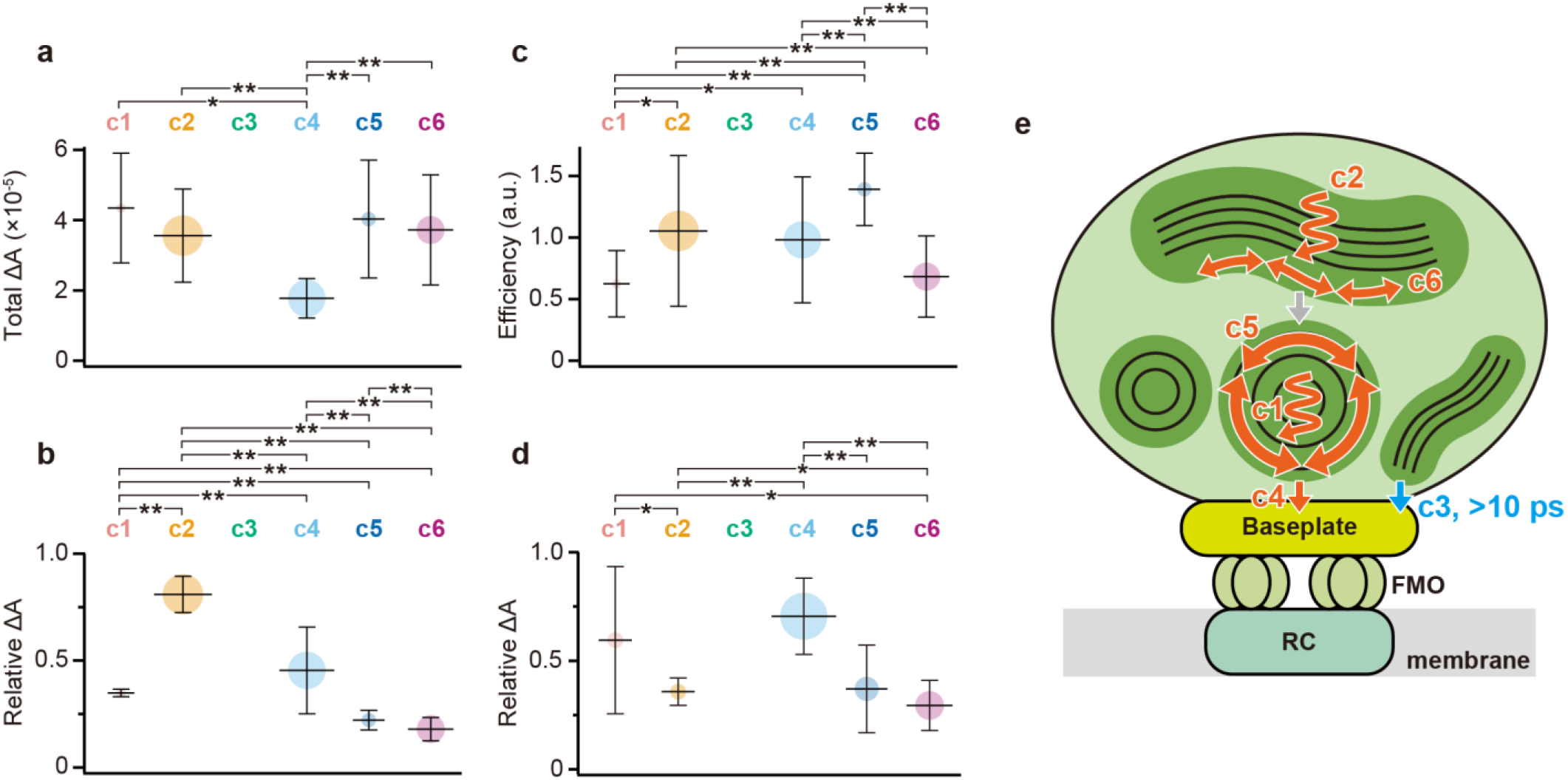
2D distribution analysis. **a**–**d**, Distributions of total Δ*A* (a), relative Δ*A* (b), and effective EET efficiency (c) for each *τ* component in *tep_*WT chlorosomes (n = 162), and the corresponding distribution of relative Δ*A* for each *τ* component in *tep_bchQ* chlorosomes (n = 111) (d), obtained using the 800-nm probe. Each component is colored as in Fig. 2. The mean values are indicated by horizontal bars. The 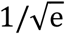. full widths are indicated by vertical bars. The area of each circle represents the population of the corresponding *τ* component. Statistically significant differences assessed using Welch’s *t*-test between components are denoted by asterisks (\**p* < 0.05 and \*\**p* < 0.01) (see Supplementary Note 10). **e**, Schematic illustration of excitation dynamics within chlorosomal aggregates and toward the baseplate. Tubular and lamellar aggregates are represented by black circles and black wavy lines, respectively. The orange arrows represent excitation processes assigned to each *τ* component. The gray arrow indicates EET between tubular and lamellar aggregates, which was not identified in the present measurements. The c3 and >10-ps components, which were minor in the wild type, are indicated in light blue.

## Discussion

Based on mutant analysis, components c1, c4, and c5 were assigned to tubular aggregates, whereas c2, c3, and c6 were attributed to lamellar aggregates (Fig. 2m, n). Previous studies employing hole-burning, single-molecule, and TA spectroscopies have suggested multiple spectrally distinct domains within chlorosomes^15,19,25^. TA signals in the long-wavelength region (770 and 800 nm) following 740 nm excitation reflect excitation dynamics involving lower-energy excited states and energy transfer to the baseplate^22,25^. The shortest time constants, c1 (0.1–0.2 ps) and c2 (0.6 ps), correspond to sub-ps components previously reported by TA spectroscopy, 2D electronic spectroscopy, and theoretical simulations^22,23,29,35-37^, and were therefore assigned to ultrafast relaxation processes occurring immediately after photoexcitation within tubular and lamellar aggregates, respectively.

Reported time constants for EET to the baseplate span a wide range, including ∼3 ps^22^, ∼7–12 ps^23^, 30–40 ps^25^, and >100 ps^23^. Quantum chemical calculations have also estimated values from a few to several tens of ps^29,36^. Furthermore, time constants around 1 ps have been discussed in the context of intra-baseplate EET^23,24^. By combining single-chlorosome TA at 770 and 800 nm with mutant analysis, c3 (0.9 ps) and c4 (1.1 ps) were assigned to baseplate-directed EET from lamellar and tubular aggregates, respectively. In *tep_*WT chlorosomes, where c3 was not detected, c4 showed greater saturation under intense excitation than the longer-*τ* components, supporting its assignment (see Supplementary Note 5). Although components with *τ* > 10 ps were also observed, as previously reported, their fraction decreased from 9.0% at 770 nm to 2.5% at 800 nm. This fraction increased with elongation of the BChl side chain (Fig. S29d). These components may therefore reflect EET from lamellar aggregates to the baseplate via higher-energy exciton states, representing minor subsidiary pathways (see Supplementary Note 11). The TA signal from baseplate BChl *a* was likely obscured by its low abundance (∼1% of total BChl)^38^.

The remaining components, c5 (2.8–3.1 ps) and c6 (5.4–5.9 ps), were attributed to intra- or inter-aggregate EET within their respective aggregate types, occurring after the initial excitation relaxation, consistent with previously reported *τ* values of ∼1.7 and ∼5.8 ps^25^. Specifically, c5 originated from tubular aggregates, whereas c6 arose from lamellar aggregates. EET between tubular and lamellar aggregates may also occur, but is likely confined to their interfacial regions, indicating a minor contribution. In summary, we identified excitation relaxation, intra- or inter-aggregate EET within each aggregate type, and baseplate-directed EET, and linked each component to tubular or lamellar aggregates, as summarized in Fig. 3e.

In *tep_*WT chlorosomes, after excitation relaxation (c1, c2), EET occurred in both tubular (c5) and lamellar (c6) aggregates (Fig. 3e). EET to the baseplate originated predominantly from tubular aggregates (c4), with no detectable c3 contribution from lamellar aggregates. The 2D distribution analysis of *τ* versus effective EET efficiency to the baseplate (Fig. 3c) showed higher effective EET efficiency in chlorosomes with the tubular components c4 or c5 than in those with the lamellar component c6, with *p* < 2 × 10^−5^. A similar trend was observed in *tep_bchQ* chlorosomes (Fig. S4f). These results indicate that tubular aggregates function as the primary donors for EET to the baseplate. Given that c3 was absent in *tep_*WT (Fig. 2e) but prominent in *lim_bciD-limQ* chlorosomes (Fig. 2l), EET from lamellar aggregates to the baseplate may also occur, but appears to be minor in the wild type.

Within aggregates, intermolecular interactions cause mixing of molecular excited states, leading to delocalized electronic states spanning multiple pigments with collective transition dipole moments. The absorbance change, Δ*A*, reflects the magnitude of the dipole moment of excitonic domains involved in excitation dynamics^39^, and thus can serve as an indicator of antenna size. A 2D distribution analysis of *τ* versus total Δ*A* (Fig. 3a) revealed that chlorosomes exhibiting c4 had relatively small total Δ*A*, whereas those with c5 or c6 had significantly larger values (*p* < 7 × 10^−5^ compared to c4), suggesting that chlorosomes with larger effective antenna sizes contain greater amounts of both tubular and lamellar aggregates.

With the 770-nm probe, lamellar-associated components (c2, c6) predominated (Fig. 2e, blue), suggesting that lamellar aggregates preferentially contribute to the shorter-wavelength TA response and transfer excitation energy to energetically downstream tubular aggregates. This dominance of lamellar components at 770 nm was broadly observed across wild-type and mutant chlorosomes (Fig. 2e, f, k, and l, blue). Additionally, 2D distribution analyses of *τ* versus total Δ*A* at 770 nm (Fig. S2a–d) showed that lamellar components in both wild-type and mutant chlorosomes tended to exhibit broader distributions of total Δ*A*. Thus, variation in lamellar aggregate content is likely to modulate light-harvesting capacity. This division-of-labor model (Fig. 3e) is also consistent with the photobleaching behavior (Fig. S5; see Supplementary Note 3) and excitation-fluence dependence (Fig. S6; see Supplementary Note 5).

The functional differences between the aggregate types may arise from distinct excitonic dynamics. When the lamellar components c2 and c6 were compared in the 2D distribution of *τ* versus relative Δ*A* (Fig. 3b), the relative Δ*A* of the EET component c6 was only 22% of that of the excitation-relaxation component c2 (*p* < 10^−10^). A similar decrease was observed at 770 nm (Fig. S3a–d). Given that Δ*A* reflects the magnitude of the dipole moment of each excitonic domain involved in the process^39^, this decrease suggests that excitation is initially delocalized over excitonic domains with large dipole moments and then relaxes into domains with smaller dipole moments. This behavior is consistent with exciton localization onto finite molecular segments within aggregates, from which EET proceeds^25,35,40^. Such localization is thought to arise from energetic disorder^41^, likely caused by structural and environmental heterogeneities. The previously estimated exciton size of about 2–3 pigments may correspond to localized domains^39,42^.

In contrast, for tubular aggregates, the relative Δ*A* of the EET component c5 was as high as 64% of that of the excitation-relaxation component c1 (Fig. 3b). This trend was also observed in *tep_bchQ* chlorosomes enriched in tubular aggregates (Fig. 3d), where the relative Δ*A* of c5 was 62% of that of c1 (*p* = 0.05). Therefore, the initially formed exciton states appear to remain relatively delocalized before EET. Tubular aggregates have a helical cylindrical structure^8,12^, producing two higher-energy exciton states with dipole moments perpendicular to the cylinder axis and one lower-energy state with a dipole moment parallel to it^43,44^. Moreover, as the cylinder diameter increases, the tubular excitonic states tend to shift to lower energy^15,36,45^, suggesting that in multilayered tubular structures, EET proceeds directionally from inner to outer tubes. Accordingly, after photoexcitation, tubular aggregates relax from higher to lower exciton states with dipole reorientation, possibly underlying the previously observed transient absorption anisotropy^23,24^. This relaxation may involve directional energy flow from inner to outer tubes while retaining excitonic delocalization. Finally, excitation energy is transferred to the baseplate via lower-energy exciton states, such as those probed at 800 nm.

Exciton delocalization is highly sensitive to structural disorder^41,44,46^. For instance, in *tep_bchQ* chlorosomes, where side-chain shortening at the C8^2^ position of BChl *c* leads to more uniform molecular packing, even lamellar aggregates appear to retain delocalized exciton states, as indicated by the relative Δ*A* of c6 being close to that of c2 (Fig. 3d). By contrast, in *tep_*WT chlorosomes with higher structural disorder, tubular aggregates predominantly retain delocalized exciton states, likely due to their helical cylindrical architecture. In such structures, the reduced density of states (DOS) in the lower-energy region of the exciton band suppresses exciton scattering that causes localization^46^. The suppression intensifies with increasing cylinder diameter^44^. Consequently, the tubular system can retain a large dipole moment even after relaxation, likely promoting efficient EET to the baseplate.

Because aggregate-specific EET is difficult to resolve experimentally in structurally complex chlorosomes, mechanistic interpretations have relied substantially on theoretical models^29,36^. Here, based on our single-chlorosome TA analysis, we propose the model shown in Fig. 3e (see also Supplementary Note 12), in which lamellar and tubular aggregates play distinct functional roles as modulators of light-harvesting capacity and mediators of EET to the baseplate, respectively. Adjusting their relative abundances may enable flexible adaptation to varying light conditions.

This balance may be governed by intermolecular packing interactions between BChls^47^, which can be altered by steric hindrance caused by methylation at the C8^2^ and C12^1^ positions^31^. Under low-light conditions, green sulfur bacteria increase chlorosomal BChl *c* content and broaden the Q_y_ absorption peak, whereas mutants with suppressed methylation lack these responses and exhibit smaller chlorosomes and slower growth^31^. Accordingly, methylation likely promotes lamellar aggregate formation, increasing antenna size and thereby improving growth in dim light. A key factor in this adaptive mechanism is structural heterogeneity, a crucial modulator of excitonic properties. Importantly, heterogeneity is not simply minimized or maximized, but rather appears to be tuned to maintain a functional balance (see also Supplementary Note 13). This principle may guide the design of artificial photosynthetic systems based on pigment aggregates without protein scaffolds.

## Methods

### Preparation of wild-type and mutant chlorosomes

Chlorosomes containing BChl *c* aggregates were isolated from green sulfur bacterial cells of the wild-type and *bchQ* mutant strains of *C. tepidum*, referred to as *tep_*WT and *tep_bchQ*, respectively, and of the *bciD* and *bciD*-*limQ* mutant strains of *C. limnaeum*, referred to as *lim_bciD* and *lim_bciD-limQ*, respectively^32^. Isolation was performed according to previously reported methods^48,49^, with certain modifications, as outlined below.

The green sulfur bacterial cells were cultivated in approximately 5 L of CL medium^50^ supplemented with the appropriate antibiotics until they reached the late-exponential growth phase in batch culture. The cells were harvested by centrifugation at 10,000×*g*, yielding approximately 10 g of wet cell pellet. After the cells were completely resuspended in 200 mL of SCN buffer (10 mM sodium phosphate, 2 M sodium thiocyanate, pH 7.0) by homogenization, the cells were disrupted by sonication (SFX-250, BRANSON). Following removal of unbroken cells and debris by centrifugation at 10,000×*g* for 20 min, the supernatant was subjected to ultracentrifugation at 150,000×*g* for 2 h at 4 °C. The resulting pellet containing the chlorosomes was resuspended in 15 mL of the SCN buffer. The chlorosomes, after detachment from the cytoplasmic membrane, were then separated on a discontinuous sucrose density gradient (15–40% (w/v) in 5% steps) prepared in the SCN buffer by ultracentrifugation (150,000×*g*, 12 h, 4 °C). The deep green band around 20% sucrose was collected and dialyzed against 20 mM sodium phosphate buffer (pH 7.0) at 4 °C to remove sucrose and thiocyanate from the solution. After the purity was confirmed by steady-state absorption spectroscopy, the resulting chlorosome preparation was frozen and stored at – 80 °C until use.

### Modification of the alkyl substituent at the C8^2^ position of BChl *c*

The C8^2^ side-chain composition of BChl *c* was modulated by genetic modifications affecting side-chain methylation. The C8^2^ alkyl substituents of BChl *c* include ethyl, propyl, isobutyl, and neopentyl groups, whose relative abundances are regulated by pigment biosynthetic methyltransferases^31,32^. The compositions of these substituents in the wild-type and mutant chlorosomes used in this study were taken from previous reports^31,32,34^ and are summarized in Fig. S1a. Because progressive methylation at the C8^2^ position lengthens the alkyl side chain in the order ethyl < propyl < isobutyl < neopentyl, the effective C8^2^ side-chain length increases in the order *tep_bchQ* < *tep_*WT < *lim_bciD* < *lim_bciD-limQ*. A longer C8^2^ side chain is expected to increase intermolecular steric hindrance during supramolecular self-assembly and thereby favor the formation of lamellar aggregates, whereas a shorter C8^2^ side chain tends to promote the formation of tubular aggregates^9^ (see Supplementary Note 1).

### Transient absorption microscopy

Details of our experimental setup have been described previously^26^. Briefly, a Ti:sapphire laser (Synergy, Spectra-Physics) emitted pulses at a repetition rate of 78 MHz, with a pulse duration of ∼14 fs, a central wavelength of ∼790 nm, and a spectral width of ∼120 nm (FWHM). The laser output was divided into pump and probe beams by a beam splitter. A quarter-wave plate (WPM03-Q-NIR, Newlight Photonics) was used to reduce the degree of linear polarization, defined as (*I*_max_ −*I*_min_)/(*I*_max_ +*I*_min_), to 0.09 for the pump beam and 0.25 for the probe beam. Here, *I*_max_ and *I*_min_ denote, respectively, the maximum and minimum intensities of light transmitted through a linear polarizer. This near-circular polarization suppresses the orientation dependence of the sample response.

The pump beam was spectrally narrowed to a central wavelength of 740 nm with a bandwidth of ∼10 nm using a 750-nm band-pass filter (FBH750-10, Thorlabs) tilted by ∼20° with respect to the optical axis (Fig. 1b, dark gray). The probe beam was spectrally selected using a 770-nm band-pass filter (FBH770-10, Thorlabs) or an 800-nm band-pass filter (FBH800-10, Thorlabs) (Fig. 1b, gray and light gray, respectively), and then directed to an optical delay line (scanDelay USB-15, APE) to scan the time interval between pump and probe pulses. The pump and probe beams were combined in a parallel geometry using a beam splitter and then divided by another beam splitter into signal and reference beams for balanced detection. The reference beam was focused onto the reference detection channel of a balanced photodiode (Nirvana, Newport) after passing through two long-pass filters (FELH0750, Thorlabs) to remove pump light. The signal beam, after passing through an oil-immersion objective with a numerical aperture (NA) of 1.45 (UPLXAPO100XO, Olympus), was reflected by a mirror positioned a few μm above the sample and subsequently focused onto the sample plane, as illustrated in Fig. S7. Light transmitted through the sample was collected by the same objective, passed through the beam splitter, and then focused onto the signal detection channel of the balanced photodiode after passing through two long-pass filters (FELH0750, Thorlabs) to remove pump light.

The pump beam was modulated at 45 kHz by an optical chopper (310CD, Scitec Instruments). The differential output from the balanced photodiode was demodulated at the chopper frequency using a lock-in amplifier (LI5640, NF Corporation). The pump–probe delay was scanned over a 14.4-ps range, including a 10-ps time window to record transient decays, at a rate of 4 Hz. Signal acquisition was performed using a digitizer board (DIG-100M1002-PCI, CONTEC) with a sampling rate of 100 kHz, corresponding to an effective sampling interval of 1 fs in delay time. The data points were averaged in 30-fs bins to obtain the TA decay trace.

Fluorescence was reflected by the mirror placed above the sample and collimated by the same objective. It was then reflected by the two beam splitters, transmitted through an edge-pass filter used as a dichroic mirror (FELH0750, Thorlabs) and a long-pass filter (FF01-793/LP-25, Semrock), focused into a multimode fiber, and finally detected by an avalanche photodiode (SPCM-AQRH-14-FC, Excelitas). The sample position was raster-scanned using a piezo stage (Nano-LP100, Mad City Labs) to acquire fluorescence images.

The spatial resolution of our instrument, defined as the FWHM of the focal spot along the *x, y*, and *z* axes, was estimated to be 336/243/1145 nm (*x*/*y*/*z*) for absorption microscopy and 309/257/1031 nm (*x*/*y*/*z*) for fluorescence microscopy^26^.

Conventional absorption microscopes typically use a two-objective configuration, in which one of the opposing objectives focuses the incident beam onto the sample and the other collects the transmitted beam^51,52^. In our microscope, the mirror placed above the sample mounted on a coverslip enables both focusing and collection through a single objective, eliminating misalignment and mechanical drift associated with two-objective configurations. The gap between the coverslip carrying the sample and the mirror surface was approximately 3–8 μm and filled with immersion oil to ensure refractive-index matching (Fig. S7). The detection limit for TA signals, defined as the Δ*A* corresponding to a signal-to-noise ratio (SNR) of 1, was experimentally determined to be on the order of 10^−6^ with a signal-beam power of 4 μW (corresponding to ∼14 μW before the objective), a lock-in amplifier time constant of 100 μs, and a probe wavelength of 770 nm^26^.

### Single-chlorosome measurements

Chlorosome solutions were diluted to an optical density (OD) of ≈ 2 at the Q_y_ absorption band in 40 mM Tris–HCl (pH 8.0) buffer. Because the redox state substantially affects the photophysical properties of chlorosomes, 10 mM sodium dithionite was added to maintain reducing conditions (see Supplementary Note 2). To remove dissolved oxygen, an enzymatic oxygen-scavenging system consisting of 1 U mL^−1^ protocatechuate 3,4-dioxygenase (PCD) and 2.5 mM protocatechuic acid (PCA) was added. After the addition of 1% (w/v) polyvinyl alcohol (PVA), the sample solution was spin-coated onto a coverslip at 3000 rpm for 1 min. A 1-μL aliquot of immersion oil was spin-coated onto a silver mirror (PF03-03-P01, Thorlabs) at 6000 rpm for 1 min to form a thin oil layer. The oil-coated mirror was then placed over the sample. The assembly was sealed by placing an O-ring and capping it with an additional coverslip to prevent oxygen ingress. The sample was left at room temperature in the dark for at least one night to allow further oxygen removal (see Supplementary Note 3). The PVA and oil coatings did not substantially affect either the fluorescence peak of the BChl aggregates or the apparent EET efficiency from the BChl aggregates to the baseplate, suggesting that the coating procedure had only a minimal effect on the BChl aggregates and on EET from the aggregates to the baseplate (see Supplementary Note 4).

The pump photon fluence was set to 1.5 × 10^14^ photons pulse^−1^ cm^−2^. At higher pump fluences, the TA response changed (see Supplementary Note 5). The probe photon fluences were 9.3 × 10^14^ photons pulse^−1^ cm^−2^ at 770 nm and 1.3 × 10^15^ photons pulse^−1^ cm^−2^ at 800 nm. TA signals acquired over ∼0.5–2 min were averaged to improve the SNR.

Fluorescence imaging was performed using 740-nm excitation (Fig. 1b, dark gray) with fluorescence detected at >805 nm (Fig. 1b, dashed line). The excitation photon fluence was 1.7 × 10^13^ photons pulse^−1^ cm^−2^. The raster scan was performed with 0.1-μm steps and a 20-ms integration time per step. For each particle, the fluorescence intensity was calculated as the mean intensity within a 5 × 5 pixel region centered on the TA measurement position.

### Data analysis and statistics

TA traces from individual chlorosomes were fitted with a biexponential function convolved with a Gaussian instrument response function (IRF). An SNR cutoff of 3 was applied, and the fitting results obtained from traces with an SNR > 3 were retained for further statistical analyses. Time constants exceeding the 10-ps measurement window were excluded from the main *τ*-distribution analysis because they were poorly constrained. For each of the two strain pairs (*tep_*WT/*tep_bchQ* and *lim_bciD*/*lim_bciD-limQ*), the four *τ* distributions obtained with probes at 770 and 800 nm were globally fitted with five Gaussian components. 2D distributions of *τ* versus total Δ*A*, relative Δ*A*, or effective EET efficiency were analyzed using constrained 2D Gaussian fitting. Pairwise differences between the center positions of the fitted parameter distributions were assessed using Welch’s *t*-test, with effective sample sizes estimated from the component populations. Further details of the fitting procedures, evaluation of the number of components, uncertainty estimation, and robustness analyses are provided in Supplementary Notes 6–10.

## Acknowledgements

The authors thank Yu Takaba at Chuo University and Kyoka Tani and Kazuki Terauchi at Ritsumeikan University for technical support with sample preparation and Yutaka Shibata at Tohoku University for helpful discussions.

