## Supplementary Materials for "Protein-independent light harvesting governed by structural heterogeneity"

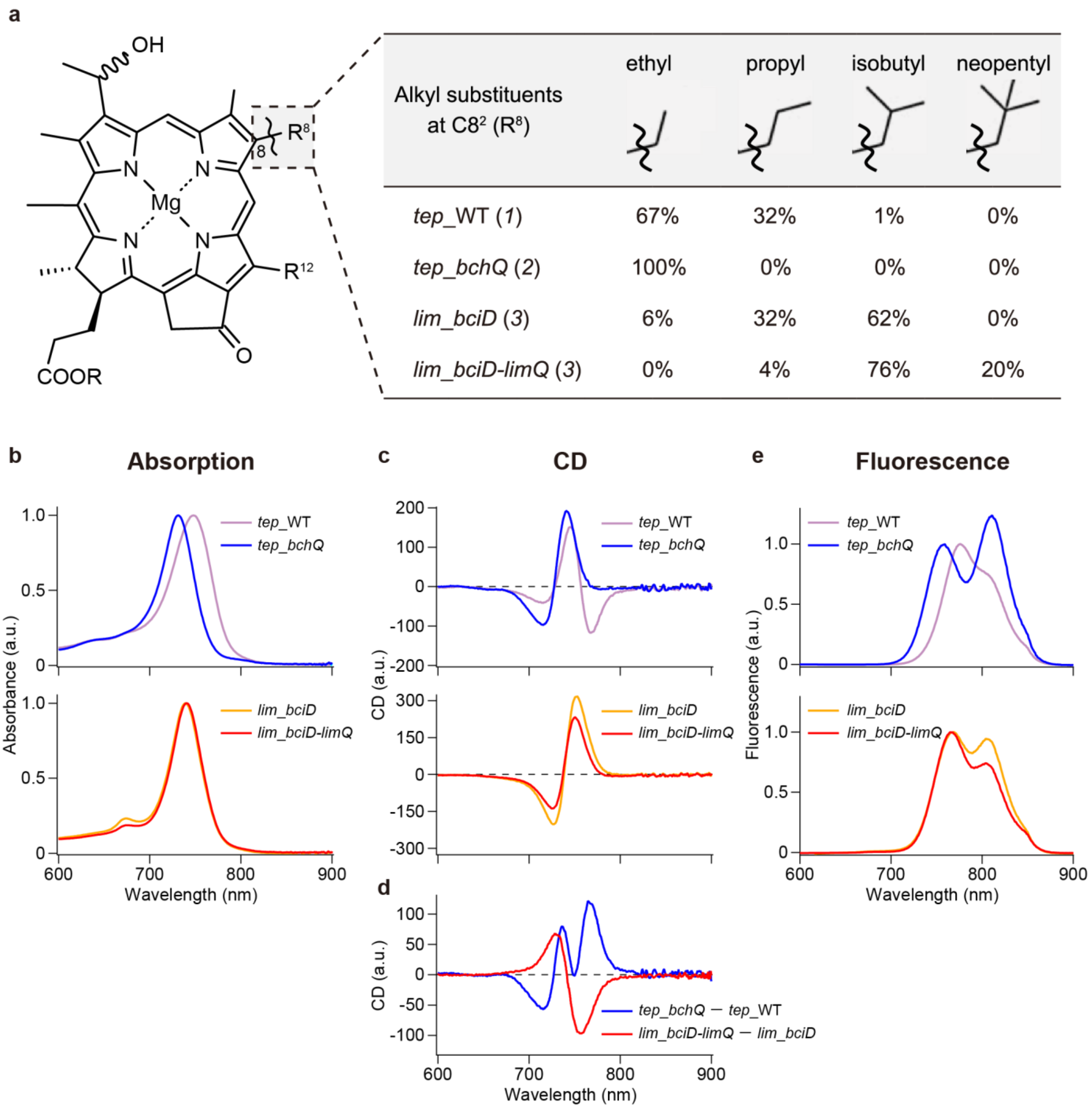

**Fig. S1 | Relative abundances of BChl *c* homologs and spectroscopic characterization of wild-type and mutant chlorosomes. a**, Molecular structure of BChl *c* (left) and relative abundances of BChl *c* homologs with different alkyl substituents at the C8$^2$ position in wild-type and mutant chlorosomes (right). References are indicated in parentheses. In *tep*_WT, ethyl and propyl substituents predominate, accounting for 67% and 32%, respectively[1]. In *tep_bchQ*, only ethyl substituents were observed[2]. In *lim_bciD*, isobutyl substituents are most abundant (62%), followed by propyl substituents (32%)[3]. By contrast, in *lim_bciD-limQ*, isobutyl and neopentyl substituents account for 76% and 20%, respectively[3]. **b**, Absorption spectra in the $Q_y$ region of *tep*_WT (light purple), *tep_bchQ* (blue), *lim_bciD* (orange), and *lim_bciD-limQ* (red) chlorosomes measured in solution and normalized to their respective peak intensities. **c**, CD spectra in the $Q_y$ region of the corresponding samples. The CD signal intensities were normalized to the maximum of the $Q_y$ absorption peak in each sample. **d**, Difference CD spectra calculated by subtracting the CD spectrum of *tep*_WT from that of *tep_bchQ* (blue) and by subtracting that of *lim_bciD* from that of *lim_bciD-limQ* (red). **e**, Fluorescence spectra of the corresponding samples normalized to their respective BChl *c* peaks in the 740–780 nm range. The absorption and fluorescence spectra of *tep*_WT chlorosomes (light purple) shown in (b) and (e), respectively, are replotted over the wavelength range of 700–870 nm in Fig. 1b. Details of the spectra are provided in Supplementary Note 1.

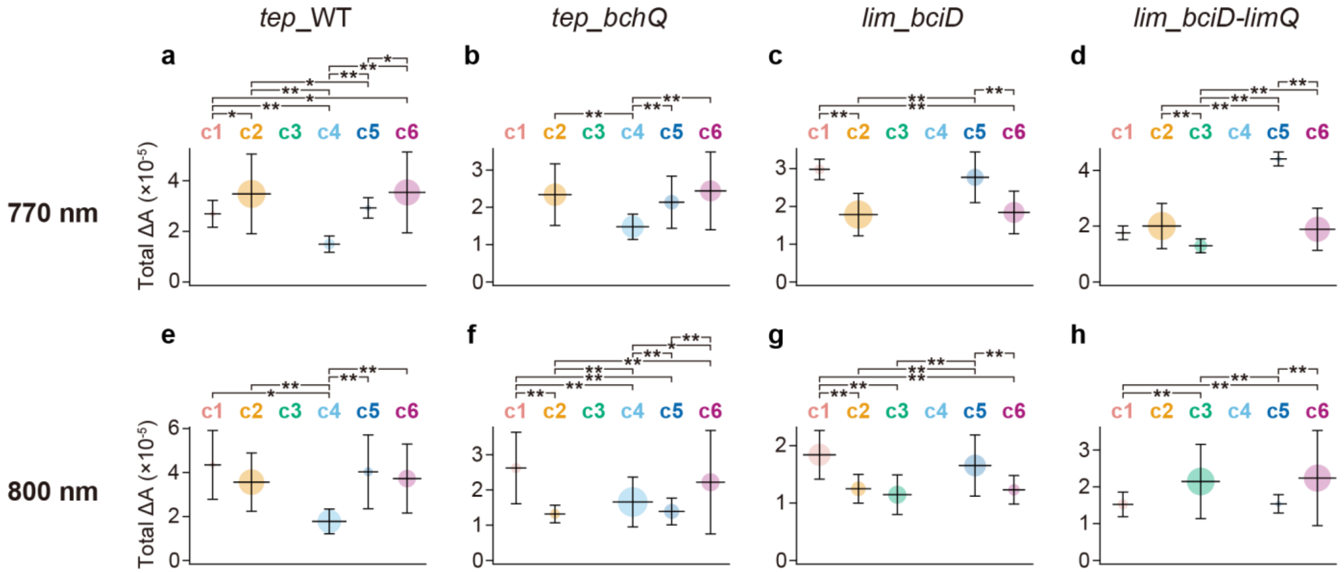

**Fig. S2 | Estimation of total $\Delta A$ associated with each $\tau$ component. a–d**, Estimated distributions of total $\Delta A$ for each $\tau$ component in *tep*_WT (a), *tep_bchQ* (b), *lim_bciD* (c), and *lim_bciD-limQ* (d) chlorosomes, obtained using the 770-nm probe. The mean values are indicated by horizontal bars, and the $1/\sqrt{e}$ full widths are indicated by vertical bars. The fitted distributions associated with each $\tau$ component are colored as in Fig. 2. The area of each circle represents the population of the corresponding $\tau$ component. Statistically significant differences assessed using Welch's *t*-test between components are denoted by asterisks (*$p < 0.05$ and **$p < 0.01$). **e–h**, Corresponding results obtained using the 800-nm probe. The estimated distributions for *tep*_WT chlorosomes measured with the 800-nm probe in (e) are replotted in Fig. 3a. Details of the analysis are provided in Supplementary Note 10.

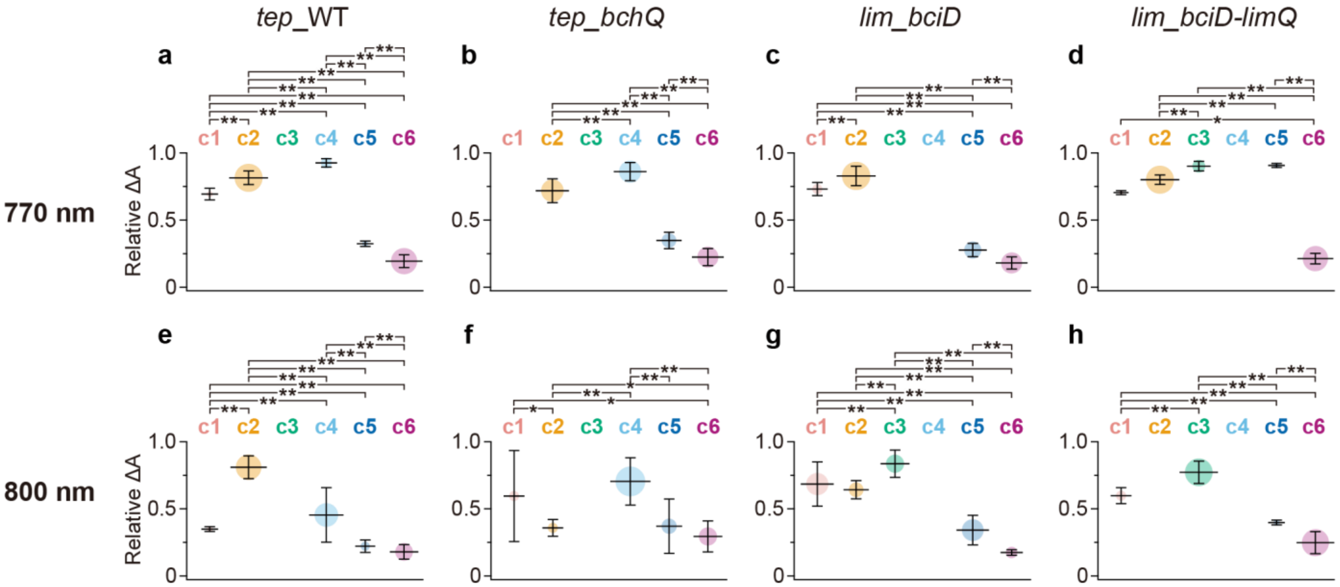

**Fig. S3 | Estimation of relative Δ*A* associated with each *τ* component.** Estimated distributions of relative Δ*A* for each *τ* component. The plots are presented in the same format as in Fig. S2. The estimated distributions for *tep*_WT (e) and *tep*_*bchQ* (f) chlorosomes measured with the 800-nm probe are replotted in Fig. 3b and d, respectively.

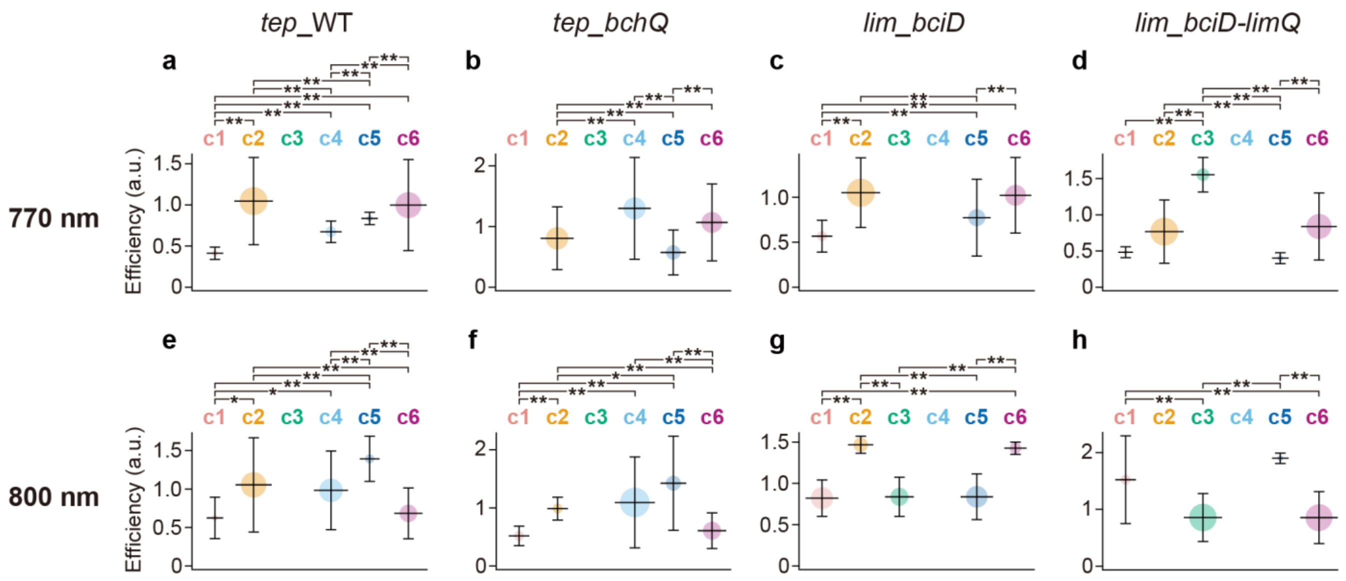

**Fig. S4 | Estimation of effective EET efficiency associated with each $\tau$ component.** Estimated distributions of effective EET efficiency for each $\tau$ component. For each sample and probe wavelength, the effective EET efficiency values were normalized to the median of the corresponding overall distribution. The plots are presented in the same format as in Fig. S2. The estimated distributions for *tep*_WT chlorosomes measured with the 800-nm probe in (e) are replotted in Fig. 3c.

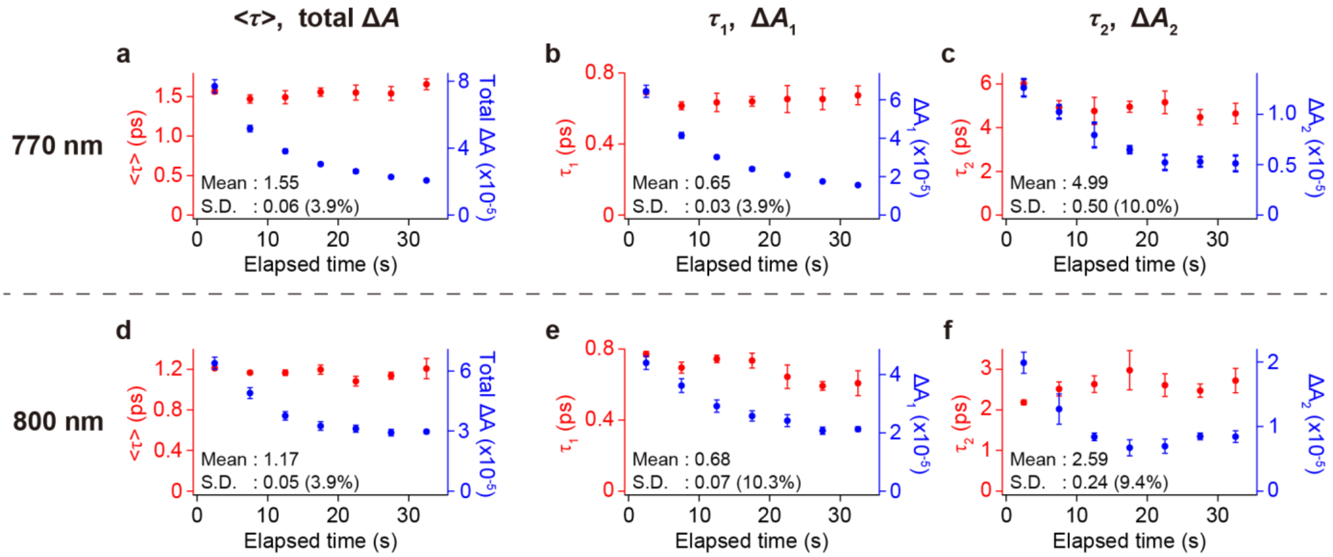

**Fig. S5 | Effects of photobleaching on excitation kinetics in chlorosomes. a–c**, Temporal changes in (a) the amplitude-weighted mean time constant $\langle\tau\rangle$ and the total amplitude $\Delta A$, (b) the shorter time constant $\tau_1$ and its amplitude $\Delta A_1$, and (c) the longer time constant $\tau_2$ and its amplitude $\Delta A_2$, estimated from TA signals averaged over 194 individual *tep*_WT chlorosomes measured with the 770-nm probe. **d–f**, Corresponding results obtained for 162 individual *tep*_WT chlorosomes measured with the 800-nm probe. The time constants and corresponding amplitudes are shown in red and blue, respectively. Mean values and standard deviations (S.D.) of the time constants over the time course, with coefficients of variation (CVs) shown in parentheses, are indicated in each panel. Error bars represent standard deviations estimated from five resampling trials, with any given chlorosome allowed to be selected at most twice in each trial until the total number of selected chlorosomes reached 194 for the 770-nm probe and 162 for the 800-nm probe. Detailed interpretations and further analyses are provided in Supplementary Note 3.

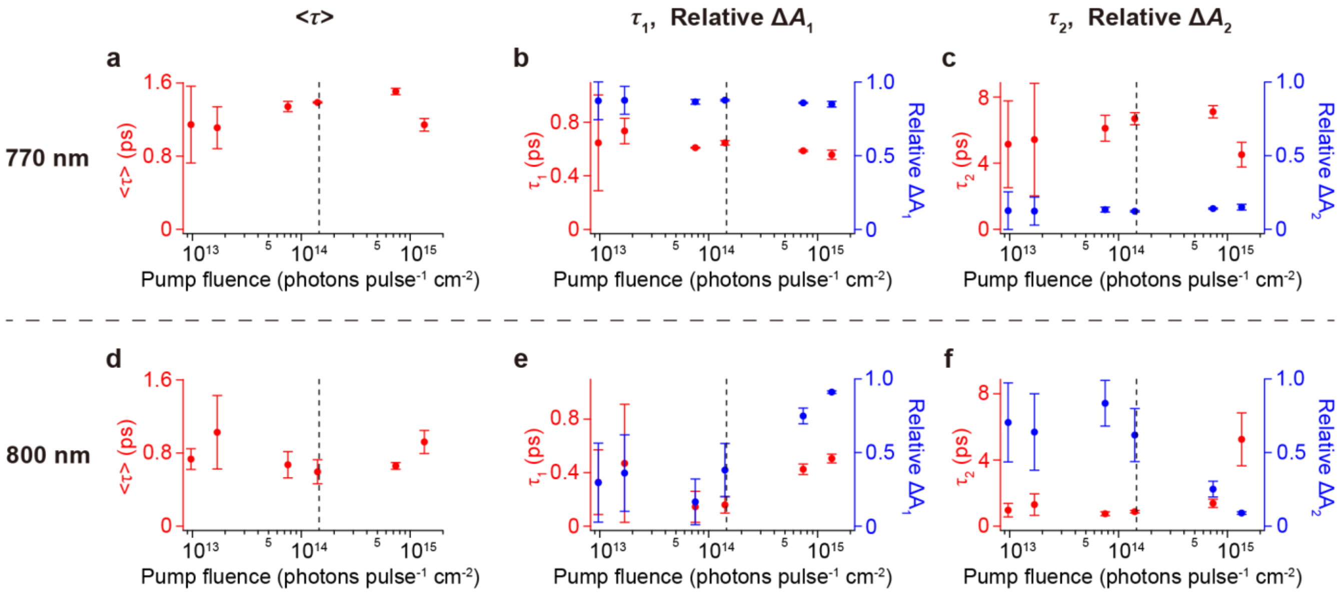

**Fig. S6 | Effects of pump fluence on excitation kinetics in chlorosomes. a**–**c**, Pump-fluence dependence of (a) the amplitude-weighted mean time constant $\langle\tau\rangle$, (b) the shorter time constant $\tau_1$ and its relative amplitude, and (c) the longer time constant $\tau_2$ and its relative amplitude, estimated from TA signals measured for an ensemble of *tep*_WT chlorosomes using the 770-nm probe. **d**–**f**, Corresponding results obtained using the 800-nm probe. The time constants and corresponding relative amplitudes are shown in red and blue, respectively. Error bars represent standard deviations estimated from three experiments. The pump fluence used in the present single-chlorosome TA measurements is indicated by a black dashed line. Detailed interpretations are provided in Supplementary Note 5.

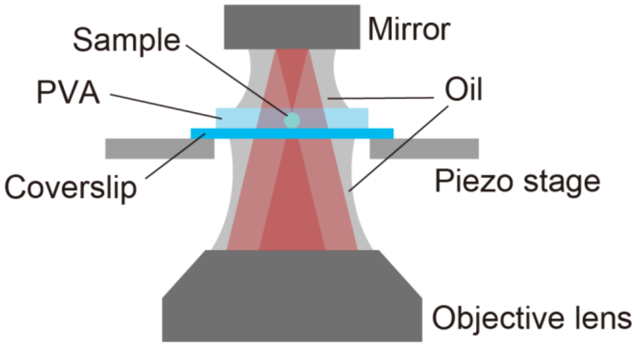

**Fig. S7 | Single-objective layout for absorption microscopy.**

The incident beam is focused by an objective onto the sample via reflection from a mirror, and the transmitted beam is collected by the same objective. The sample particles were spin-coated onto a coverslip and thus embedded in a thin polyvinyl alcohol (PVA) film. The spaces between the coverslip and objective and between the PVA film and mirror surface were filled with immersion oil to ensure refractive-index matching. Images were acquired by scanning the sample position with the piezo stage.

### Supplementary Notes

#### Supplementary Note 1: Spectroscopic properties of the wild-type and mutant chlorosomes

Absorption spectra

The absorption spectra of the wild-type and mutant chlorosomes in solution are shown in Fig. S8. The signal intensities were normalized to the maximum intensity of the $Q_y$ peak in each spectrum. The $Q_y$ peak of *tep*_WT chlorosomes is at 748 nm (Fig. S8a, light purple), whereas that of *tep_bchQ* is at 731 nm (Fig. S8a, blue). The *lim_bciD* and *lim_bciD-limQ* chlorosomes both exhibit $Q_y$ peaks at 740 nm (Fig. S8b). In all cases, the $Q_y$ peaks are red-shifted with respect to the bacteriochlorophyll (BChl) *c* monomer peak at ~670 nm, indicative of supramolecular aggregation[4]. Compared with *tep*_WT chlorosomes, the mutant chlorosomes show $Q_y$ peaks shifted to shorter wavelengths. The FWHMs of the $Q_y$ peaks are 53 nm (*tep*_WT), 43 nm (*tep_bchQ*), 41 nm (*lim_bciD*), and 40 nm (*lim_bciD-limQ*), respectively. As discussed previously, differences in the peak wavelengths and widths can be attributed to factors such as the angle between the transition dipole moment of each BChl *c* molecule and the molecular packing direction[5], the rolling angle of tubular aggregates[6], and variations in the compositional ratios of BChl *c* homologs bearing distinct side chains[2,7].

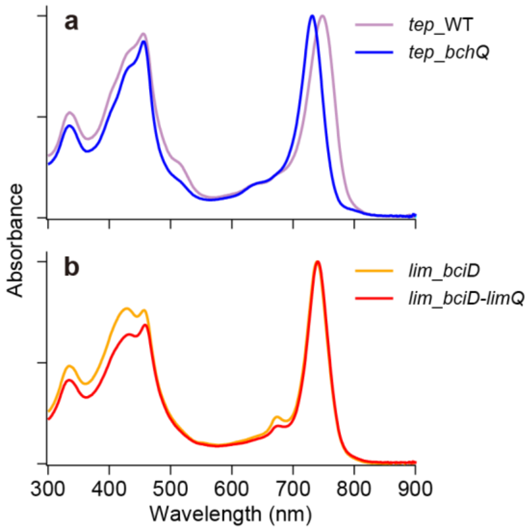

**Fig. S8 | Absorption spectra of wild-type and mutant chlorosomes.**

Absorption spectra of *tep*_WT (**a**, light purple), *tep_bchQ* (**a**, blue), *lim_bciD* (**b**, orange), and *lim_bciD-limQ* (**b**, red) chlorosomes measured in solution and normalized to their respective $Q_y$ peak intensities. The absorption spectrum of *tep*_WT chlorosomes (a, light purple) is replotted over the wavelength range of 700–870 nm in Fig. 1b. All absorption spectra are replotted over the wavelength range of 600–900 nm in Fig. S1b.

Circular dichroism (CD) spectra

The CD spectra of the chlorosomes were measured to assess mutation-induced changes in supramolecular architecture (Fig. S9). The CD signal intensities were normalized to the maximum of the $Q_y$ absorption peak in each sample. In *tep*_WT chlorosomes, the CD spectrum exhibits a sequence of negative–positive–negative bands across the $Q_y$ region from shorter to longer wavelengths (Fig. S9a, light purple). In *tep_bchQ* chlorosomes, the $Q_y$ region shows only negative–positive bands (Fig. S9a, blue), lacking the negative band at longer wavelengths observed in *tep*_WT. These observations are consistent with prior reports[7].

Cryo-electron microscopy (cryo-EM) analyses have shown that *tep*_WT chlorosomes contain both tubular and lamellar aggregates[8], suggesting that the CD spectra include contributions not only from tubular and lamellar aggregate morphologies but also from inter-aggregate interactions and contacts with the baseplate[9]. By contrast, shortening of the BChl side chain has been shown by cryo-EM to promote the formation of multilayered tubular aggregates[8]. Thus, the *tep_bchQ* spectrum is attributed mainly to tubular aggregates. To isolate the component that increases upon side-chain shortening, we calculated the difference spectrum defined as *tep_bchQ* minus *tep*_WT (Fig. S9c, blue). In the $Q_y$ region around 750 nm, this difference spectrum exhibits a negative band at shorter wavelengths and two positive bands toward longer wavelengths.

The CD spectrum of *lim_bciD* chlorosomes displays negative–positive features across the $Q_y$ region (Fig. S9b, orange), likely reflecting a mixture of tubular and lamellar contributions. Upon elongation of the BChl side chain in *lim_bciD-limQ*, the overall CD amplitude decreases (Fig. S9b, red). The difference spectrum defined as *lim_bciD-limQ* minus *lim_bciD* shows a sequence of positive–negative bands from shorter to longer wavelengths in the $Q_y$ region (Fig. S9c, red). This line shape is clearly distinct from that of the *tep_bchQ* minus *tep*_WT difference spectrum (Fig. S9c, blue), which reflects enrichment of tubular aggregates. This line shape is therefore attributed to an increased fraction of lamellar aggregates accompanying elongation of the BChl side chain.

Chlorosome CD spectra, which are known to vary substantially among samples, can be interpreted as a linear combination of two basis spectra exhibiting inverse signatures, i.e., positive–negative and negative–positive patterns from shorter to longer wavelengths[10]. The two difference spectra in Fig. S9c, each reflecting distinct aggregate morphologies, can be interpreted as linear combinations of the two basis spectra. These results suggest that the genetic modulation of pigment biosynthesis, which shortens or lengthens BChl side chains, systematically shifts the composition of chlorosomal aggregates between tubular and lamellar forms.

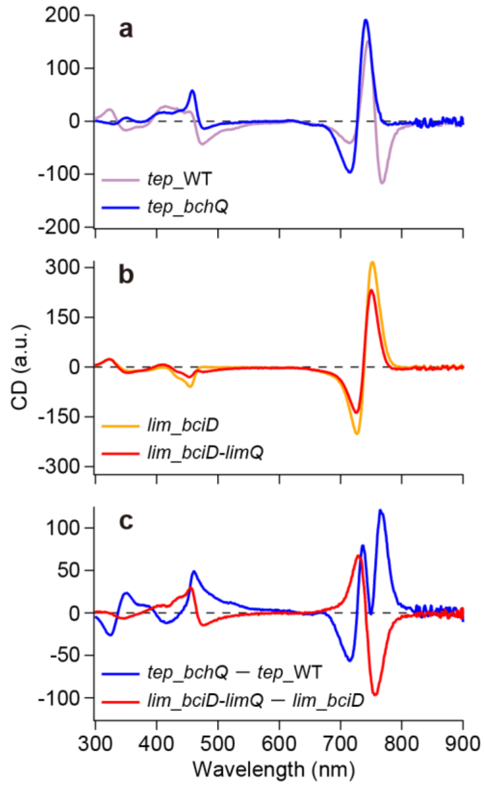

**Fig. S9 | CD spectra of wild-type and mutant chlorosomes.**

CD spectra of *tep*_WT (**a**, light purple), *tep_bchQ* (**a**, blue), *lim_bciD* (**b**, orange), and *lim_bciD-limQ* (**b**, red) chlorosomes measured in solution. The CD signal intensities were normalized to the $Q_y$ absorption maximum of each sample. (**c**) Difference spectra calculated by subtracting the CD spectrum of *tep*_WT from that of *tep_bchQ* (blue) and that of *lim_bciD* from that of *lim_bciD-limQ* (red). All CD spectra and difference spectra are replotted over the wavelength range of 600–900 nm in Fig. S1c and d, respectively.

Fluorescence spectra

The fluorescence spectra of the wild-type and mutant chlorosomes in solution exhibit multiple peaks (Fig. S10a). To identify these peaks, we fitted each spectrum with a sum of Gaussian functions (Fig. S10b–e and Table S1). In *tep*_WT chlorosomes, the spectrum includes a peak at 777 nm and shoulder peaks at 813 and 837 nm (Fig. S10b). The former arises from BChl aggregates, whereas the latter two are thought to originate from BChl *a* in the baseplate[11]. In *tep_bchQ* chlorosomes, the aggregate peak is observed at 757 nm, indicating a 20-nm blue shift relative to *tep*_WT (Fig. S10c), consistent with the shift in the $Q_y$ absorption peak (Fig. S8a). Additionally, the baseplate peaks at >800 nm show increased intensity relative to those of *tep*_WT. The spectrum of *lim_bciD* chlorosomes shows a peak at 767 nm with shoulder peaks above 800 nm (Fig. S10d), while that of *lim_bciD-limQ* chlorosomes exhibits a similar line shape without substantial peak shifts but with reduced intensity above 800 nm (Fig. S10e). Compared with *tep*_WT, the aggregate peaks at <800 nm are blue-shifted in the other three samples, whereas the baseplate peaks at >800 nm show little shift (Table S1). This suggests that the BChl side-chain mutations exert little influence on the spectral properties of the baseplate.

In chlorosomes, part of the energy absorbed by BChl aggregates is emitted from the aggregates as fluorescence, whereas excitation energy transferred to the baseplate ultimately gives rise to baseplate-derived fluorescence. Accordingly, the fraction of fluorescence arising from the baseplate relative to the total fluorescence provides an apparent measure of the energy-transfer efficiency from the aggregates to the baseplate. By dividing the summed area of the two baseplate-derived peaks at >800 nm by the total area of all three peaks, we estimated this efficiency to be 0.25 and 0.49 for *tep*_WT and *tep_bchQ*, respectively, indicating a ~2-fold enhancement upon shortening of the BChl side chain. In contrast, *lim_bciD* and *lim_bciD-limQ* showed nearly identical values of 0.37 and 0.34, respectively. These results suggest that the shorter BChl side chain, which is associated with enrichment of tubular aggregates, facilitates more efficient energy transfer to the baseplate. This supports the light-harvesting model in Fig. 3e, in which tubular aggregates mediate energy transfer to the baseplate.

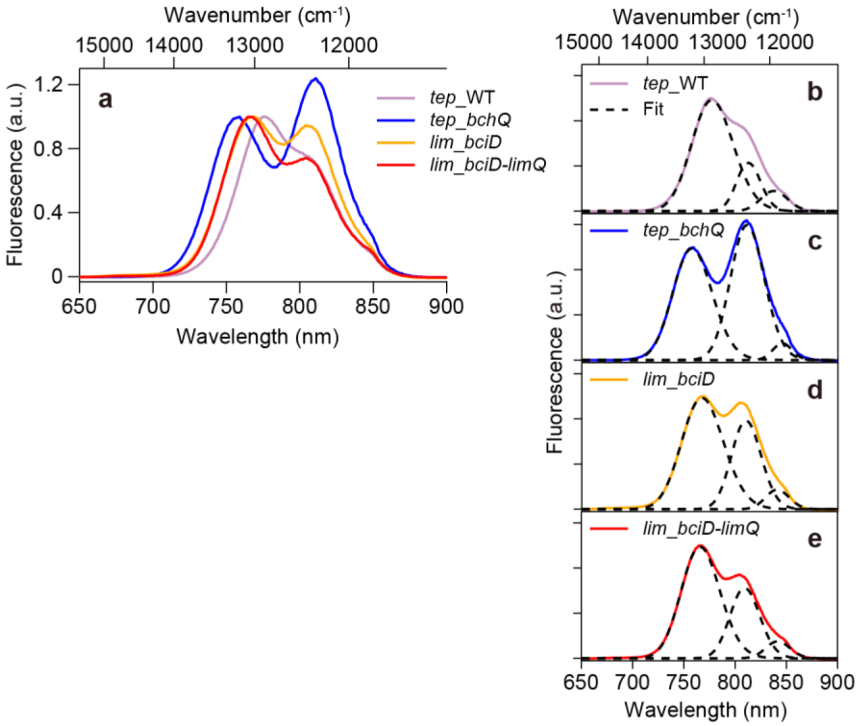

**Fig. S10 | Fluorescence spectra of wild-type and mutant chlorosomes.**

(**a**) Fluorescence spectra of *tep*_WT (light purple), *tep_bchQ* (blue), *lim_bciD* (orange), and *lim_bciD-limQ* (red) chlorosomes measured in solution and normalized to their respective BChl *c* peaks in the 740–780 nm range. (**b**–**e**) Gaussian fits decomposing the spectra into three Gaussian peaks (dashed black lines) for *tep*_WT (b), *tep_bchQ* (c), *lim_bciD* (d), and *lim_bciD-limQ* (e). The fluorescence spectrum of *tep*_WT chlorosomes shown in (a, light purple) is replotted over the wavelength range of 700–870 nm in Fig. 1b. All fluorescence spectra are replotted over the wavelength range of 600–900 nm in Fig. S1e.

**Table S1**

Population, center position, and FWHM of each peak estimated from three-component Gaussian fits to the fluorescence spectra of wild-type and mutant chlorosomes.

| | Peak 1 | | | Peak 2 | | | Peak 3 | | |
|---|---|---|---|---|---|---|---|---|---|
| | Population (%) | Center position (nm) | FWHM (nm) | Population (%) | Center position (nm) | FWHM (nm) | Population (%) | Center position (nm) | FWHM (nm) |
| *tep*_WT | 75 | 777 | 48 | 17 | 813 | 28 | 8 | 837 | 31 |
| *tep_bchQ* | 51 | 757 | 45 | 46 | 811 | 39 | 3 | 846 | 22 |
| *lim_bciD* | 63 | 767 | 48 | 32 | 810 | 34 | 5 | 841 | 27 |
| *lim_bciD-limQ* | 66 | 766 | 45 | 29 | 809 | 34 | 5 | 842 | 28 |

#### Supplementary Note 2: Redox conditions associated with excitation quenching

Green sulfur bacteria grow under anaerobic conditions, and their chlorosomes are in a reduced state. Under aerobic conditions, the excitation energy in chlorosomes is rapidly quenched[12], indicating that reducing conditions should be maintained during single-chlorosome transient absorption (TA) measurements. The fluorescence intensity of chlorosomes is higher in the reduced state, whereas it is lower in the oxidized state due to excitation quenching[13]. Accordingly, using fluorescence intensity as an indicator, we verified that the chlorosome samples used in the present TA measurements, which contained 10 mM dithionite, remained in a reduced state.

A *tep*_WT chlorosome solution at an optical density (OD) of ≈ 2 at the $Q_y$ band, containing 40 mM Tris–HCl (pH 8.0), 1% (w/v) PVA, 1 U $mL^{-1}$ protocatechuate 3,4-dioxygenase (PCD), and 2.5 mM protocatechuic acid (PCA), was spin-coated onto a coverslip after the addition of 10 mM dithionite. From fluorescence images, the fluorescence intensity of each chlorosome was estimated, and the resulting distribution is shown in Fig. S11a. The mean (median) fluorescence intensity was 1905 (1686) cps (dashed black and solid red lines, respectively). To further ensure reduction, the sample was kept at 4 °C in the dark for ~16 h after the addition of 10 mM dithionite, yet similar values of 1776 (1394) cps were obtained (Fig. S11b). By contrast, when 1 mM potassium ferricyanide was added to oxidize chlorosomes, the mean (median) intensity decreased to 312 (275) cps, which was close to the background level (Fig. S11c). Even when the excitation fluence was increased ~10-fold, the fluorescence remained low, at 506 (450) cps (Fig. S11d). These results are consistent with a previous report demonstrating that the fluorescence quantum yield of oxidized chlorosomes decreases to approximately one-fortieth of that in the reduced state[13]. Thus, chlorosomes in the presence of 10 mM dithionite were substantially more fluorescent than those under oxidized conditions, indicating that the chlorosomes used in the present TA measurements were sufficiently reduced.

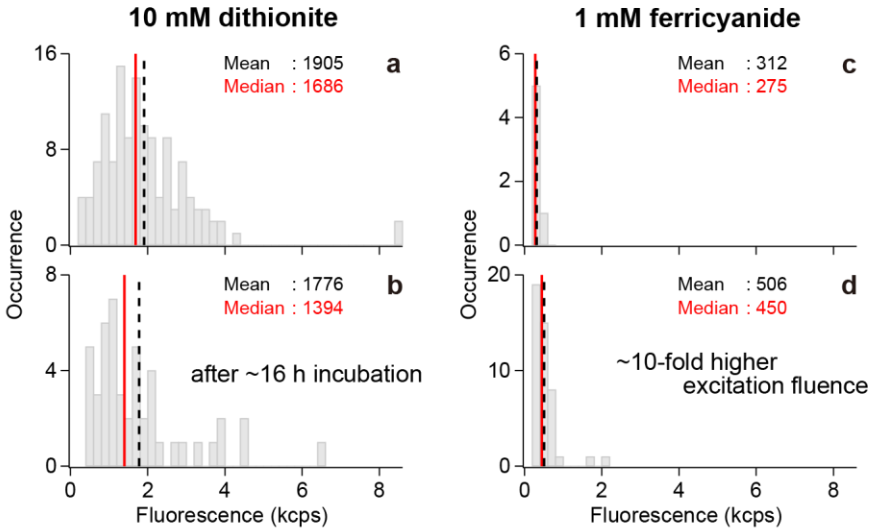

**Fig. S11 | Fluorescence intensity distributions of individual chlorosomes under reducing and oxidizing conditions.**

Fluorescence intensity distributions of individual *tep*_WT chlorosomes were measured at an excitation fluence of 1.5 × $10^{13}$ photons pulse$^{-1}$ cm$^{-2}$ (**a**) immediately after the addition of 10 mM dithionite and (**b**) approximately 16 h later. Corresponding distributions were measured in the presence of 1 mM ferricyanide (**c**) at the same excitation fluence and (**d**) at a higher excitation fluence of 1.5 × $10^{14}$ photons pulse$^{-1}$ cm$^{-2}$. The numbers of chlorosomes used to construct the distributions in (a) to (d) were 130, 47, 6, and 45, respectively. The bin size is 0.2 kcps. The mean and median of each distribution are indicated by dashed black and solid red lines, respectively.

#### Supplementary Note 3: Photobleaching process

Suppression by oxygen-scavenging systems

Oxygen dissolved in the sample solution can be converted into reactive oxygen species through photoexcitation processes, leading to photobleaching. To mitigate this risk, an enzymatic oxygen-scavenging system was added to reduce dissolved oxygen[14,15]. In addition, the sample was spin-coated onto a coverslip and immediately sealed using an O-ring and an additional coverslip to suppress oxygen ingress. The time course of the fluorescence intensity of an individual *tep*_WT chlorosome measured immediately after sealing is shown in Fig. S12a. The excitation wavelength was 720 nm and the photon fluence was 3.4 × $10^{13}$ photons pulse$^{-1}$ cm$^{-2}$. The photobleaching time constant was determined by single-exponential fitting (Fig. S12a, blue). Measurements of 43 chlorosomes yielded a distribution of photobleaching times with a mean (median) of 10.9 (10.4) s (Fig. S12b).

By contrast, in a sample measured ~8 h after the oxygen-scavenging system was added and the sample was sealed, photobleaching was slower (Fig. S12c). From 42 chlorosomes, the mean (median) photobleaching time was estimated to be 48.7 (26.6) s (Fig. S12d), corresponding to a 4.5-fold (2.6-fold) increase relative to the sample measured immediately after sealing (Fig. S12b).

This improvement likely reflected further depletion of oxygen in the sealed volume over time, which suppressed photobleaching. Accordingly, single-chlorosome TA measurements were performed on samples stored for at least one night after sealing.

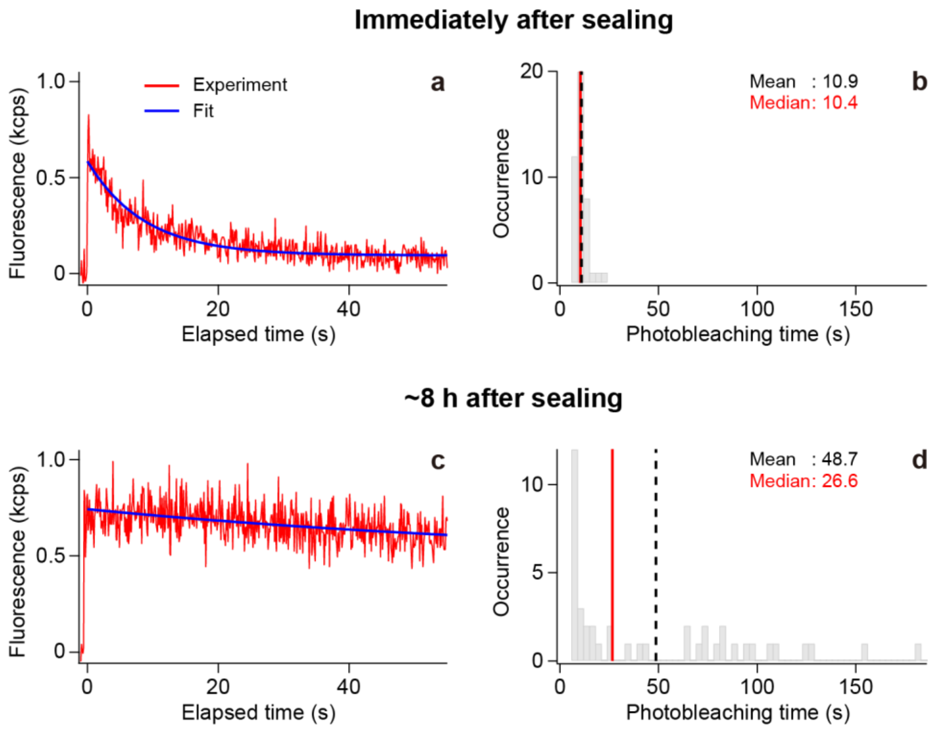

**Fig. S12 | Photobleaching times of individual chlorosomes in the presence of an oxygen-scavenging system.**

(**a**) Representative time-dependent decrease in fluorescence intensity due to photobleaching, measured from an individual *tep*_WT chlorosome immediately after sealing the sample in the presence of the oxygen-scavenging system. The blue line shows a single-exponential fit. (**b**) Distribution of photobleaching times. (**c**) Representative time-dependent decrease in fluorescence intensity measured ~8 h after sealing, together with a single-exponential fit (blue). (**d**) Corresponding distribution of photobleaching times. The numbers of chlorosomes used to construct the distributions in (b) and (d) were 43 and 42, respectively. The mean and median of each distribution are indicated by dashed black and solid red lines, respectively. The bin size for the distributions is 3 s.

TA decay kinetics perturbed by photobleaching

To assess how photobleaching affects the excitation energy transfer (EET) process in chlorosomes, we analyzed temporal changes in TA signals. TA signals from individual *tep*_WT chlorosomes were recorded at 0.25-s intervals (4 Hz), then averaged over 194 chlorosomes (770-nm probe) and 162 chlorosomes (800-nm probe). To improve the signal-to-noise ratio (SNR), the TA signals were averaged over 5-s intervals, yielding a time series of TA traces at 5-s intervals. At each elapsed time point since the start of acquisition, the TA trace was fitted with a biexponential function to estimate the time constants ($\tau_1$ and $\tau_2$) and the corresponding amplitudes

($\Delta A_1$ and $\Delta A_2$) for the two decay components (see Supplementary Note 6). We then obtained time courses of $\tau$ and $\Delta A$ for each component, as well as those of total $\Delta A$, defined as the sum of the amplitudes of the two decay components, and the amplitude-weighted mean time constant $\langle\tau\rangle$, given by $(\Delta A_1 \cdot \tau_1 + \Delta A_2 \cdot \tau_2)$ / total $\Delta A$ (Fig. S5).

Error bars for each parameter were estimated from repeated resampling of the data, as described below for the case of the 770-nm probe. First, when calculating the averaged TA signal from the 194 chlorosomes measured with the 770-nm probe, the chlorosomes were randomly resampled, with each chlorosome allowed to be selected at most twice. Then, the resampled average TA trace was fitted to obtain $\Delta A$, $\tau$, total $\Delta A$, and $\langle\tau\rangle$ for each 5-s interval. Finally, this resampling procedure was repeated five times to estimate the standard deviation of each parameter. These standard deviations are indicated by error bars for each data point in Fig. S5.

The results for the 770-nm probe are shown in Fig. S5a–c. The total $\Delta A$ decayed exponentially with elapsed time (Fig. S5a, blue), indicating gradual photobleaching. By contrast, $\langle\tau\rangle$ remained nearly constant (Fig. S5a, red). Examining the two time constants separately revealed that $\tau_1$ for the fast decay component varied with a standard deviation of ~0.03 ps and the coefficient of variation (CV), defined as the standard deviation divided by the mean, was only 3.9% (Fig. S5b, red). Likewise, $\tau_2$ of the slow decay component exhibited a small CV of 10.0% (Fig. S5c, red). Similar behavior was observed with the 800-nm probe (Fig. S5d–f), where CVs of 10.3% and 9.4% were estimated for $\tau_1$ and $\tau_2$, respectively. Accordingly, the time constants changed only minimally during photobleaching. Nevertheless, photobleaching times differed among chlorosomes exhibiting the components c1–c6, as described in the following section.

Photobleaching times associated with each $\tau$ component

To quantify photobleaching times for each $\tau$ component, we performed a two-dimensional (2D) distribution analysis of the time constant $\tau$ versus photobleaching time (see Supplementary Note 10). For each *tep*_WT chlorosome, the TA signals recorded at 0.25-s intervals were averaged over 0.5-s intervals. The averaged trace for each 0.5-s interval was integrated along the delay-time axis (the horizontal axis) to obtain the total signal intensity, which was then plotted against the elapsed time since the start of acquisition. The temporal change in this integrated TA signal intensity was fitted with a single-exponential function to determine the photobleaching time for each chlorosome. Consequently, both $\tau$ values and the photobleaching time were obtained for individual chlorosomes, enabling construction of the 2D distributions (Fig. S13a, c). Chlorosomes with estimated photobleaching times longer than the total acquisition time comprised ~9.6% (34/356) of the data set. Because the fitted photobleaching times for these cases were unreliable, we excluded them from subsequent analyses and analyzed the remaining chlorosomes.

With the 770-nm probe (Fig. S13b), the photobleaching times for chlorosomes exhibiting the lamellar components c2 and c6 were longer than those for chlorosomes exhibiting the tubular components c1 and c5, with $p = 1.8 \times 10^{-4}$ for c1 versus c2 and $p = 6.8 \times 10^{-8}$ for c5 versus c6, indicating significant differences. With the 800-nm probe (Fig. S13d), the prolongation of photobleaching times was even more pronounced for c6 relative to c5 ($p = 1.9 \times 10^{-9}$). These results are consistent with the light-harvesting model proposed in Fig. 3e, in which excitation energy flows from lamellar to tubular aggregates. In this picture, tubular aggregates situated downstream in the EET pathway would experience greater accumulation of excitation energy and thus have a higher probability of photobleaching, whereas lamellar aggregates upstream in the pathway would accumulate energy less readily and therefore exhibit longer photobleaching times.

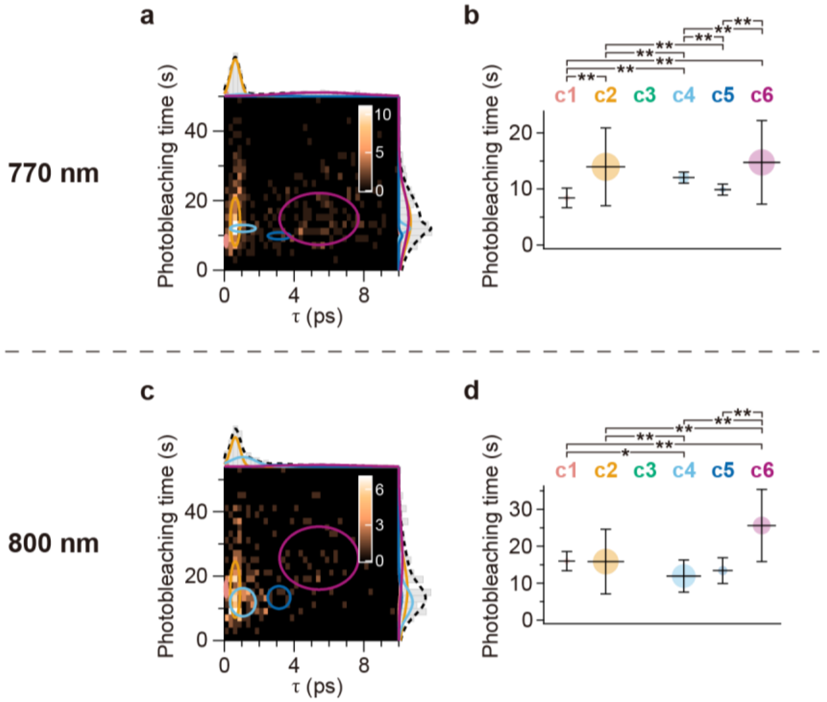

**Fig. S13 | Estimation of photobleaching times associated with each $\tau$ component.**
(**a**) 2D distribution of photobleaching time versus $\tau$ for *tep*_WT chlorosomes, obtained using the 770-nm probe. The bin sizes are 2.0 s for photobleaching time and 0.25 ps for $\tau$. The circles denote contours at $1/\sqrt{\mathrm{e}}$ of the peak height (i.e., one-standard-deviation contours) for each $\tau$ component, estimated from a five-component 2D Gaussian fit. The projections of the 2D distribution onto the $x$- and $y$-axes are shown as gray histograms at the top and right of the panel, respectively. The fitted distributions associated with each $\tau$ component are represented by solid lines colored as in Fig. 2, and their sum is indicated by the dashed black line. (**b**) Estimated distributions of photobleaching time for each $\tau$ component. The mean values are indicated by horizontal bars, and the $1/\sqrt{\mathrm{e}}$ full widths are indicated by vertical bars. The area of each circle represents the population of the corresponding $\tau$ component. Statistically significant differences assessed using Welch's $t$-test between components are denoted by asterisks (*$p < 0.05$ and **$p < 0.01$). (**c**, **d**) Corresponding results obtained using the 800-nm probe.

#### Supplementary Note 4: Effects of PVA and oil coating

The individual chlorosomes used for the TA measurements were spin-coated with PVA and then covered with an oil-coated mirror. To assess how the coating treatment affected sample conditions, we compared the fluorescence spectrum averaged over individual *tep*_WT chlorosomes under PVA- and oil-coated conditions with that in solution (Fig. S14). The fluorescence spectra of individual chlorosomes were acquired with a charge-coupled device (CCD) spectrometer (Kymera 328i and iDus 416, Andor).

The fluorescence spectrum averaged over 30 individual *tep*_WT chlorosomes under coated conditions (Fig. S14, cyan) showed minor alterations at >800 nm compared with that in solution (Fig. S14, light purple). These alterations were quantified by Gaussian fitting, which decomposed the spectrum into three peaks, as was done for the spectrum measured in solution (Fig. S10b, black dashed lines). The fit suggested that the BChl-aggregate-derived peak at <800 nm was largely unchanged, whereas for the two baseplate-derived peaks at >800 nm, the shorter-wavelength peak decreased and the longer-wavelength peak increased and broadened (Table S2). Given that the microscopic structures of chlorophyll-derivative aggregates were barely affected by a nearly identical coating treatment[16], even though these aggregates, unlike chlorosomes, were not enclosed by a lipid layer, the coating treatment likely had only a minimal effect on the BChl aggregates within chlorosomes. The observed spectral differences were therefore attributed to interactions of the baseplate with the PVA environment. Nevertheless, the apparent EET efficiency from the BChl aggregates to the baseplate, calculated by dividing the summed area of the two baseplate-derived peaks at >800 nm by the total area of all three peaks as described in Supplementary Note 1, was estimated to be 0.26, closely matching the value in solution (0.25). This result suggests that the EET process from the aggregates to the baseplate was largely unaffected by the coating treatment.

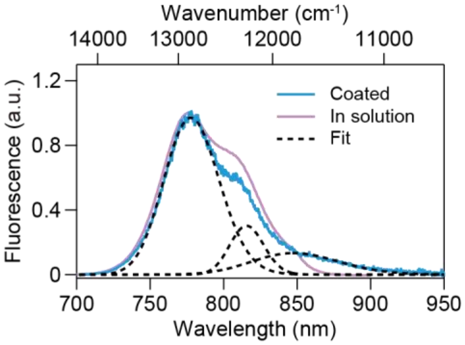

**Fig. S14 | Comparison of fluorescence spectra of wild-type chlorosomes measured under PVA- and oil-coated conditions and in solution.**

The fluorescence spectrum averaged over 30 individual *tep*_WT chlorosomes measured under PVA- and oil-coated conditions (cyan) is compared with that measured in solution (light purple), which was replotted from Fig. S10. The spectra were normalized to their respective main peaks. The three peaks estimated from a three-component Gaussian fit to the spectrum measured under PVA- and oil-coated conditions are overlaid (black dashed lines).

**Table S2**

Population, center position, and FWHM of each peak estimated from a three-component Gaussian fit to the averaged fluorescence spectrum of *tep*_WT chlorosomes under PVA- and oil-coated conditions, compared with the corresponding values for *tep*_WT chlorosomes in solution listed in Table S1.

| | Peak 1 | | | Peak 2 | | | Peak 3 | | |
|---|---|---|---|---|---|---|---|---|---|
| | Population (%) | Center position (nm) | FWHM (nm) | Population (%) | Center position (nm) | FWHM (nm) | Population (%) | Center position (nm) | FWHM (nm) |
| In solution | 75 | 777 | 48 | 17 | 813 | 28 | 8 | 837 | 31 |
| Coated condition | 74 | 778 | 44 | 13 | 816 | 27 | 13 | 848 | 69 |

#### Supplementary Note 5: Excitation-fluence dependence

Variations in time constants at higher pump fluences

The pump-fluence dependence of the TA signal was examined as follows. A chlorosome sample was prepared at a high concentration by sequentially depositing twelve 5-μL aliquots of a solution containing *tep*_WT chlorosomes at an OD of ≈ 15 at the $Q_y$ band, 10 mM dithionite, 1 U $mL^{-1}$ PCD, 2.5 mM PCA, and 20 mM Tris–HCl (pH 8.0) onto a coverslip. The highly concentrated ensemble sample provided sufficient TA signal amplitude even at low pump fluences. A 5-μL aliquot of a solution containing 1% (w/v) PVA and 40 mM Tris–HCl (pH 8.0) was then spin-coated at 3000 rpm for 1 min over the sample, after which an oil-coated mirror was placed above it as illustrated in Fig. S7. To minimize photobleaching effects, the sample stage was moved continuously at 1 μm $ms^{-1}$ to shift the focal position during TA acquisition. TA signals accumulated over 50 s were fitted with a biexponential function. This procedure was repeated three times to estimate the mean and standard deviation of the time constants $\tau_1$ and $\tau_2$, the relative $\Delta A$ values ($\Delta A_1$ / total $\Delta A$ and $\Delta A_2$ / total $\Delta A$, where total $\Delta A = \Delta A_1 + \Delta A_2$), and the amplitude-weighted mean time constant $\langle \tau \rangle$. The pump-fluence dependence of these parameters is shown in Fig. S6 for pump fluences ranging from $9.7 \times 10^{12}$ to $1.4 \times 10^{15}$ photons $pulse^{-1}$ $cm^{-2}$.

With the 770-nm probe, $\langle \tau \rangle$ remained unchanged within the error bars at lower pump fluences but decreased by 24% when the pump fluence was increased from $7.4 \times 10^{14}$ to $1.4 \times 10^{15}$ photons $pulse^{-1}$ $cm^{-2}$ (Fig. S6a). The relative $\Delta A_1$ and $\Delta A_2$ remained nearly constant over the whole pump-fluence range (Fig. S6b, c, blue). Given that $\tau_1$ for the fast decay component showed little change (Fig. S6b, red), this reduction can be attributed mainly to a shortening of $\tau_2$ for the slow decay component (Fig. S6c, red). At the high pump fluence of $1.4 \times 10^{15}$ photons $pulse^{-1}$ $cm^{-2}$, $\tau_1$ and $\tau_2$ were 0.6 and 4.5 ps, respectively.

With the 800-nm probe, $\langle \tau \rangle$ increased slightly at pump fluences higher than $7.4 \times 10^{14}$ photons $pulse^{-1}$ $cm^{-2}$ (Fig. S6d). While $\tau_1$ remained unchanged within the error bars at lower fluences, it increased at fluences above $7.4 \times 10^{14}$ photons $pulse^{-1}$ $cm^{-2}$, with an accompanying increase in relative $\Delta A_1$ (Fig. S6e). Similarly, $\tau_2$ increased, whereas relative $\Delta A_2$ decreased with increasing pump fluence (Fig. S6f). At $1.4 \times 10^{15}$ photons pulse$^{-1}$ cm$^{-2}$, $\tau_1$ and $\tau_2$ were 0.5 and 5.2 ps, respectively, which are similar to those obtained with the 770-nm probe.

In summary, with both the 770-nm and 800-nm probes, changes in the time constants were observed when the pump fluence exceeded $7.4 \times 10^{14}$ photons pulse$^{-1}$ cm$^{-2}$. At a pump fluence of $1.4 \times 10^{14}$ photons pulse$^{-1}$ cm$^{-2}$, comparable to that used in the present single-chlorosome TA measurements, such undesired variations associated with high pump fluence were not clearly observed. At this pump fluence, $\langle \tau \rangle$ was estimated to be 1.4 ps with the 770-nm probe and 0.6 ps with the 800-nm probe. These $\langle \tau \rangle$ values are 1.1- to 1.8-fold smaller than those obtained in the single-chlorosome TA measurements (Fig. S16a, e), likely due to subtle differences in the redox conditions of the sample (see Supplementary Note 2). Nevertheless, given that the decay kinetics remained largely unaffected up to approximately five times the pump fluence used in the present single-chlorosome TA measurements, the effects of high excitation are expected to be negligible in these TA measurements.

Interpretation of the excitation-fluence dependence based on the light-harvesting model

Under strong excitation, multiple excited states can be created, leading to exciton–exciton annihilation and consequent shortening of excitation lifetimes[17]. In contrast, with the 800-nm probe, we observed increases in the time constants in chlorosomes at high pump fluence. Although this might seem counterintuitive at first glance, the high-fluence behavior observed in the ensemble measurements may be explained by the light-harvesting model proposed in Fig. 3e.

With the 800-nm probe at low pump fluence, $\tau_2$ was ~1 ps (Fig. S6f). This value corresponds to the c4 component, assigned to EET from tubular aggregates to the baseplate, suggesting that EET to the baseplate proceeds efficiently under weak illumination. When the pump fluence was raised to $1.4 \times 10^{15}$ photons pulse$^{-1}$ cm$^{-2}$, both the 770-nm and 800-nm probes yielded $\tau_1 \approx 0.6$ ps and $\tau_2 \approx 5$ ps, which correspond to the lamellar components c2 and c6, respectively. In other words, under strong illumination, the c4 component corresponding to EET from tubular aggregates to the baseplate was not clearly observed. Moreover, the c5 component, ascribed to EET within tubular aggregates, was also not prominent. At high excitation fluence, triplet states can accumulate on the baseplate[18], which is the terminal energy acceptor in chlorosomes, potentially impeding further transfer of excitation energy from upstream tubular aggregates. Furthermore, if the tubular aggregates also undergo excited-state saturation, lamellar EET components may preferentially remain detectable. Thus, the observed excitation-fluence dependence is consistent with the light-harvesting model (Fig. 3e).

#### Supplementary Note 6: Fitting analysis of TA signals

The number of decay components and the instrument response function (IRF)

To estimate time constants from the TA signals, we performed a fitting analysis using the following response function, obtained by convolving exponential decays with a Gaussian IRF:

$$\mathrm{f}(t) = y_0 + \frac{1}{2}\sum_{i=1}^{N} A_i \exp\left(\frac{t_0 - t}{\tau_i} + \frac{t_\mathrm{w}^2}{4\tau_i^2}\right) \mathrm{erfc}\left(\frac{t_0 - t + \frac{t_\mathrm{w}^2}{2\tau_i}}{t_\mathrm{w}}\right), \quad (1)$$

where $y_0$ is the baseline, $N$ is the number of exponential components, $t_0$ is a temporal offset, $t_w$ is the 1/e half-width of the IRF, and $A_i$ and $\tau_i$ are the amplitude and time constant of component $i$, respectively. erfc denotes the complementary error function, defined as

$$\mathrm{erfc}(t) = \frac{2}{\sqrt{\pi}}\int_t^\infty \exp(-z^2)\ \mathrm{d}z. \tag{2}$$

We first determined the number of decay components and the IRF width by fitting averaged TA traces. These averaged TA traces were constructed using only individual traces whose mean amplitude over the five data points near 0 ps exceeded three times the standard deviation of the background signal measured at positions without chlorosomes. For *tep*_WT chlorosomes measured with the 770-nm probe, 191 TA traces were averaged (Fig. S15a, red). The same averaging procedure was applied to the mutant chlorosomes (Fig. S15b–d, red). A global fit was then performed on the four averaged traces from *tep*_WT and the three mutants, with the IRF width shared as a global parameter to identify the decay components. To ensure uniform weighting across traces in the global fit, each averaged trace was normalized to unity prior to fitting. The residual sum of squares (RSS), defined as the sum of squared residuals between the data and the fitted curves, was used as a measure of goodness of fit and compared among models in which $N$ ranged from 1 to 3.

A one-component model was insufficient to reproduce the experimental traces (Fig. S15a–d, cyan). Introducing a second component yielded good agreement (Fig. S15a–d, blue), reducing RSS to 17–36% of that for the one-component fit (Table S3a, b). In contrast, although introducing a third component decreased RSS to 15–27% of that for the one-component fit (Table S3c), the results were almost indistinguishable from those of the two-component fit (Fig. S15a–d, green). Therefore, we concluded that two decay components were sufficient. The IRF obtained from the global analysis had an FWHM of 313 fs (Table S3b).

The TA signals recorded with the 800-nm probe were similarly examined. A one-component model did not reproduce the experimental traces, whereas a two-component model reproduced them (Fig. S15e–h, cyan and blue, respectively), with RSS decreasing to 5–80% of that for the one-component fit (Table S3d, e). Introducing a third component again produced results nearly identical to those obtained with the two-component fit (Fig. S15e–h, green). Although adding the third component also reduced RSS relative to the one-component fit (Table S3f), the RSS values remained comparable to those obtained with the two-component fit. These results were similar to those obtained with the 770-nm probe. Thus, two decay components were also sufficient for the 800-nm probe. The IRF FWHM from the global analysis was 247 fs (Table S3e).

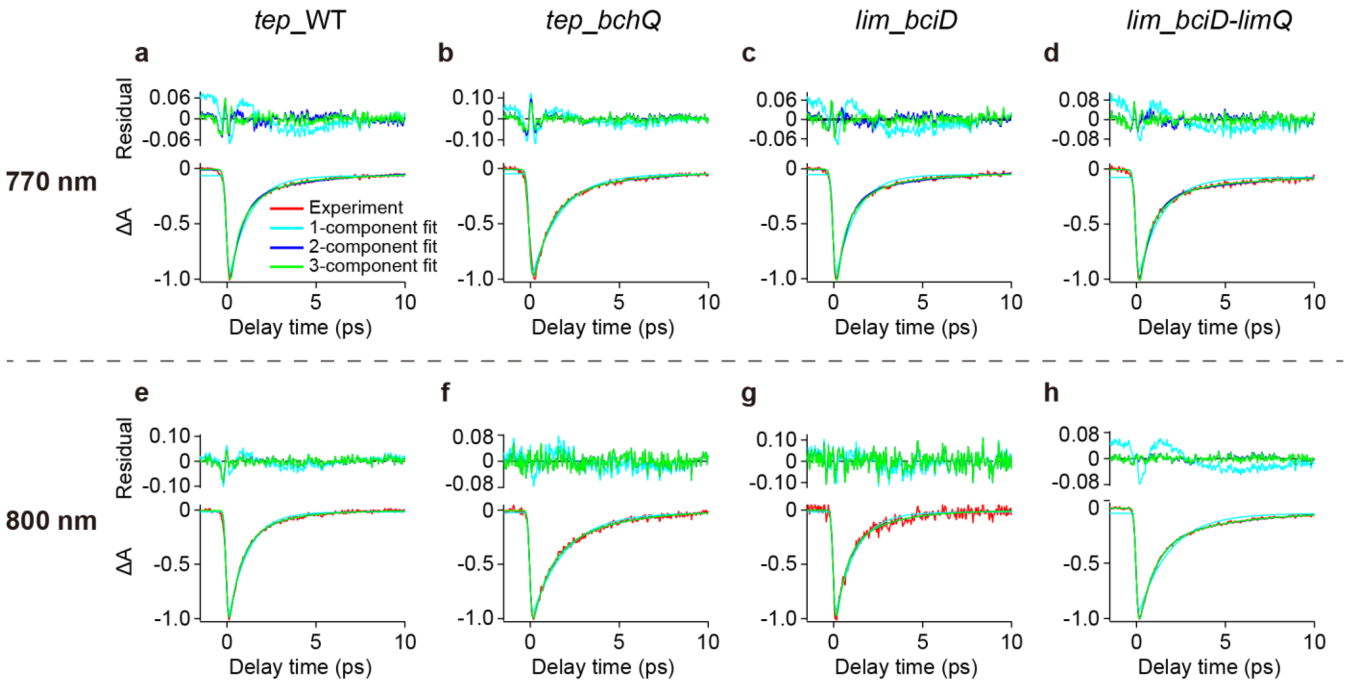

**Fig. S15 | Exponential fits to TA signals from wild-type and mutant chlorosomes.**

(**a**–**h**) TA signals (red) measured from *tep*_WT (a, e), *tep_bchQ* (b, f), *lim_bciD* (c, g), and *lim_bciD-limQ* (d, h) chlorosomes using probes at 770 nm (a–d) and 800 nm (e–h). Each trace was obtained by averaging TA signals from 191, 138, 162, and 123 chlorosomes for the 770-nm probe and 166, 95, 42, and 136 chlorosomes for the 800-nm probe, respectively. The signal intensities were normalized to their respective minima. For each probe wavelength, global fits to the four averaged traces from the wild type and three mutants were performed using a shared IRF width as a global parameter. Fitted curves obtained from the global fits using one, two, and three components (cyan, blue, and green, respectively) are superimposed. Residuals from each fit are shown in the upper panels.

**Table S3**

Parameters estimated from one-, two-, and three-component global exponential fits to the averaged TA signals measured for wild-type and mutant chlorosomes using probes at 770 nm (**a**–**c**) and 800 nm (**d**–**f**).

| 770 nm probe | **a** 1–component fit | | | **b** 2–component fit | | | | **c** 3–component fit | | | | |
|---|---|---|---|---|---|---|---|---|---|---|---|---|
| | τ (ps) [ΔA] | IRF (fs) | RSS | τ (ps) [ΔA] | | IRF (fs) | RSS | τ (ps) [ΔA] | | | IRF (fs) | RSS |
| | $\tau_1$ | | | $\tau_1$ | $\tau_2$ | | | $\tau_1$ | $\tau_2$ | $\tau_3$ | | |
| *tep*_WT | 1.2 [1] | | 0.42 (100%) | 0.7 [0.81] | 5.3 [0.19] | | 0.09 (21%) | 0.3 [0.51] | 1.2 [0.40] | 9.8 [0.09] | | 0.07 (18%) |
| *tep_bchQ* | 1.6 [1] | 238 | 0.51 (100%) | 1.1 [0.80] | 5.8 [0.20] | 313 | 0.18 (36%) | 0.9 [0.67] | 3.1 [0.31] | 402.7 [0.02] | 345 | 0.14 (27%) |
| *lim_bciD* | 1.2 [1] | | 0.49 (100%) | 0.6 [0.81] | 5.0 [0.19] | | 0.12 (24%) | 0.3 [0.55] | 1.3 [0.37] | 9.4 [0.08] | | 0.10 (21%) |
| *lim_bciD-limQ* | 1.3 [1] | | 0.85 (100%) | 0.7 [0.80] | 8.0 [0.20] | | 0.15 (17%) | 0.3 [0.47] | 1.2 [0.41] | 11.7 [0.12] | | 0.13 (15%) |

| 800 nm probe | **d** 1–component fit | | | **e** 2–component fit | | | | **f** 3–component fit | | | | |
|---|---|---|---|---|---|---|---|---|---|---|---|---|
| | τ (ps) [ΔA] | IRF (fs) | RSS | τ (ps) [ΔA] | | IRF (fs) | RSS | τ (ps) [ΔA] | | | IRF (fs) | RSS |
| | $\tau_1$ | | | $\tau_1$ | $\tau_2$ | | | $\tau_1$ | $\tau_2$ | $\tau_3$ | | |
| *tep*_WT | 1.1 [1] | | 0.21 (100%) | 0.6 [0.69] | 2.1 [0.31] | | 0.09 (42%) | 0.6 [0.71] | 2.1 [0.19] | 2.4 [0.10] | | 0.09 (41%) |
| *tep_bchQ* | 1.9 [1] | 204 | 0.31 (100%) | 0.8 [0.52] | 3.0 [0.48] | 247 | 0.16 (53%) | 0.8 [0.52] | 2.6 [0.19] | 3.4 [0.29] | 249 | 0.16 (53%) |
| *lim_bciD* | 1.2 [1] | | 0.76 (100%) | 0.6 [0.68] | 2.4 [0.32] | | 0.60 (80%) | 0.6 [0.66] | 2.0 [0.23] | 3.3 [0.11] | | 0.61 (80%) |
| *lim_bciD-limQ* | 1.8 [1] | | 0.55 (100%) | 0.9 [0.75] | 6.2 [0.25] | | 0.03 (5%) | 0.8 [0.66] | 3.1 [0.25] | 13.2 [0.09] | | 0.03 (5%) |

Fitting analysis of TA signals from individual chlorosomes

Using the IRF widths determined above as fixed parameters, we performed two-component fits to TA signals from individual chlorosomes (Fig. S16). Among the 1589 chlorosomes initially analyzed across the wild-type and mutant samples, 0.4% (7/1589) exhibited a minor fitted component with a small negative amplitude, and these chlorosomes were excluded from the present analysis. For the subsequent statistical analysis, we used the fitted parameters obtained for chlorosomes whose TA signals had an SNR > 3, where the SNR was calculated as the ratio of the signal intensity, defined as the maximum value of the fitted curve, to the noise level, defined as the standard deviation of the background signal. The numbers of chlorosomes included in the statistical analysis are listed in Table S4.

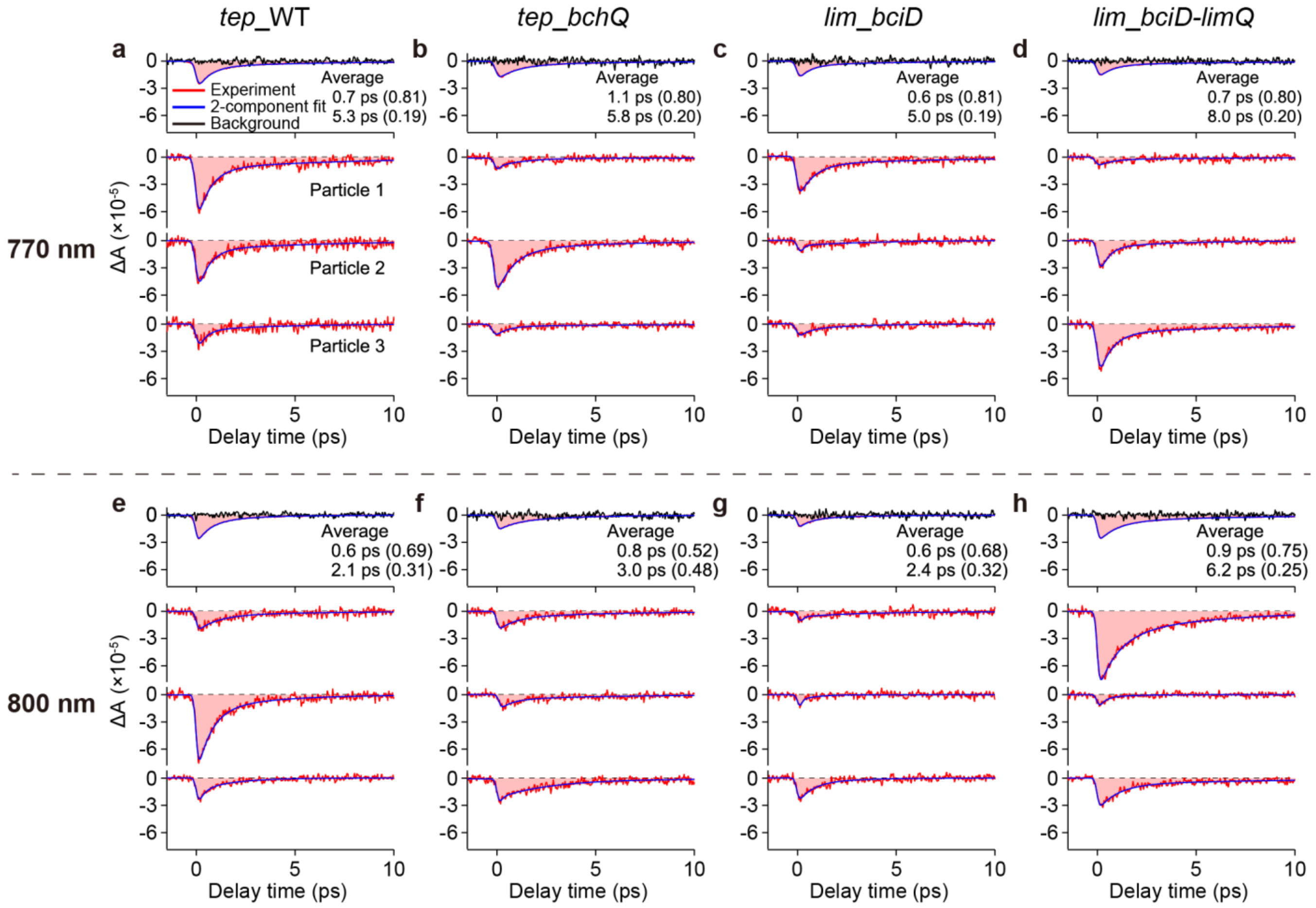

**Fig. S16 | TA signals from individual wild-type and mutant chlorosomes.**

(**a**–**h**) TA signals from individual chlorosomes of *tep*_WT (a, e), *tep_bchQ* (b, f), *lim_bciD* (c, g), and *lim_bciD-limQ* (d, h), measured using probes at 770 nm (a–d) and 800 nm (e–h). Each panel shows the averaged TA signal (top) and three representative traces from distinct individual chlorosomes (bottom) for the corresponding sample and probe wavelength. The averaged TA signals correspond to those in Fig. S15 but are shown here without intensity normalization. Biexponential fits (blue) and background signals (black) are overlaid. The time constants $\tau$ estimated from fits to the averaged TA signals, with the corresponding relative $\Delta A$ values in parentheses, are also shown at the top of each panel. The three TA signals from distinct individual *tep*_WT chlorosomes and their fitted curves shown in (a, bottom) are replotted in Fig. 1d with the vertical axis rescaled.

**Table S4**

Number of chlorosomes used for statistical analysis relative to the number of chlorosomes measured.

| | 770 nm | 800 nm |
|---|---|---|
| *tep*_WT | 194 / 200 | 162 / 180 |
| *tep_bchQ* | 148 / 204 | 111 / 204 |
| *lim_bciD* | 159 / 200 | 54 / 200 |
| *lim_bciD-limQ* | 124 / 200 | 140 / 201 |

#### Supplementary Note 7: Distributions of multiple parameters resolved by single-chlorosome spectroscopy

Distributions of time constants $\tau$, total $\Delta A$, and relative $\Delta A$

TA signals measured from individual wild-type and mutant chlorosomes were analyzed using the fitting procedure described in Supplementary Note 6, yielding two decay time constants $\tau_1$ and $\tau_2$ and their amplitudes $\Delta A_1$ and $\Delta A_2$. The distributions of the amplitude-weighted mean time constant $\langle\tau\rangle$ are shown as histograms with a bin size of 0.25 ps in Fig. S17a. The mean and median are indicated by a black dashed line and a red solid line, respectively. Importantly, conventional ensemble measurements would provide an ensemble-averaged value corresponding to this distribution rather than resolve the distribution itself. For *tep*_WT chlorosomes probed at 770 nm, the mean of the $\langle\tau\rangle$ distribution was 1.5 ps (Fig. S17a, upper left). With the 800-nm probe, it decreased to 1.1 ps, which is ~0.7 times the value at 770 nm (Fig. S17a, upper right). For mutant chlorosomes, the mean of the $\langle\tau\rangle$ distribution was in the range of 1.4–1.8 ps at 770 nm and 1.1–2.0 ps at 800 nm (Fig. S17a, the lower three on each side), remaining broadly similar to *tep*_WT. However, the widths and shapes of the distributions differed markedly among samples.

For a more detailed analysis of the distributions, $\tau_1$ and $\tau_2$ obtained from each chlorosome were summarized in a single histogram with a bin size of 0.25 ps (Fig. S17b and Fig. 2a–d, g–j). Here, decay components with relative amplitudes of <0.01, which were observed in 0.6% of the chlorosomes (9/1589), were excluded. Additionally, decay components with $\tau$ values exceeding the 10-ps measurement time window were not included in the histograms because the resulting estimates were less reliable (see Supplementary Note 8). The $\tau$ values were distributed over a broad range. Both the widths and the shapes of the distributions varied substantially among samples, indicating contributions from multiple kinetic components whose relative populations differed among samples.

In addition, a histogram of the total $\Delta A$ (= $\Delta A_1 + \Delta A_2$) was constructed with a bin size of $5 \times 10^{-6}$ (Fig. S18a). Histograms of relative $\Delta A_1$ and relative $\Delta A_2$, defined as $\Delta A_1$ / total $\Delta A$ and $\Delta A_2$ / total $\Delta A$, respectively, were constructed with a bin size of 0.03 (Fig. S18b).

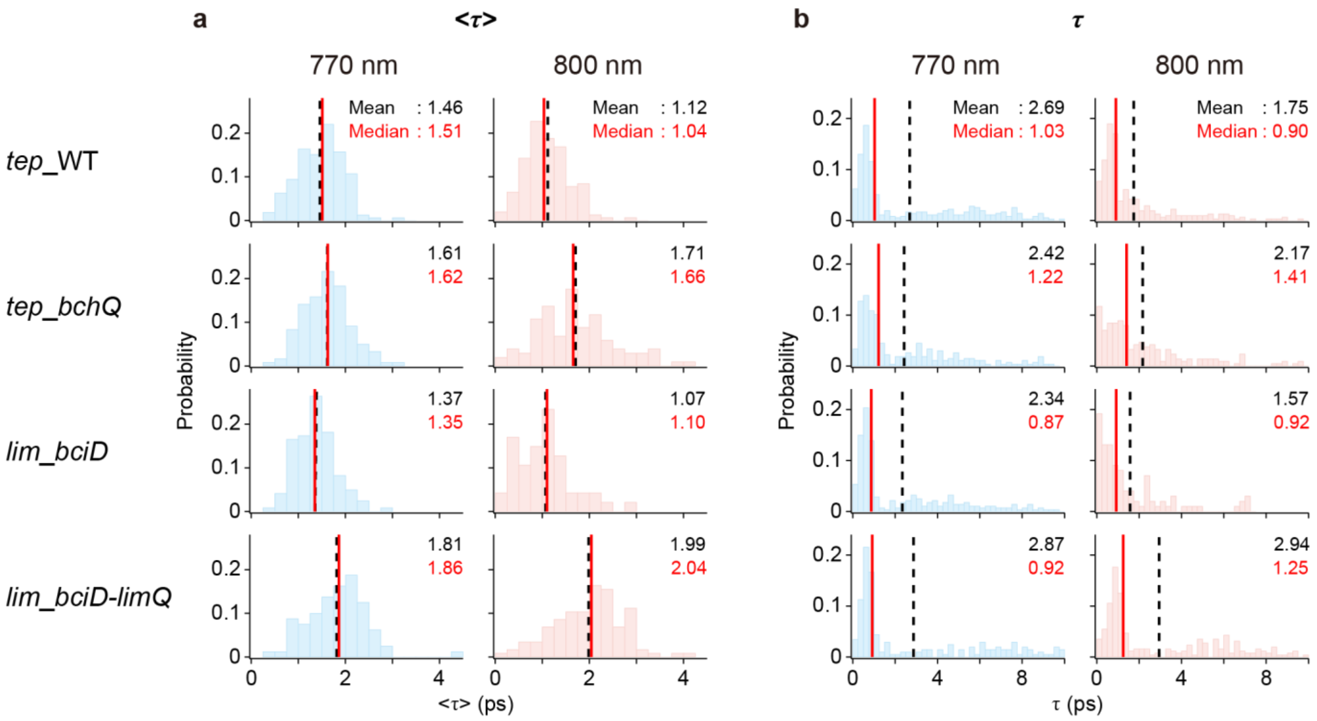

**Fig. S17 | Distributions of $\tau$.**

Distributions of (**a**) the amplitude-weighted mean time constant $\langle\tau\rangle$ and (**b**) $\tau$ values, including both $\tau_1$ and $\tau_2$, estimated from TA signals measured for *tep*_WT, *tep_bchQ*, *lim_bciD*, and *lim_bciD-limQ* chlorosomes (from top to bottom) using probes at 770 nm (left, blue) and 800 nm (right, pink). The bin size is 0.25 ps. The dashed black and solid red lines indicate the mean and median, respectively.

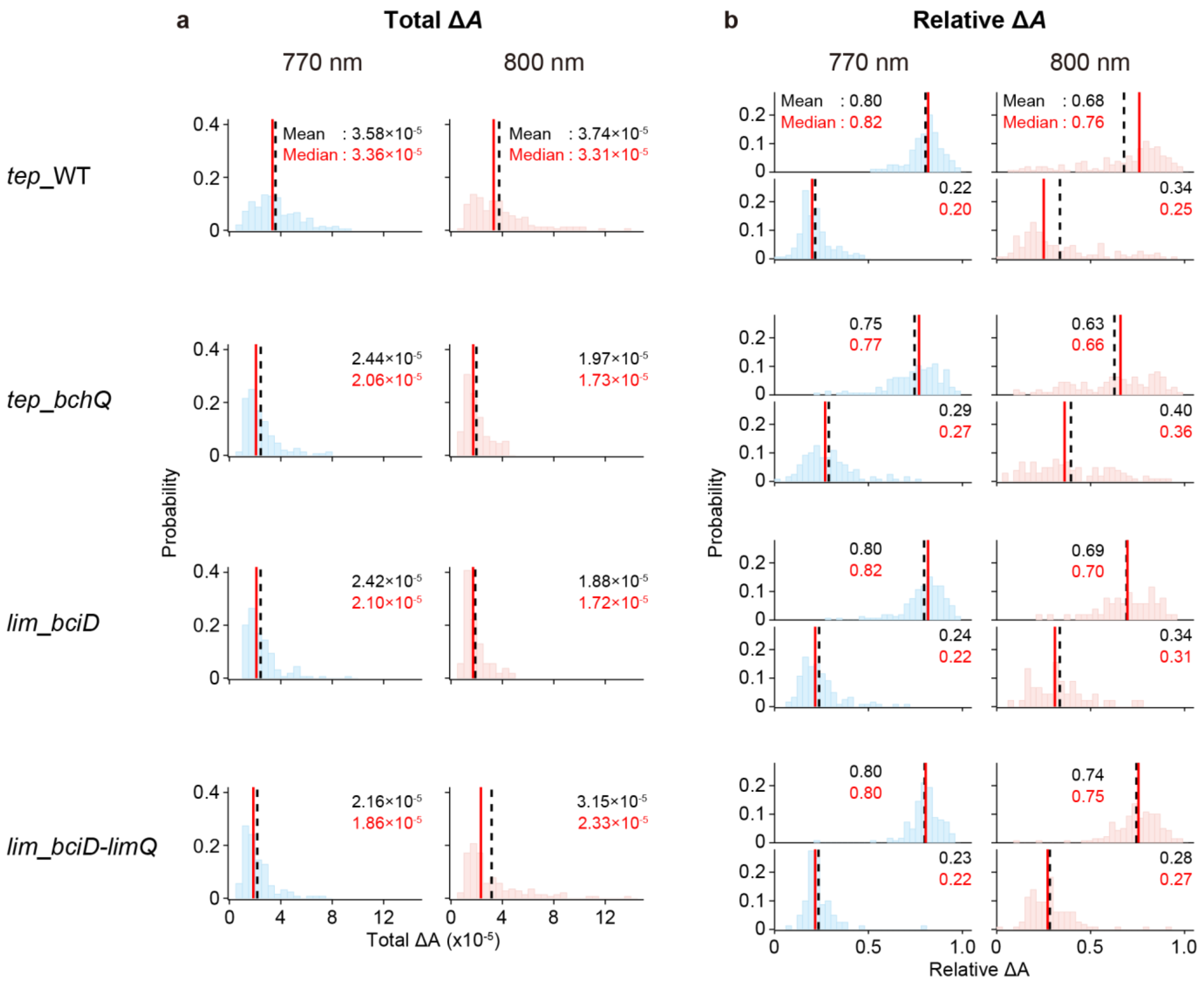

**Fig. S18 | Distributions of Δ*A*.**

Distributions of (**a**) total Δ*A* and (**b**) relative Δ*A* values, estimated from TA signals measured for *tep*_WT, *tep_bchQ*, *lim_bciD*, and *lim_bciD-limQ* chlorosomes (from top to bottom) using probes at 770 nm (left, blue) and 800 nm (right, pink). In (b), the upper and lower panels for each sample show the distributions of relative $\Delta A_1$ and relative $\Delta A_2$, respectively. The bin sizes are $5 \times 10^{-6}$ for total Δ*A* and 0.03 for relative Δ*A*. The dashed black and solid red lines indicate the mean and median, respectively.

Distributions of fluorescence intensity and effective EET efficiency

The fluorescence intensity of each chlorosome was calculated by averaging the signal within a $0.5 \times 0.5$ μm$^2$ region in the fluorescence image, centered at the TA measurement position. Histograms of fluorescence intensities constructed with a bin size of 0.3 kcps are shown in Fig. S19a. The fluorescence was detected at >805 nm, allowing emission from BChl *a* in the baseplate to be selectively observed (see Supplementary Note 1). Consequently, the fluorescence intensity

(Fig. S19a) reflects the amount of excitation energy transferred to the baseplate, whereas total $\Delta A$ (Fig. S18a) reflects the amount of excitation generated by light absorption in each chlorosome. Thus, dividing the fluorescence intensity by the total $\Delta A$ obtained at the corresponding probe wavelength yielded an effective EET efficiency to the baseplate for each chlorosome (Fig. S19b). For display, the efficiency values in each histogram were normalized to the median of the corresponding distribution.

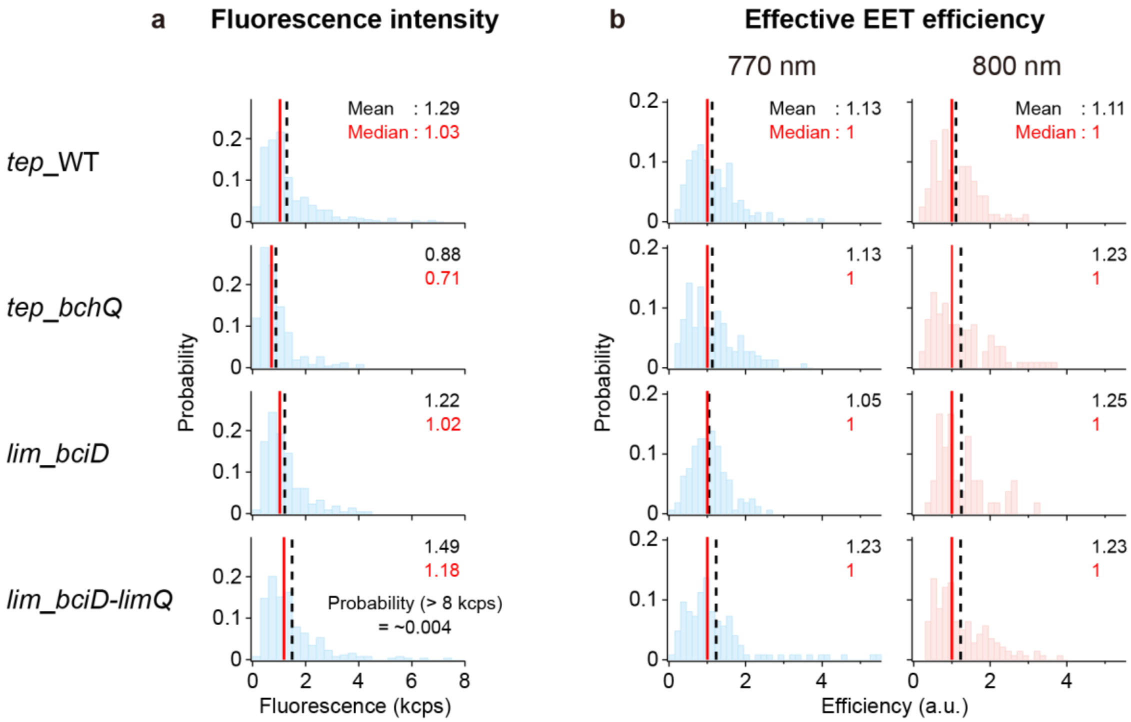

**Fig. S19 | Distributions of fluorescence intensity and effective EET efficiency.**

Distributions of (**a**) fluorescence intensity and (**b**) effective EET efficiency to the baseplate, obtained for *tep*_WT, *tep_bchQ*, *lim_bciD*, and *lim_bciD-limQ* chlorosomes (from top to bottom). Fluorescence intensities of individual chlorosomes were obtained from fluorescence images acquired under 740-nm excitation with emission detected at >805 nm. For *lim_bciD-limQ* chlorosomes, the probability of fluorescence intensities exceeding 8 kcps was ~0.004. Effective EET efficiencies were calculated by dividing these fluorescence intensities by total $\Delta A$ values obtained with probes at 770 nm (left, blue) and 800 nm (right, pink). The bin sizes are 0.3 kcps for fluorescence intensity and 0.15 for effective EET efficiency. The dashed black and solid red lines indicate the mean and median, respectively. The effective EET efficiency values were normalized to their respective medians such that each median equals 1.

#### 2D distributions of $\tau$ versus other parameters

As described above, in addition to the $\tau$ distribution (Fig. S17b), we obtained the distributions of total $\Delta A$ (Fig. S18a), relative $\Delta A$ (Fig. S18b), and effective EET efficiency (Fig. S19b), which enabled us to construct 2D histograms of $\tau$ versus total $\Delta A$, relative $\Delta A$, or effective EET efficiency.

In the 2D histogram of $\tau$ versus relative $\Delta A$, the relative $\Delta A_1$ and relative $\Delta A_2$ distributions were combined into a single distribution. By fitting these 2D histograms (see Supplementary Note 10), we estimated the mean values and distribution widths of total $\Delta A$, relative $\Delta A$, and effective EET efficiency for each $\tau$ component (Figs. S2–S4, S20–S22).

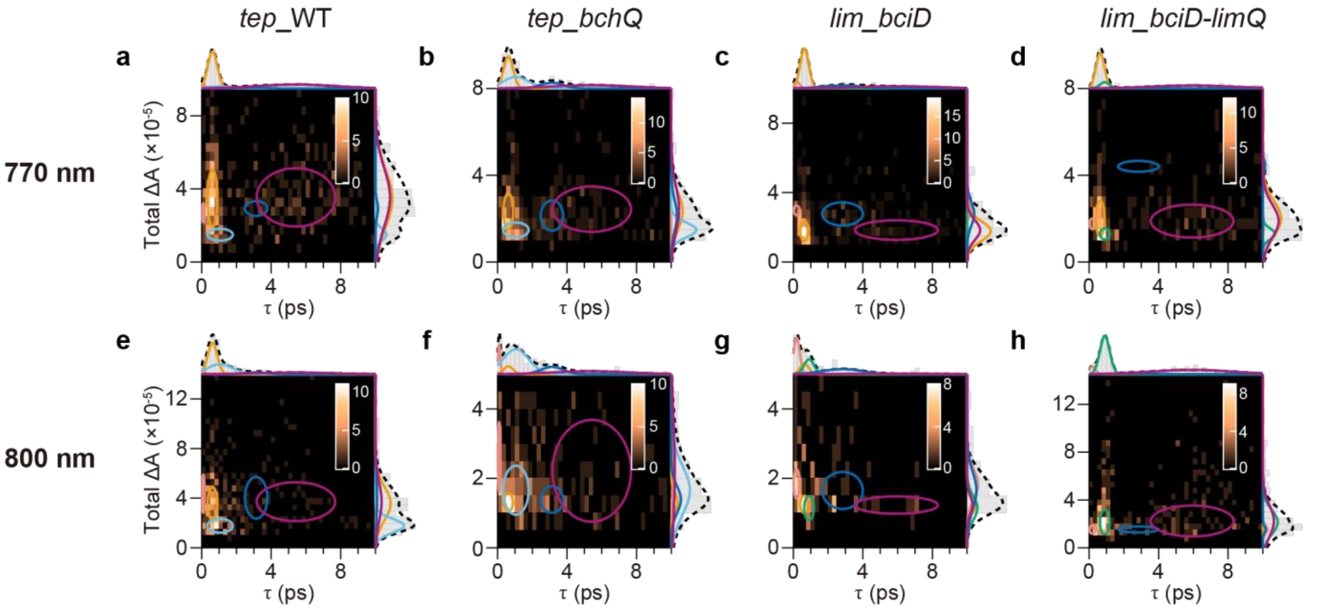

**Fig. S20 | Estimation of total $\Delta A$ associated with each $\tau$ component.**

(**a**–**d**) 2D distributions of total $\Delta A$ versus $\tau$ for *tep*_WT (a), *tep_bchQ* (b), *lim_bciD* (c), and *lim_bciD-limQ* (d) chlorosomes, obtained using the 770-nm probe. The bin sizes are $5 \times 10^{-6}$ for total $\Delta A$ and 0.25 ps for $\tau$. The circles denote contours at $1/\sqrt{e}$ of the peak height (i.e., one-standard-deviation contours) for each $\tau$ component, estimated from five-component 2D Gaussian fits. The projections of the 2D distributions onto the *x*- and *y*-axes are shown as gray histograms at the top and right of each panel, respectively. The fitted distributions associated with each $\tau$ component are represented by solid lines colored as in Fig. 2, and their sum is indicated by the dashed black line. (**e**–**h**) Corresponding results obtained using the 800-nm probe. The fitted total $\Delta A$ distributions are summarized in Fig. S2.

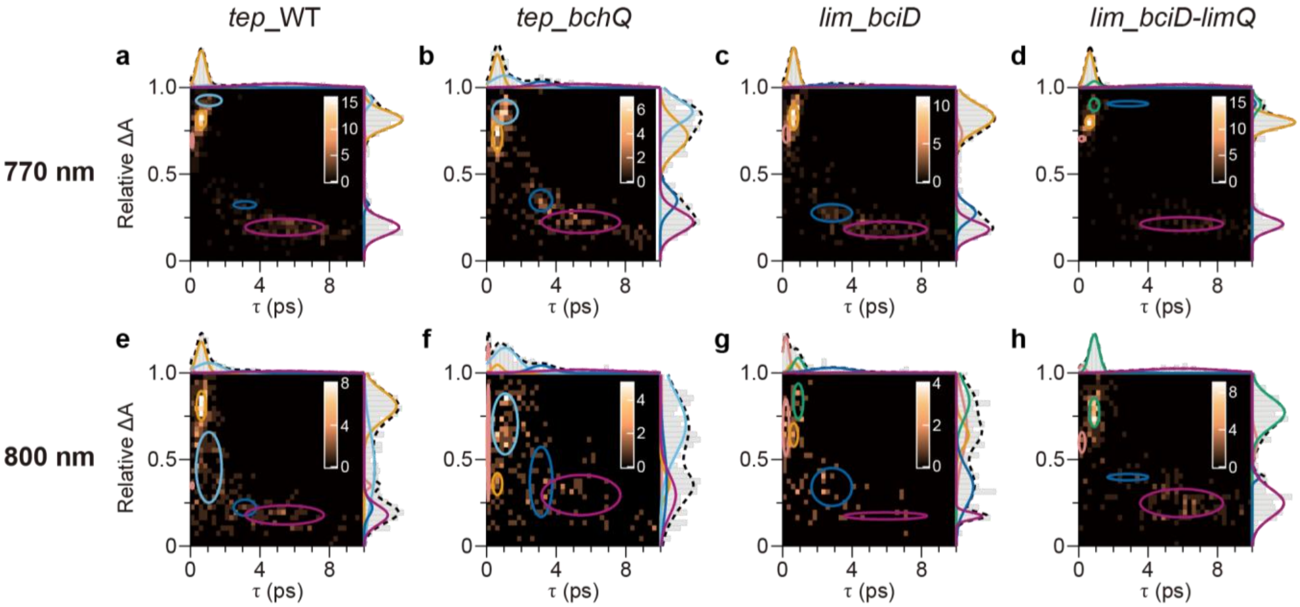

**Fig. S21 | Estimation of relative $\Delta A$ associated with each $\tau$ component.**

2D distributions of relative $\Delta A$ versus $\tau$ for each sample and probe wavelength. The bin sizes are 0.03 for relative $\Delta A$ and 0.25 ps for $\tau$. The plots are presented in the same format as in Fig. S20. The fitted relative $\Delta A$ distributions are summarized in Fig. S3.

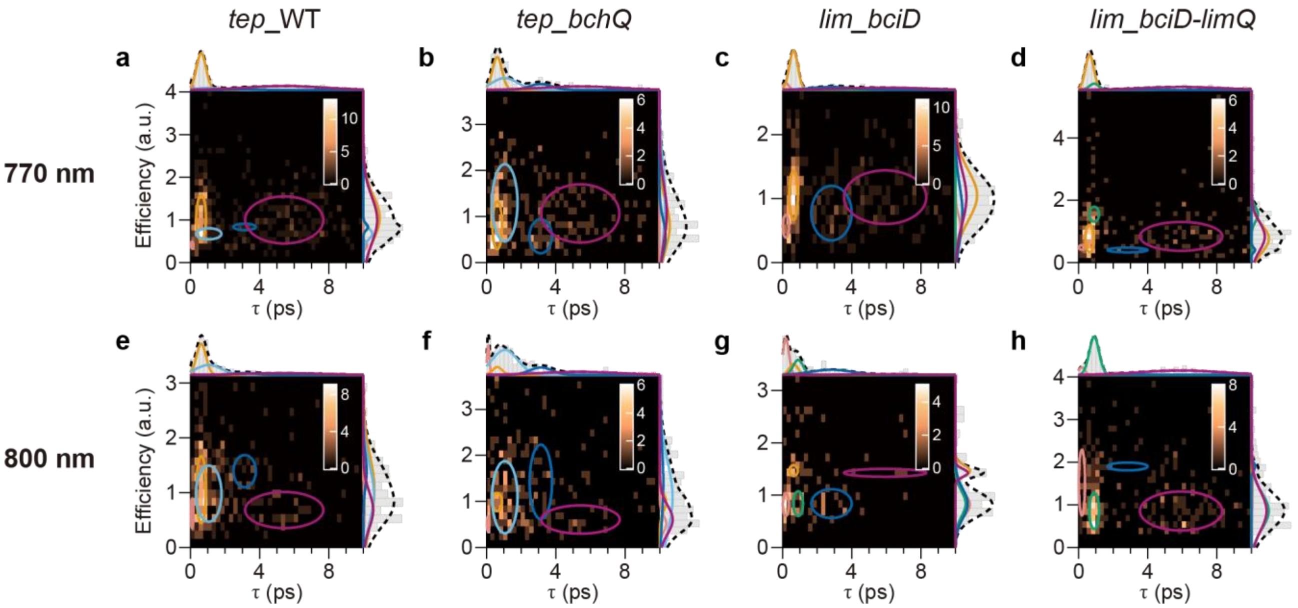

**Fig. S22 | Estimation of effective EET efficiency associated with each $\tau$ component.**

2D distributions of effective EET efficiency to the baseplate versus $\tau$ for each sample and probe wavelength. The effective EET efficiency values were normalized to the median of the corresponding overall distribution. The bin sizes are 0.15 for effective EET efficiency and 0.25 ps for $\tau$. The plots are presented in the same format as in Fig. S20. The fitted effective EET efficiency distributions are summarized in Fig. S4.

#### Supplementary Note 8: Reliability of time constant estimation relative to the TA measurement time window

The temporal window of the single-chlorosome TA measurements was 10 ps. In all samples, time constants longer than 10 ps were estimated by exponential fits to TA decay traces. To evaluate the reliability of such long $\tau$ estimates, we performed simulations as follows. First, we generated single-exponential decay traces over 10 ps (Fig. S23a, orange), with $\tau$ set to values between 5 and 40 ps and the SNR fixed at 6.8, which corresponds to the median SNR obtained from the experimental data. The FWHM of the IRF was fixed at 313 fs (Table S3b). Then, we fitted the simulated traces with a single-exponential function to estimate $\tau$ (Fig. S23a, blue). The standard deviation of the estimated $\tau$ values was evaluated from 1,000 trials of simulation and fitting. By dividing the standard deviation by the set $\tau$, we quantified the relative fitting error, which was plotted as a function of the set $\tau$ over the range of 5–40 ps (Fig. S23c, orange). The relative error remained below 0.06 up to 10 ps, whereas it increased with $\tau$ above 10 ps. This indicates that the fitting of components with $\tau$ longer than the measurement window is less reliable. Similarly, we evaluated the fitting error at an SNR of 3.0, corresponding to the lower-limit SNR cutoff used in our analysis of the experimental data (Fig. S23b). The fitting reliability decreased compared with that at an SNR of 6.8. Nevertheless, the relative error was still below 0.12 up to 10 ps, whereas it rose for $\tau$ above 10 ps (Fig. S23c, pink). Accordingly, the $\tau$ distributions (Fig. 2a–d, g–j) were constructed using only components with $\tau$ not exceeding 10 ps. The origin of the >10-ps component is discussed in Supplementary Note 11.

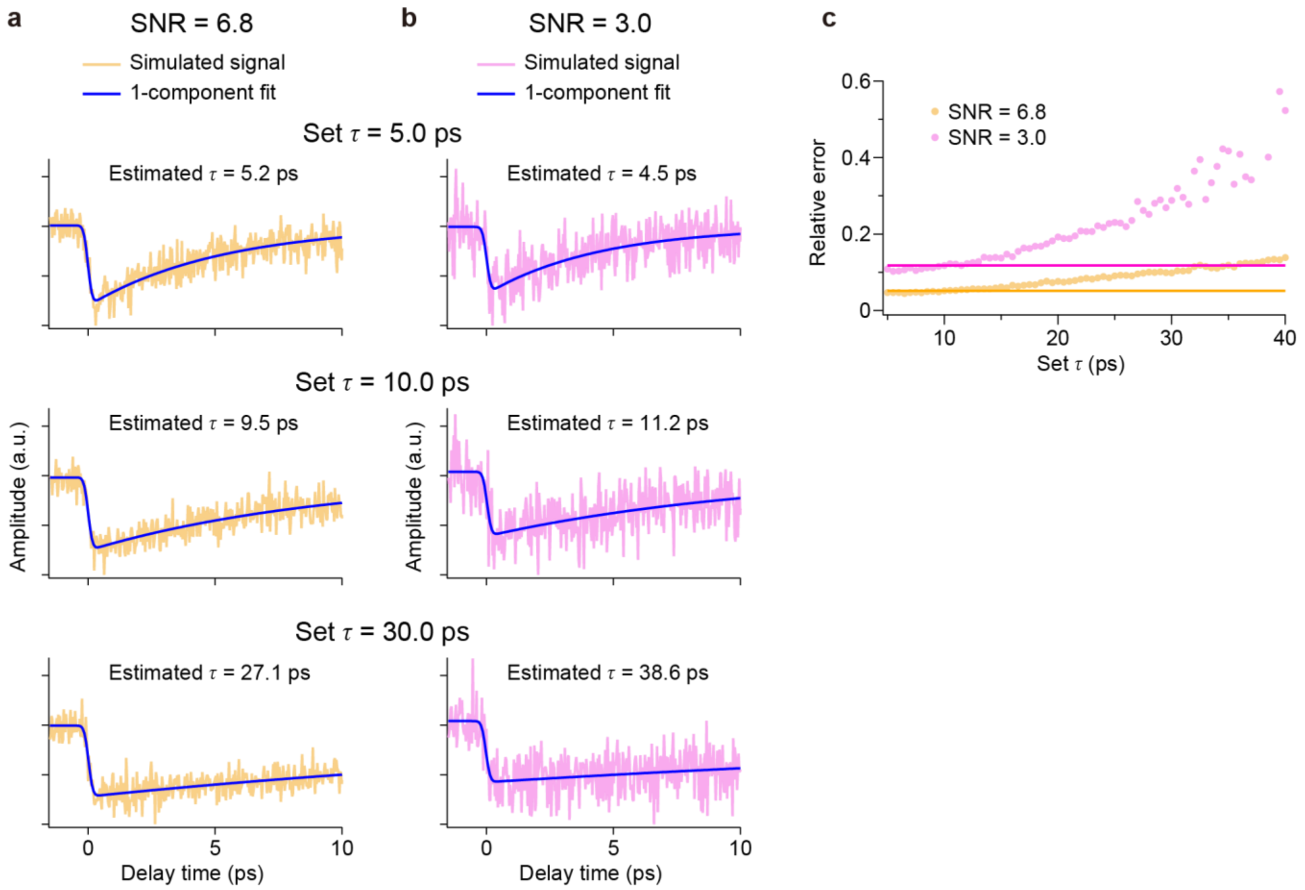

**Fig. S23 | Assessment of the reliability of time-constant estimates.**

(**a**, **b**) Simulated TA traces generated using a single-exponential function with set $\tau$ values of 5.0, 10.0, and 30.0 ps (from top to bottom) over a 10-ps time window at SNRs of 6.8 (a) and 3.0 (b). The fitted curves are indicated by blue lines. Estimated $\tau$ values are given in each panel. (**c**) Relative fitting errors of the estimated $\tau$ values at SNRs of 6.8 (orange) and 3.0 (pink), plotted as a function of the set $\tau$ values. The relative fitting errors for the set $\tau$ value of 10.0 ps, which matches the 10-ps time window, are indicated by horizontal lines for each SNR condition.

#### Supplementary Note 9: $\tau$-distribution analysis

<u>Decomposition of the $\tau$ distribution by global fitting with multiple Gaussian functions</u>

As shown in Fig. 2 and Fig. S17b, the histograms of the time constant $\tau$ appeared to comprise multiple components. To identify them, we performed fits using the following function, which represents a sum of Gaussian components:

$$\mathrm{f}(\tau) = \sum_{i=1}^{N} A_i \exp\left(-\frac{1}{2}\left(\frac{\tau - \tau_{0i}}{\tau_{wi}}\right)^2\right). \quad (3)$$

Here, $N$ is the number of components, and $A_i$, $\tau_{0i}$, and $\tau_{wi}$ denote, respectively, the amplitude, center position, and $1/\sqrt{e}$ half-width of the Gaussian distribution associated with component $i$. The FWHM is obtained by multiplying the $1/\sqrt{e}$ half-width by the factor $2\sqrt{2\ ln2}$.

The four histograms in Fig. 2a–d, obtained with the 770-nm and 800-nm probes for wild-type and mutant chlorosomes from *C. tepidum* (*tep*_WT and *tep_bchQ*), were globally fitted with $N$ ranging from 4 to 6. The result of the four-component Gaussian fit is shown in Fig. S24a. In the <1-ps region, the distribution was reproduced either by two components with nearly identical centers but different widths or by a single broad component, yielding an overly coarse fit. Moreover, the distribution showed a pronounced discontinuity at 0 ps. Thus, four components were insufficient. When the number of components was increased to five, RSS decreased to 85% of the value for the four-component fit, and the <1-ps region was well reproduced (Fig. S24b). A six-component fit also reduced RSS to 69% relative to the four-component fit, but it yielded two components with nearly identical time constants of 0.6 and 0.7 ps, separated by less than the histogram bin size of 0.25 ps. In addition, several components with $\tau < 2$ ps exhibited negligible populations (Fig. S24c). Furthermore, the fitted values varied substantially with the binning conditions, as described below. These results indicate that a six-component model was overparameterized and could produce artifactual components. We therefore concluded that the $\tau$ distributions for *tep*_WT and *tep_bchQ* were adequately described by five components (Fig. S24b and Fig. 2a–d). The populations of the $\tau$ components are summarized in Fig. 2e, f, and Table S5a.

A similar analysis was conducted for chlorosomes from *C. limnaeum* mutants (*lim_bciD* and *lim_bciD-limQ*). A global fit with four components failed to reproduce the <2-ps region of the distribution for *lim_bciD* probed at 800 nm (Fig. S24d). When the number of components was increased to five, RSS decreased to 59% of the value for the four-component fit, and the <2-ps region was satisfactorily reproduced (Fig. S24e). Further increasing the number of components to six decreased RSS only modestly, from 59% to 55% of the value for the four-component fit (Fig. S24f). At the same time, the instability with respect to the binning conditions increased, as described below. Accordingly, as with *tep*_WT and *tep_bchQ*, we concluded that five components were sufficient to reproduce the $\tau$ distributions of *lim_bciD* and *lim_bciD-limQ* (Fig. S24e and Fig. 2g–j). The populations of the $\tau$ components are summarized in Fig. 2k, l, and Table S5b.

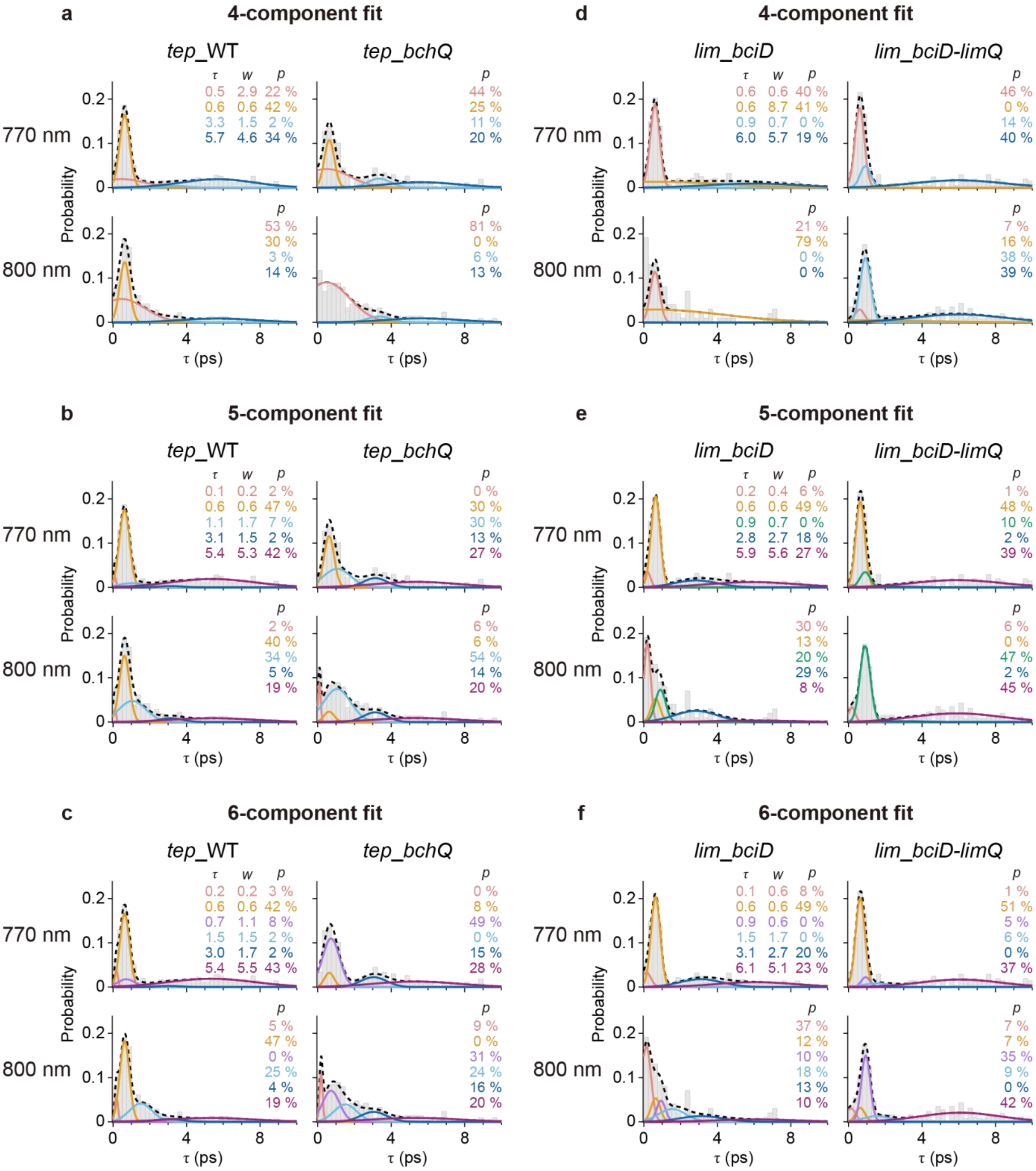

**Fig. S24 | Identification of multiple $\tau$ components using Gaussian fitting.**

(**a**–**c**) Global fits with four (a), five (b), and six (c) Gaussian components to four $\tau$ distributions obtained for *tep*_WT (left) and *tep_bchQ* (right) chlorosomes using probes at 770 nm (top) and 800 nm (bottom). Fitted curves for the $\tau$ components (solid lines in distinct colors), together with their sum (dashed black line), are overlaid on the $\tau$ distributions. The center positions ($\tau$) and FWHMs ($w$) for each $\tau$ component, which were estimated as global parameters shared among the four $\tau$ distributions, are shown in the upper-left subplot of each panel. The populations ($p$) for each $\tau$ component, which were estimated as local parameters specific to each distribution, are also shown in each subplot. These fitted parameters are indicated in the same colors as the fitted curves. (**d**–**f**) Corresponding global fits to the four $\tau$ distributions obtained for *lim_bciD* (left) and *lim_bciD-limQ* (right) chlorosomes using probes at 770 nm (top) and 800 nm (bottom). The bin size for the $\tau$ distributions is 0.25 ps. The $\tau$ distributions and their five-component fits for *tep*_WT and *tep_bchQ* chlorosomes in (b) and for *lim_bciD* and *lim_bciD-limQ* chlorosomes in (e) are replotted in Fig. 2a–d and Fig. 2g–j, respectively, with expanded views of the 0–2 ps region also shown.

**Table S5**

Center position, FWHM, and population for each $\tau$ component estimated from five-component global Gaussian fits to the $\tau$ distributions obtained for (**a**) *tep*_WT and *tep_bchQ* chlorosomes and (**b**) *lim_bciD* and *lim_bciD-limQ* chlorosomes using probes at 770 nm and 800 nm. The values are shown as fitted values ± standard deviations, where the standard deviations were estimated from five resampling trials of the $\tau$-distribution analysis. For populations, standard deviations smaller than 0.1 are indicated as <0.1.

| **a** | Center position (ps) | FWHM (ps) | Population (%) | | | |
|---|---|---|---|---|---|---|
| | | | *tep*_WT | | *tep_bchQ* | |
| | | | 770 nm | 800 nm | 770 nm | 800 nm |
| c1 | 0.09 ± 0.02 | 0.16 ± 0.04 | 1.7 ± 0.7 | 1.9 ± 1.2 | 0.0 ± <0.1 | 6.0 ± 1.7 |
| c2 | 0.62 ± 0.03 | 0.65 ± 0.07 | 47.6 ± 3.2 | 39.9 ± 5.5 | 30.7 ± 5.8 | 6.4 ± 4.8 |
| c3 | - | - | - | - | - | - |
| c4 | 1.06 ± 0.15 | 1.73 ± 0.23 | 6.9 ± 2.8 | 33.8 ± 6.2 | 29.7 ± 4.8 | 53.8 ± 9.8 |
| c5 | 3.13 ± 0.19 | 1.52 ± 0.23 | 1.9 ± 2.8 | 5.4 ± 2.7 | 13.0 ± 3.7 | 13.9 ± 5.0 |
| c6 | 5.42 ± 0.36 | 5.32 ± 0.86 | 41.9 ± 4.6 | 18.9 ± 5.8 | 26.6 ± 6.3 | 19.9 ± 3.8 |

| **b** | Center position (ps) | FWHM (ps) | Population (%) | | | |
|---|---|---|---|---|---|---|
| | | | *lim_bciD* | | *lim_bciD-limQ* | |
| | | | 770 nm | 800 nm | 770 nm | 800 nm |
| c1 | 0.19 ± 0.06 | 0.41 ± 0.05 | 5.9 ± 5.3 | 30.5 ± 1.4 | 1.0 ± 3.1 | 5.7 ± 1.2 |
| c2 | 0.64 ± 0.03 | 0.58 ± 0.05 | 49.2 ± 4.2 | 12.7 ± 6.4 | 48.2 ± 4.2 | 0.0 ± 7.3 |
| c3 | 0.90 ± 0.06 | 0.67 ± 0.09 | 0.0 ± <0.1 | 20.2 ± 9.1 | 9.8 ± 5.3 | 47.0 ± 6.5 |
| c4 | - | - | - | - | - | - |
| c5 | 2.83 ± 0.30 | 2.75 ± 0.20 | 17.6 ± 4.7 | 28.6 ± 5.5 | 1.7 ± 3.9 | 2.2 ± 2.4 |
| c6 | 5.93 ± 0.30 | 5.61 ± 1.01 | 27.2 ± 6.1 | 7.9 ± 4.0 | 39.3 ± 5.5 | 45.1 ± 3.4 |

Robustness of the $\tau$-distribution analysis to histogram binning conditions

The $\tau$-distribution profile can vary with histogram bin size, potentially affecting the results obtained from the $\tau$-distribution analysis. To assess robustness to binning conditions, we performed five-component global fits to histograms constructed using bin sizes that were varied by ±20% relative to 0.25 ps. These distinct bin sizes led to slightly different fitting results for the histograms obtained with the 770-nm and 800-nm probes for *tep*_WT and *tep_bchQ* (Fig. S25a–c) and those for *lim_bciD* and *lim_bciD-limQ* (Fig. S25d–f). From each data set, the means and standard deviations of the fitted values across these binning conditions were calculated for each $\tau$ component (Table S6a and b, respectively). For the fitted center positions and FWHMs, the standard deviations were generally smaller than the bin size of 0.25 ps, and even in the case with the largest standard deviation, the CV defined as the standard deviation divided by the mean was only 4.9%. Similarly, for the $\tau$-component populations, the standard deviations were at most 5.0%, corresponding to a CV of 10.7%. These results indicate that the $\tau$-distribution analysis was robust with respect to histogram bin size.

By contrast, the six-component fits were less stable with respect to the binning conditions (Fig. S26 and Table S7). The mean (median) CV of the $\tau$-component populations, calculated for $\tau$ components with populations of at least 1%, was 34.2% (20.3%), representing ~1.7-fold (~2.2-fold) increases relative to the corresponding values obtained from the five-component fit. In particular, for *tep*_WT probed at 800 nm (Fig. S26a–c, lower left), *tep_bchQ* probed at 770 nm (Fig. S26a–c, upper right), and *lim_bciD* probed at 800 nm (Fig. S26d–f, lower left), the relative ordering of the $\tau$-component populations changed markedly depending on the binning conditions. These results suggest that the six-component model was overparameterized and prone to overfitting, supporting the conclusion that a five-component model is sufficient for the present $\tau$-distribution analysis.

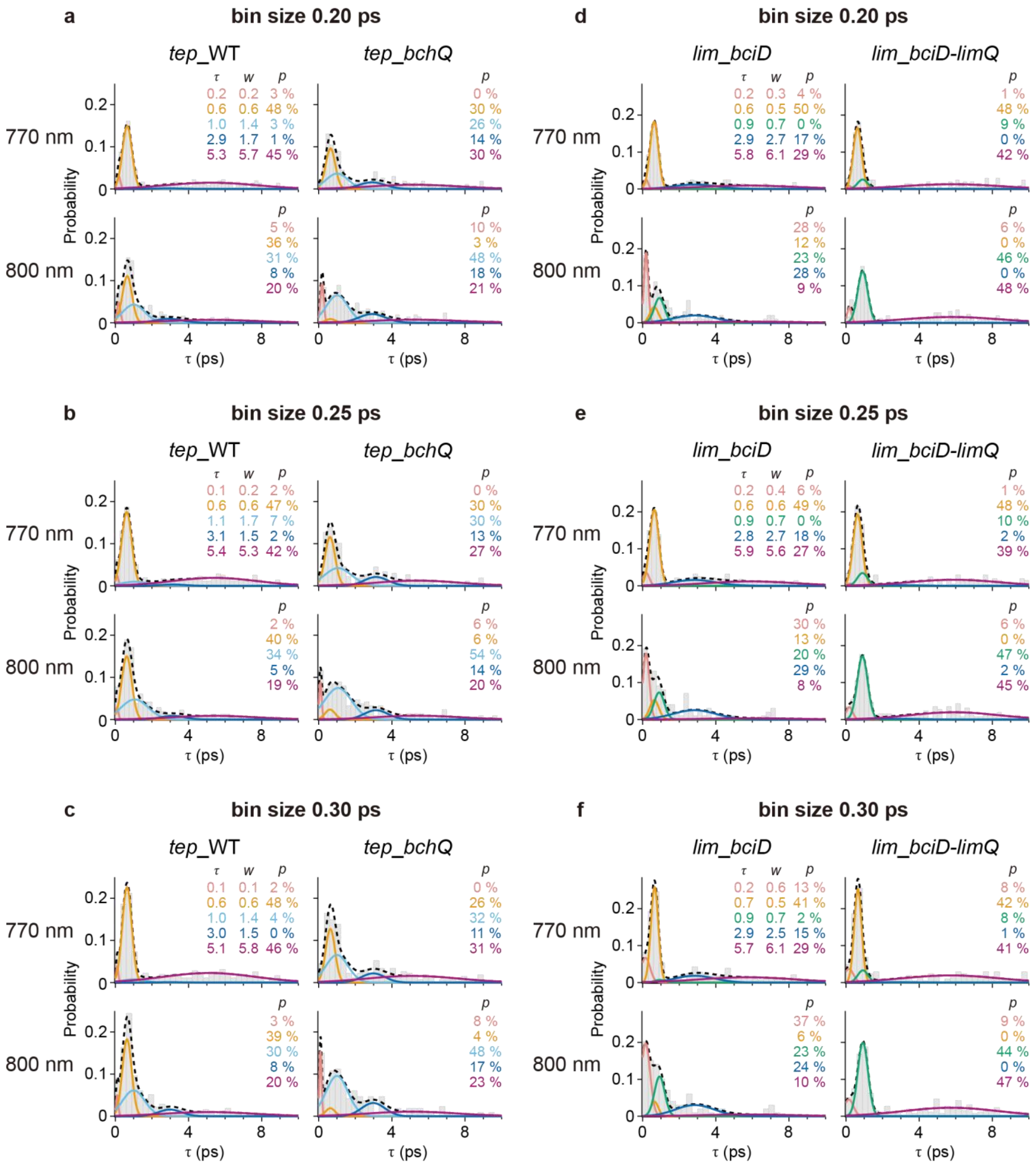

**Fig. S25 | Five-component $\tau$-distribution analysis with bin sizes varied by ±20%.**

(**a**–**c**) Five-component global fits to four $\tau$ distributions constructed with bin sizes varied by −20% (a), 0% (b), and +20% (c), corresponding to bin sizes of 0.20, 0.25, and 0.30 ps, respectively. The $\tau$ distributions were obtained for *tep*_WT (left) and *tep_bchQ* (right) chlorosomes using probes at

770 nm (top) and 800 nm (bottom). (**d**–**f**) Corresponding global fits to the four $\tau$ distributions constructed with the same bin-size variations, obtained for *lim_bciD* (left) and *lim_bciD-limQ* (right) chlorosomes using probes at 770 nm (top) and 800 nm (bottom). The fitted curves and the estimated parameters, including the center position ($\tau$), FWHM ($w$), and population ($p$) for each $\tau$ component, are shown in the same format as in Fig. S24.

**Table S6**

Parameters from five-component global Gaussian fits to $\tau$ distributions under different binning conditions. Center position, FWHM, and population for each $\tau$ component were estimated from the fits to the $\tau$ distributions constructed with three distinct binning conditions (−20%, 0%, and +20%) for (**a**) *tep*_WT and *tep_bchQ* chlorosomes and (**b**) *lim_bciD* and *lim_bciD-limQ* chlorosomes measured with probes at 770 nm and 800 nm. The values are shown as means ± standard deviations across the three binning conditions. For FWHMs and populations, standard deviations smaller than 0.01 and 0.1 are indicated as <0.01 and <0.1, respectively.

| **a** | Center position (ps) | FWHM (ps) | Population (%) | | | |
|---|---|---|---|---|---|---|
| | | | *tep*_WT | | *tep_bchQ* | |
| | | | 770 nm | 800 nm | 770 nm | 800 nm |
| c1 | 0.13 ± 0.05 | 0.17 ± 0.03 | 2.4 ± 0.9 | 3.3 ± 1.8 | 0.0 ± <0.1 | 8.1 ± 2.2 |
| c2 | 0.63 ± 0.01 | 0.62 ± 0.02 | 47.7 ± 0.1 | 37.9 ± 2.4 | 29.1 ± 2.6 | 4.4 ± 1.8 |
| c3 | - | - | - | - | - | - |
| c4 | 1.02 ± 0.03 | 1.51 ± 0.18 | 4.4 ± 2.1 | 31.8 ± 1.8 | 29.2 ± 2.6 | 49.9 ± 3.4 |
| c5 | 3.01 ± 0.11 | 1.57 ± 0.11 | 1.2 ± 0.7 | 7.3 ± 1.6 | 12.6 ± 1.4 | 16.3 ± 2.2 |
| c6 | 5.28 ± 0.14 | 5.63 ± 0.27 | 44.3 ± 2.1 | 19.7 ± 0.7 | 29.1 ± 2.3 | 21.2 ± 1.3 |

| **b** | Center position (ps) | FWHM (ps) | Population (%) | | | |
|---|---|---|---|---|---|---|
| | | | *lim_bciD* | | *lim_bciD-limQ* | |
| | | | 770 nm | 800 nm | 770 nm | 800 nm |
| c1 | 0.18 ± 0.02 | 0.42 ± 0.15 | 7.5 ± 4.9 | 31.7 ± 4.8 | 3.2 ± 3.7 | 6.8 ± 1.7 |
| c2 | 0.64 ± 0.01 | 0.53 ± 0.05 | 47.0 ± 5.0 | 10.3 ± 3.7 | 46.1 ± 3.1 | 0.0 ± <0.1 |
| c3 | 0.91 ± 0.01 | 0.66 ± <0.01 | 0.5 ± 0.9 | 22.1 ± 1.6 | 8.8 ± 1.1 | 45.7 ± 1.5 |
| c4 | - | - | - | - | - | - |
| c5 | 2.85 ± 0.02 | 2.66 ± 0.14 | 16.6 ± 1.2 | 27.2 ± 2.4 | 0.8 ± 0.9 | 0.7 ± 1.3 |
| c6 | 5.82 ± 0.10 | 5.95 ± 0.29 | 28.4 ± 1.0 | 8.7 ± 0.8 | 41.0 ± 1.6 | 46.8 ± 1.6 |

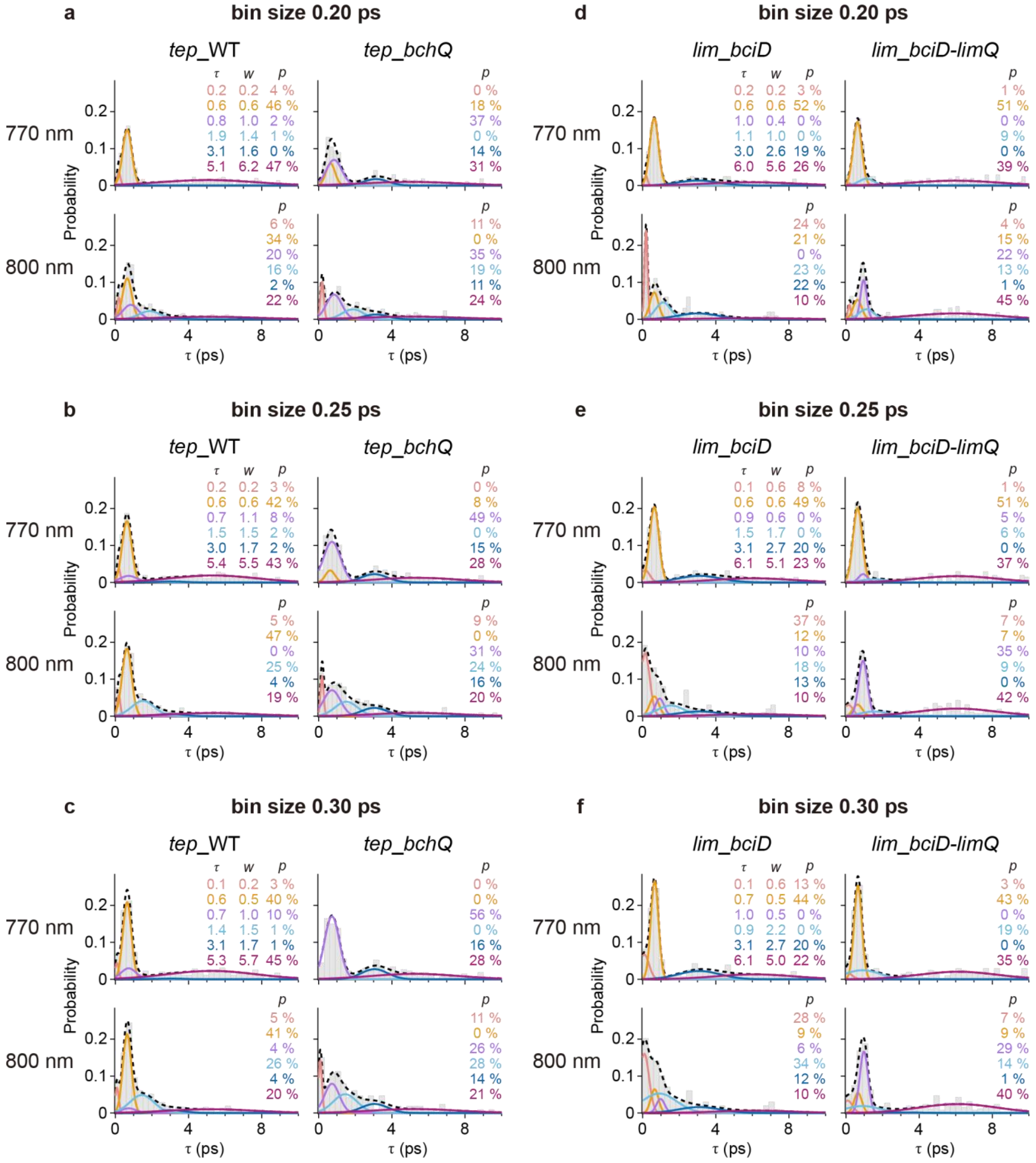

**Fig. S26 | Six-component $\tau$-distribution analysis with bin sizes varied by ±20%.**

(**a**–**c**) Six-component global fits to four $\tau$ distributions constructed with bin sizes varied by −20% (a), 0% (b), and +20% (c), corresponding to bin sizes of 0.20, 0.25, and 0.30 ps, respectively. The $\tau$ distributions were obtained for *tep*_WT (left) and *tep_bchQ* (right) chlorosomes using probes at

770 nm (top) and 800 nm (bottom). (**d**–**f**) Corresponding global fits to the four $\tau$ distributions constructed with the same bin-size variations, obtained for *lim_bciD* (left) and *lim_bciD-limQ* (right) chlorosomes using probes at 770 nm (top) and 800 nm (bottom). The fitted curves and the estimated parameters, including the center position ($\tau$), FWHM ($w$), and population ($p$) for each $\tau$ component, are shown in the same format as in Fig. S24.

**Table S7**

Parameters from six-component global Gaussian fits to $\tau$ distributions under different binning conditions. Center position, FWHM, and population for each $\tau$ component were estimated from the fits to the $\tau$ distributions constructed with three distinct binning conditions (−20%, 0%, and +20%) for (**a**) *tep*_WT and *tep_bchQ* chlorosomes and (**b**) *lim_bciD* and *lim_bciD-limQ* chlorosomes measured with probes at 770 nm and 800 nm. The six $\tau$ components are numbered 1–6 in order of increasing time constant. The values are shown as means ± standard deviations across the three binning conditions. For center positions and populations, standard deviations smaller than 0.01 and 0.1 are indicated as <0.01 and <0.1, respectively.

| a | Center position (ps) | FWHM (ps) | Population (%) | | | |
|---|---|---|---|---|---|---|
| | | | *tep*_WT | | *tep_bchQ* | |
| | | | 770 nm | 800 nm | 770 nm | 800 nm |
| 1 | 0.15 ± 0.06 | 0.21 ± 0.01 | 3.5 ± 0.4 | 5.6 ± 0.8 | 0.0 ± <0.1 | 10.5 ± 1.3 |
| 2 | 0.64 ± <0.01 | 0.58 ± 0.03 | 42.8 ± 3.2 | 40.2 ± 6.4 | 8.8 ± 9.2 | 0.0 ± <0.1 |
| 3 | 0.76 ± 0.07 | 1.01 ± 0.07 | 6.5 ± 4.4 | 8.1 ± 10.6 | 47.0 ± 10.1 | 31.1 ± 4.4 |
| 4 | 1.61 ± 0.23 | 1.45 ± 0.09 | 1.2 ± 0.6 | 22.2 ± 5.6 | 0.0 ± <0.1 | 23.4 ± 3.9 |
| 5 | 3.08 ± 0.05 | 1.64 ± 0.07 | 0.9 ± 0.9 | 3.6 ± 1.4 | 15.1 ± 0.8 | 13.5 ± 2.6 |
| 6 | 5.23 ± 0.15 | 5.76 ± 0.37 | 45.0 ± 2.5 | 20.4 ± 1.4 | 29.2 ± 2.0 | 21.4 ± 1.9 |

| b | Center position (ps) | FWHM (ps) | Population (%) | | | |
|---|---|---|---|---|---|---|
| | | | *lim_bciD* | | *lim_bciD-limQ* | |
| | | | 770 nm | 800 nm | 770 nm | 800 nm |
| 1 | 0.14 ± 0.05 | 0.47 ± 0.23 | 8.1 ± 5.4 | 30.2 ± 6.5 | 1.5 ± 1.6 | 6.0 ± 1.8 |
| 2 | 0.64 ± 0.01 | 0.55 ± 0.05 | 48.8 ± 4.3 | 14.0 ± 6.2 | 48.3 ± 4.5 | 10.5 ± 4.0 |
| 3 | 0.96 ± 0.02 | 0.50 ± 0.09 | 0.1 ± 0.1 | 5.3 ± 5.0 | 1.8 ± 3.1 | 28.4 ± 6.5 |
| 4 | 1.19 ± 0.32 | 1.64 ± 0.64 | 0.0 ± <0.1 | 25.2 ± 8.3 | 11.4 ± 6.5 | 12.1 ± 2.8 |
| 5 | 3.07 ± 0.04 | 2.66 ± 0.04 | 19.4 ± 0.7 | 15.6 ± 5.2 | 0.0 ± <0.1 | 0.6 ± 0.6 |
| 6 | 6.06 ± 0.08 | 5.21 ± 0.30 | 23.6 ± 2.0 | 9.8 ± 0.2 | 36.9 ± 2.1 | 42.4 ± 2.6 |

Estimation of uncertainties in the $\tau$-distribution analysis

Uncertainties in the $\tau$-distribution analysis were estimated from repeated resampling of the data as follows. For each sample and probe wavelength, we first randomly selected chlorosomes from the analyzed data set with replacement, allowing each chlorosome to be selected at most twice, until the number of selected chlorosomes matched the original count. The time constants for these resampled chlorosomes were used to construct histograms. The resulting distributions were then subjected to the five-component global fitting as described above to obtain the center position, width, and population of each $\tau$ component. In addition, the population of the >10-ps component was calculated by dividing the number of >10-ps time constants by the total number of estimated time constants. Differences in these $\tau$-component populations were calculated between samples (*tep_bchQ* minus *tep*_WT and *lim_bciD* minus *lim_bciD-limQ*) and between probe wavelengths (800 nm minus 770 nm). This entire procedure was repeated five times. The uncertainty associated with each estimated value was determined by calculating the standard deviation across the five trials (Table S5). The uncertainties in the populations are shown as error bars in Fig. 2e, f, k, and l, and in Fig. S29a, b. The uncertainties in the population differences are also shown as error bars in Fig. 2m–p and in Fig. S29c–f.

#### Supplementary Note 10: 2D distribution analysis

2D distribution fitting with multiple Gaussian distributions

From the single-chlorosome TA analysis, six $\tau$ components were identified (see the main text). The time constants and distribution widths for each $\tau$ component are summarized in Table S5. In addition, the three parameters of total $\Delta A$, relative $\Delta A$, and effective EET efficiency were estimated for individual chlorosomes (see Supplementary Note 7). We then examined how these parameters were associated with each $\tau$ component[16]. Specifically, 2D histograms were constructed with $\tau$ on the horizontal axis and one of total $\Delta A$, relative $\Delta A$, or effective EET efficiency on the vertical axis (Figs. S20–S22), using bin sizes of 0.25 ps for $\tau$, $5 \times 10^{-6}$ for total $\Delta A$, 0.03 for relative $\Delta A$, and 0.15 for effective EET efficiency. These 2D histograms were then fitted using the following function, which is a sum of 2D Gaussian distributions corresponding to each $\tau$ component:

$$\mathrm{f}(\tau, s) = \sum_{i=1}^{N} A_{2\mathrm{D},i} \exp\left(-\frac{1}{2}\left(\left(\frac{\tau - \tau_{0i}}{\tau_{\mathrm{w}i}}\right)^2 + \left(\frac{s - s_{0i}}{s_{\mathrm{w}i}}\right)^2\right)\right), \quad (4)$$

where $N$ is the number of $\tau$ components, and $A_{2\mathrm{D},i}$ is the amplitude of the 2D Gaussian associated with component $i$. The parameters $\tau_{0i}$ and $\tau_{\mathrm{w}i}$ denote the center position and $1/\sqrt{\mathrm{e}}$ half-width, respectively, along the horizontal $\tau$-axis, whereas $s_{0i}$ and $s_{\mathrm{w}i}$ denote the corresponding quantities along the vertical $s$-axis. The FWHM is obtained by multiplying the $1/\sqrt{\mathrm{e}}$ half-width by the factor $2\sqrt{2\,ln2}$.

In the 2D distribution fitting, the center positions $\tau_{0i}$ and widths $\tau_{\mathrm{w}i}$ for each $\tau$ component were fixed at the values estimated from the 1D distribution analysis as listed in Table S5. The population fraction $P_i$ of each $\tau$ component was also constrained to match the values obtained from the 1D distribution analysis (Table S5). From this constraint, the amplitude $A_{2\mathrm{D},i}$ can be expressed in terms of $P_i$, $\tau_{\mathrm{w}i}$, and $s_{\mathrm{w}i}$ as follows. First, the integral $I_i$ of the 2D Gaussian for component $i$ is

$$I_i = 2\pi\, A_{2D,i}\, \tau_{wi}\, s_{wi}\,. \tag{5}$$

Because $I_i$ corresponds to the volume of each 2D Gaussian, $I_{all} = \sum_i^N I_i$ gives the total volume over all $\tau$ components. Accordingly, the volume fraction $I_i/I_{all}$ equals the population fraction $P_i$, i.e., $P_i = I_i/I_{all}$. Substituting this relation into equation (5) gives

$$A_{2D,i} = \frac{P_i}{\tau_{wi}\, s_{wi}} \frac{I_{all}}{2\pi}\,. \tag{6}$$

By substituting equation (6) into equation (4) and replacing the factor $I_{all}/2\pi$, which is independent of component $i$, with a constant $A_0$, the final fitting function was derived as follows:

$$f(\tau, s) = A_0 \cdot \sum_{i=1}^{N} \frac{P_i}{\tau_{wi}\, s_{wi}} \exp\left(-\frac{1}{2}\left(\left(\frac{\tau - \tau_{0i}}{\tau_{wi}}\right)^2 + \left(\frac{s - s_{0i}}{s_{wi}}\right)^2\right)\right). \tag{7}$$

In this fitting function, $A_0$ is simply a scaling factor, and all other parameters except $s_{0i}$ and $s_{wi}$ were fixed at the values estimated from the 1D distribution analysis as listed in Table S5. During fitting, the $1/\sqrt{e}$ full width along the $s$-axis, given by $2s_{wi}$, was constrained to be no smaller than the corresponding histogram bin size. Because the 1D distribution analysis distinguished five $\tau$ components for each sample, $N$ was set to 5.

In practice, the fitting results obtained using equation (7) depended on the initial values chosen for $s_{0i}$. Therefore, to determine optimal initial values, we performed a Monte Carlo procedure in which the fitting was repeated with randomly selected initial values of $s_{0i}$ (see Fig. S27 for the case in which the $s$-axis represents total $\Delta A$). Specifically, initial values of $s_{0i}$ were randomly chosen within the range of the $s$-axis, and the fitting was then performed. Three representative fitting results obtained with different initial values, together with their corresponding RSS values, are shown in Fig. S27a, illustrating the dependence of the fitting results on the initial values. Repeating this procedure at least 20,000 times produced a distribution of RSS values (Fig. S27b), where smaller RSS values indicate better fits. Thus, we retained only the best-fitting results, defined as those with RSS values in the lowest 1% of this distribution, to construct the histograms of the estimated $s_{0i}$ values (Fig. S27c). The resulting histograms were fitted with 1D Gaussian functions to estimate their centers. The centers estimated for all $\tau$ components were used as the initial values for the final fitting, yielding the optimized results. For each 2D distribution, the $s$-axis parameter was selected from total $\Delta A$, relative $\Delta A$, and effective EET efficiency, and this Monte Carlo-based procedure was applied to estimate the distribution of the $s$-axis parameter associated with each $\tau$ component. All results are shown in Figs. S2–S4.

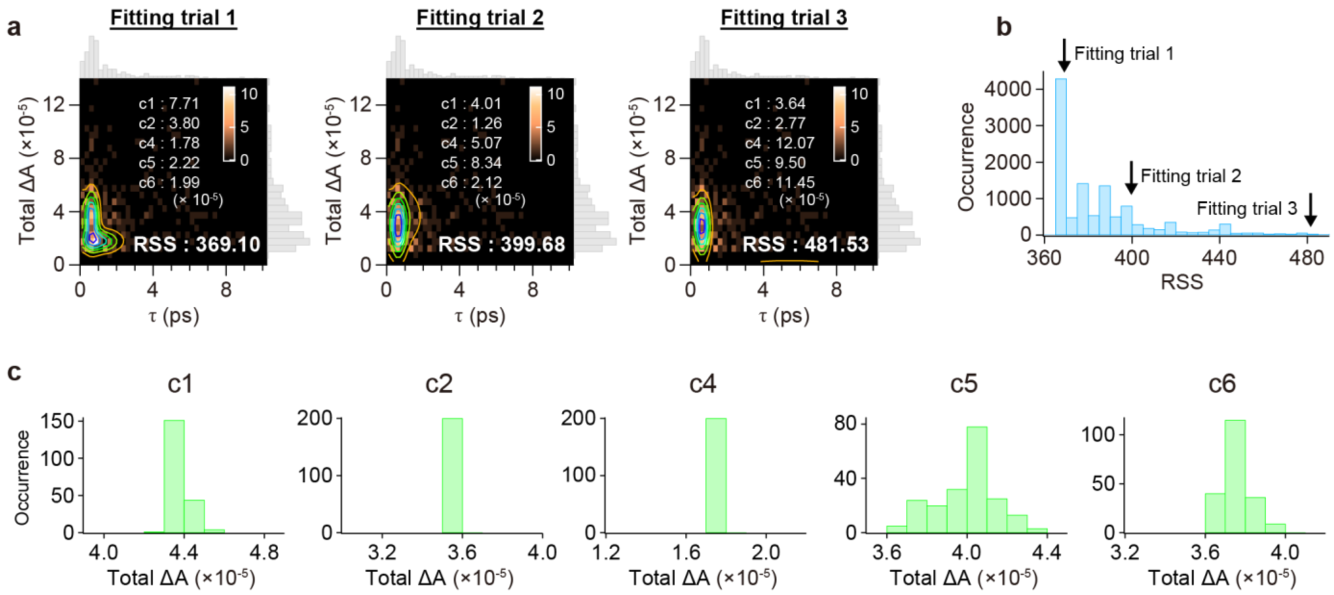

**Fig. S27 | 2D distribution analysis using five-component 2D Gaussian fitting with Monte Carlo sampling.**

(**a**) Three representative results obtained from five-component 2D Gaussian fits applied to the 2D distribution of total $\Delta A$ versus $\tau$, using randomly sampled initial total $\Delta A$ values for each $\tau$ component. The 2D distribution was obtained for *tep*_WT chlorosomes measured using the 800-nm probe. For this data set, five $\tau$ components, excluding c3, were identified (see the main text). Fitting results obtained from different initial values are shown as contours in each panel. The initial total $\Delta A$ values for each $\tau$ component and the estimated RSS value are shown in each panel. The projections of the 2D distribution onto the *x*- and *y*-axes are shown as gray histograms at the top and right of each panel, respectively. (**b**) Distribution of RSS values obtained from 20,000 trials. The RSS values for trials 1–3 in (a) are indicated by arrows. (**c**) Distributions of the fitted total $\Delta A$ values for each $\tau$ component, constructed from the trials corresponding to the lowest 1% of RSS values among the 20,000 trials.

Estimation of *p*-values for pairwise comparisons of *s* distributions

As described above, for each $\tau$ component, we obtained the distribution of the *s*-axis parameter characterized by its center position $s_{0i}$ and width $s_{wi}$. To assess whether the center positions of the *s* distributions differed significantly between components, Welch's *t*-test was applied. The *t*-statistic, which quantifies the difference between the center positions of components *i* and *j*, was calculated as

$$t = \frac{s_{0i} - s_{0j}}{\sqrt{\frac{u_i^2}{n_i} + \frac{u_j^2}{n_j}}}, \quad (8)$$

where $s_{0i}$ ($s_{0j}$) is the center position along the *s*-axis for component *i* (*j*), obtained from the 2D distribution fitting. $n_i$ ($n_j$) denotes the effective sample size for the distribution of component *i* (*j*), estimated by multiplying the total number of data points used to construct the 2D distribution by the population fraction $P_i$ ($P_j$). $u_i^2$ ($u_j^2$) is the unbiased variance for component *i* (*j*), which is related to the $1/\sqrt{e}$ half-width $s_{wi}$ ($s_{wj}$) obtained from the 2D distribution fitting, as follows:

$$u_i^2 = s_{wi}^2 \frac{n_i}{n_i - 1} \ . \tag{9}$$

From the obtained $u_i^2$ ($u_j^2$) and $n_i$ ($n_j$), the degrees of freedom (DOF) were evaluated as

$$\mathrm{DOF} = \frac{\left(\frac{u_i^2}{n_i} + \frac{u_j^2}{n_j}\right)^2}{\frac{\left(\frac{u_i^2}{n_i}\right)^2}{n_i - 1} + \frac{\left(\frac{u_j^2}{n_j}\right)^2}{n_j - 1}} \ . \tag{10}$$

Using the *t*-statistic and the DOF, we estimated *p*-values for the differences between the center positions of the *s* distributions of components *i* and *j*. Small *p*-values indicate significant differences between these center positions. Statistically significant differences between these center positions are indicated by asterisks in Figs. 3a–d, S2–S4, and S13b, d.

Robustness of the 2D distribution analysis to histogram binning conditions

As described in Supplementary Note 9, the robustness of the $\tau$-distribution analysis using histograms constructed with a bin size of 0.25 ps was confirmed. Based on these results, the 2D distribution analysis was performed using the 2D histograms constructed with a $\tau$-axis bin size of 0.25 ps, together with *s*-axis bin sizes of $5 \times 10^{-6}$ for total $\Delta A$, 0.03 for relative $\Delta A$, and 0.15 for effective EET efficiency (Figs. S20–S22). Because the 2D distribution profile can also vary with these *s*-axis bin sizes, we assessed the robustness of the 2D fitting results to binning conditions by varying each *s*-axis bin size by ±20%.

This robustness analysis was performed for *tep*_WT chlorosomes. Decreasing each *s*-axis bin size by 20% yielded 2D fitting results nearly identical to those shown in Figs. S20–S22 (Fig. S28). Likewise, increasing each *s*-axis bin size by 20% yielded largely unchanged results. For the three parameters (total $\Delta A$, relative $\Delta A$, and effective EET efficiency), the means and standard deviations of the fitted center positions and FWHMs across the tested binning conditions were calculated for each $\tau$ component (Table S8). These standard deviations were generally smaller than the corresponding bin sizes, suggesting that the variation arising from the binning conditions was negligible. These results indicate that the 2D distribution analysis was robust with respect to histogram bin size.

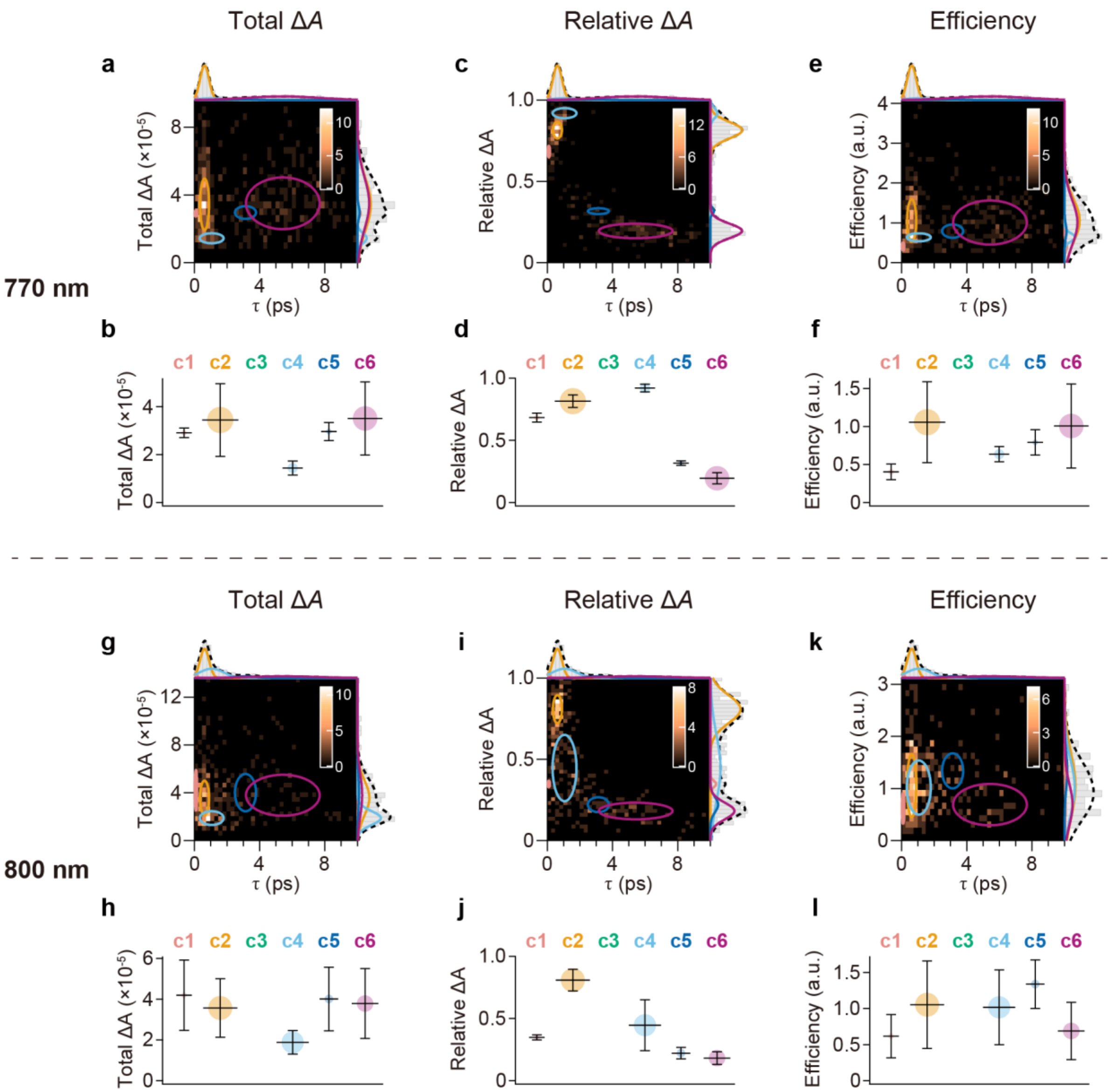

**Fig. S28 | 2D distribution analysis with bin sizes for total Δ*A*, relative Δ*A*, and effective EET efficiency decreased by 20%.**

(**a**, **c**, **e**) 2D distributions of total Δ*A* (a), relative Δ*A* (c), and effective EET efficiency (e) versus $\tau$ for *tep*_WT chlorosomes measured using the 770-nm probe, constructed with a bin size of 0.25 ps for $\tau$, together with bin sizes for total Δ*A*, relative Δ*A*, and effective EET efficiency that were decreased by 20% relative to those used in Figs. S20a, S21a, and S22a, respectively. (**b**, **d**, **f**) Estimated distributions of total Δ*A* (b), relative Δ*A* (d), and effective EET efficiency (f), obtained from five-component 2D Gaussian fits to the 2D distributions. (**g**–**l**) Corresponding results obtained using the 800-nm probe. The plots are presented in the same format as in Figs. S20 and S2.

**Table S8**

Center position and FWHM of total $\Delta A$, relative $\Delta A$, and effective EET efficiency associated with each $\tau$ component, estimated from five-component 2D Gaussian fits to the corresponding 2D distributions constructed with three distinct binning conditions (−20%, 0%, and +20%) for the respective parameter axes. The 2D distributions were obtained for *tep*_WT chlorosomes using probes at 770 nm and 800 nm. The values are shown as means ± standard deviations across the three binning conditions. For effective EET efficiency, standard deviations smaller than $0.1 \times 10^{-1}$ are indicated as <0.1.

| | Total $\Delta A$ ($\times 10^{-6}$) | | | | Relative $\Delta A$ ($\times 10^{-2}$) | | | | Efficiency ($\times 10^{-1}$) | | | |
|---|---|---|---|---|---|---|---|---|---|---|---|---|
| | 770 nm | | 800 nm | | 770 nm | | 800 nm | | 770 nm | | 800 nm | |
| | Center position | FWHM | Center position | FWHM | Center position | FWHM | Center position | FWHM | Center position | FWHM | Center position | FWHM |
| c1 | 27.3 ± 1.6 | 8.8 ± 3.9 | 41.6 ± 2.0 | 38.2 ± 2.2 | 68.7 ± 0.5 | 9.4 ± 0.9 | 34.8 ± 0.1 | 4.8 ± 0.5 | 4.5 ± 0.7 | 3.0 ± 1.6 | 6.0 ± 0.4 | 6.1 ± 1.1 |
| c2 | 34.4 ± 0.5 | 35.8 ± 1.2 | 35.7 ± 0.1 | 32.3 ± 1.3 | 81.3 ± 0.3 | 12.1 ± 0.4 | 80.9 ± 0.1 | 20.1 ± 0.2 | 10.5 ± <0.1 | 12.8 ± 0.4 | 10.5 ± <0.1 | 14.3 ± 0.2 |
| c3 | - | - | - | - | - | - | - | - | - | - | - | - |
| c4 | 15.2 ± 1.0 | 8.3 ± 1.8 | 18.2 ± 0.6 | 13.7 ± 0.5 | 92.1 ± 0.4 | 7.6 ± 0.3 | 45.1 ± 0.4 | 47.9 ± 0.2 | 6.6 ± 0.2 | 2.5 ± 0.5 | 9.9 ± 0.2 | 12.0 ± 0.2 |
| c5 | 29.1 ± 0.5 | 8.5 ± 1.3 | 39.6 ± 1.1 | 36.8 ± 2.8 | 32.1 ± 0.4 | 5.3 ± 1.3 | 22.2 ± 0.1 | 11.1 ± 0.5 | 7.7 ± 0.7 | 3.1 ± 1.2 | 13.6 ± 0.3 | 7.3 ± 0.7 |
| c6 | 35.0 ± 0.4 | 36.1 ± 1.5 | 38.0 ± 0.9 | 39.8 ± 2.6 | 19.4 ± 0.2 | 11.1 ± 0.5 | 18.3 ± 0.3 | 11.7 ± 1.3 | 10.0 ± <0.1 | 13.1 ± 0.2 | 6.9 ± <0.1 | 8.3 ± 0.9 |

#### Supplementary Note 11: Assignment of the >10-ps component

The population of components with $\tau > 10$ ps varied with probe wavelength and mutation. With the 770-nm probe, the population was 9.0% for *tep*_WT, 9.5% for *tep_bchQ*, 11.9% for *lim_bciD*, and 17.7% for *lim_bciD-limQ* (Fig. S29a, b, blue), whereas with the 800-nm probe, it decreased for all samples to 2.5%, 4.1%, 6.6%, and 8.6%, respectively (Fig. S29a, b, pink). The numbers of components with $\tau > 10$ ps are listed in Table S9.

The population difference calculated as *tep_bchQ* minus *tep*_WT, corresponding to the shorter-side-chain sample minus the longer-side-chain sample, was small (Fig. S29c). By contrast, for the samples with even longer BChl side chains, the corresponding difference calculated as *lim_bciD* minus *lim_bciD-limQ* was negative (Fig. S29d), indicating a larger fraction of the >10-ps component in *lim_bciD-limQ* than in *lim_bciD*. Together, these results suggest that elongation of the BChl side chains tends to increase the population of the >10-ps component, supporting its assignment to lamellar aggregates. Furthermore, comparison of the populations obtained with the 770-nm and 800-nm probes showed that this population was lower at 800 nm than at 770 nm in all samples (Fig. S29e, f). At 800 nm, where exciton states closer to the bottom of the energy manifold are probed, a major EET pathway to the baseplate should be preferentially observed. Therefore, the lower population at 800 nm suggests that EET to the baseplate on the >10-ps timescale is minor, although components with $\tau > 10$ ps have previously been assigned mainly to EET to the baseplate[19,20]. Accordingly, as discussed in the main text, the >10-ps component may reflect subsidiary EET pathways from lamellar aggregates to the baseplate via higher-energy exciton states (see Supplementary Note 12).

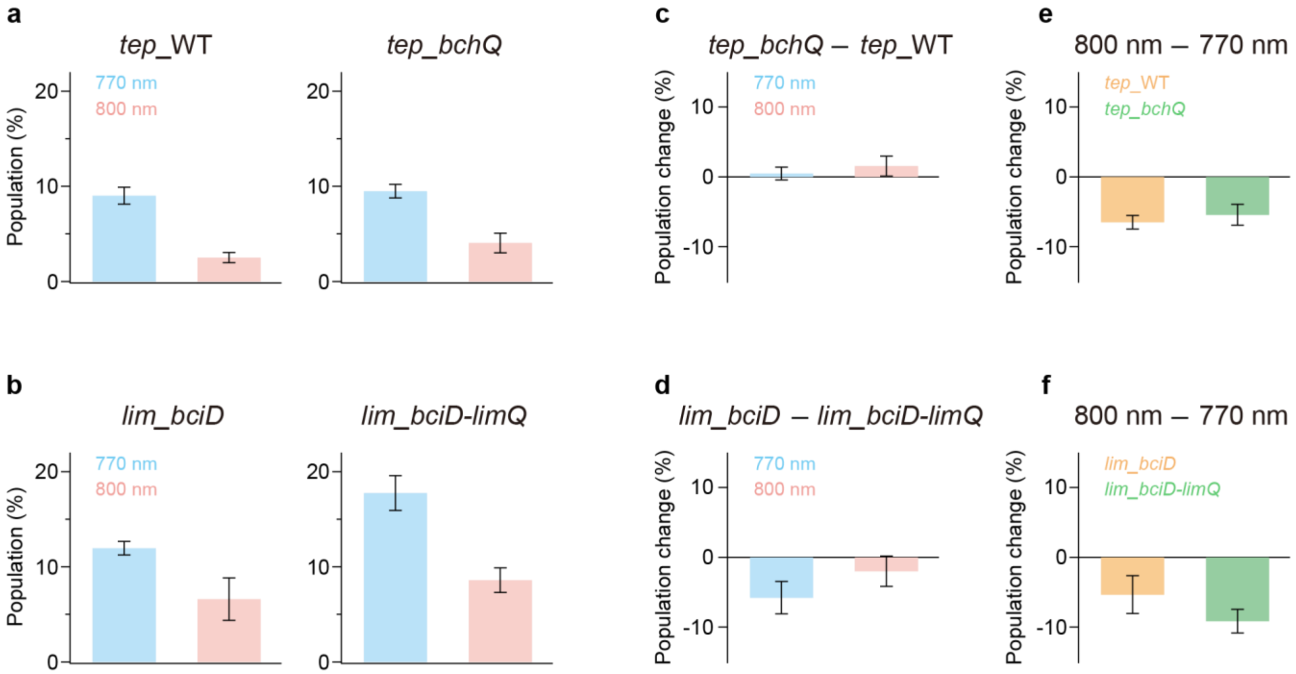

**Fig. S29 | Populations of the >10-ps components.**

(**a**, **b**) Populations of the >10-ps components in *tep*_WT and *tep_bchQ* chlorosomes (a, left and right, respectively) and those in *lim_bciD* and *lim_bciD-limQ* chlorosomes (b, left and right, respectively), obtained using probes at 770 nm (blue) and 800 nm (pink). The populations were calculated for each sample as the ratio of the number of >10-ps time constants to the total number of estimated time constants. (**c**, **d**) Mutation-induced changes in the populations of the >10-ps components, calculated by subtracting the populations in *tep*_WT from those in *tep_bchQ* (c) and those in *lim_bciD-limQ* from those in *lim_bciD* (d), at 770 nm (blue) and 800 nm (pink). (**e**, **f**) Probe-wavelength-dependent changes in the populations of the >10-ps components, calculated by subtracting the populations at 770 nm from those at 800 nm in *tep*_WT (orange) and *tep_bchQ* (green) (e), and in *lim_bciD* (orange) and *lim_bciD-limQ* (green) (f). Error bars represent standard deviations estimated from five $\tau$-distribution analyses performed on resampled data sets (see Supplementary Note 9).

**Table S9**

Number of time constants with $\tau > 10$ ps relative to the total number of estimated time constants.

| | 770 nm | 800 nm |
|---|---|---|
| *tep*_WT | 35 / 388 | 8 / 319 |
| *tep*_*bchQ* | 28 / 295 | 9 / 222 |
| *lim*_*bciD* | 38 / 318 | 7 / 106 |
| *lim*_*bciD*-*limQ* | 44 / 248 | 24 / 279 |

#### Supplementary Note 12: Light-harvesting model in chlorosomes

We propose a light-harvesting model for chlorosomes (Fig. 3e), based on our single-chlorosome TA analysis, which revealed distinct excitation characteristics of lamellar and tubular aggregates. The model can be interpreted in terms of the energy diagram shown in Fig. S30 (see also the main text). First, when lamellar aggregates absorb light, the excitation relaxes into a localized domain in 0.6 ps (c2). Subsequently, interdomain EET, likely via hopping, occurs over 5.4 ps (c6). This is proposed to be followed by EET to tubular aggregates, presumably involving lower-energy states in the outer tube. Direct EET from lamellar aggregates to the baseplate can occur on a 0.9-ps timescale (c3) but is minor in the wild type. Such direct EET may also proceed on a >10-ps timescale via higher-energy exciton states (see Supplementary Note 11).

Individual chlorosomes in which lamellar-derived EET (c6) was dominant showed longer photobleaching times (see Supplementary Note 3). In addition, the lamellar-associated components (c2 and c6) appeared to be less prone to saturation under excess excitation (see Supplementary Note 5). These results support this model, in which lamellar aggregates function as the upstream energy donors.

Upon light absorption by tubular aggregates, the initial excitation relaxes in 0.1 ps (c1), possibly accompanied by dipole reorientation and EET from the inner to the outer tube. This process may result in a more delocalized exciton than that formed in lamellar aggregates. Then, EET to the baseplate takes place in 1.1 ps (c4). Excitonic domains located away from the baseplate are likely unable to transfer energy immediately, resulting in interdomain EET over 3.1 ps (c5).

The detection of EET to and fluorescence from the baseplate even at cryogenic temperatures[21,22] implies that the 1.1-ps EET from lower-energy states in tubular aggregates to the baseplate (c4) is unlikely to be driven primarily by thermal activation. Meanwhile, in tubular aggregates, exciton states with dipole moments perpendicular to the cylindrical axis are ~100 $cm^{-1}$ higher in energy than the lowest-energy state, whose dipole moment is parallel to the axis[23]. Because this energy separation is modest, thermally activated EET to the baseplate via these higher-energy states is feasible. Given that the transition dipoles of BChl *a* in the baseplate are distributed across three-dimensional orientations[21], multiple EET pathways mediated by exciton states with different dipole orientations in tubular aggregates may facilitate efficient and robust light harvesting.

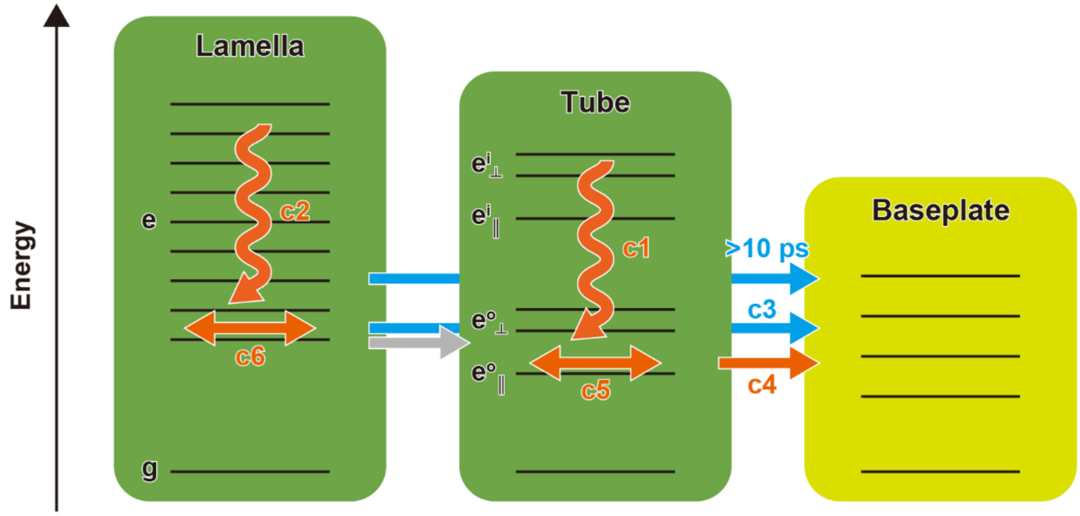

**Fig. S30 | Schematic energy diagrams of chlorosomes.**

Energy diagrams of the lamellar aggregate (left), the tubular aggregate (center), and the baseplate (right). The ground state and exciton states are denoted as g and e, respectively. The exciton states with transition dipole moments perpendicular and parallel to the cylindrical axis of the outer (inner) tubes are denoted as $e^{o}_{\perp}$ ($e^{i}_{\perp}$) and $e^{o}_{\parallel}$ ($e^{i}_{\parallel}$), respectively. The orange wavy and double-headed arrows indicate excitation relaxation (c1, c2) and interdomain EET processes (c5, c6), respectively, within the respective aggregates. The orange right-pointing arrow indicates EET from the tubular aggregates to the baseplate (c4), whereas the light-blue right-pointing arrow indicates EET from the lamellar aggregates to the baseplate (c3, >10 ps). The EET process between the lamellar and tubular aggregates, indicated by a gray arrow, was not identified in the present measurements. The assignments of the $\tau$ components c1–c6 and the >10-ps component are discussed in the main text and Supplementary Notes 11 and 12. A schematic illustration of chlorosomal light harvesting is also shown in Fig. 3e.

#### Supplementary Note 13: Light-harvesting systems balancing structural disorder and excitonic heterogeneity

In *tep_bchQ* chlorosomes, the $C8^2$ side chain of BChl *c* is shortened, promoting more uniform molecular packing and reducing structural disorder within aggregates. Consequently, not only tubular aggregates but also lamellar aggregates likely retain delocalized exciton states during excitation relaxation, as discussed in the main text. Meanwhile, the range of the relative $\Delta A$ distribution associated with tubular aggregates (c1 and c5) was greater (Fig. 3d) than the corresponding ranges in *tep*_WT chlorosomes (Fig. 3b) and other mutant chlorosomes (Fig. S3g, h). These results suggest substantial heterogeneity in the exciton states of tubular aggregates among *tep_bchQ* chlorosomes.

Reduced steric hindrance between BChl molecules may lead to denser packing, which could in turn increase structural rigidity at the cost of mechanical flexibility. If local stress exceeds a threshold, it could cause structural defects. Accordingly, while dense packing likely suppresses structural disorder and stabilizes delocalized excitons, local distortions and defects within the tubular aggregates may cause variations in excitonic domain size. Lower steric hindrance between BChl molecules promotes the formation of larger multi-tubular structures[8], which appear favorable for efficient and stable light harvesting, but could introduce trade-offs in excitonic dynamics. Ultimately, in native aggregates such as those in *tep*_WT chlorosomes, structural rigidity and flexibility are thought to be balanced by modulating packing density via steric interactions, thereby optimizing functional robustness. This may represent a unique design strategy for protein-independent antennas, which lack a protein scaffold.
